# Merle phenotypes in dogs - *SILV* SINE insertions from Mc to Mh

**DOI:** 10.1101/328690

**Authors:** Langevin Mary, Helena Synkova, Tereza Jancuskova, Sona Pekova

## Abstract

It has been recognized that the Merle coat pattern is not only a visually interesting feature, but it also exerts an important biological role, in terms of hearing and vision impairments. In 2006, the Merle (M) locus was mapped to the *SILV* gene with a SINE element in it, and the inserted retroelement was proven causative to the Merle phenotype. Mapping of the M locus was a genetic breakthrough and many breeders started implementing *SILV* SINE testing in their breeding programs. Unfortunately, the situation turned out complicated as genotypes of Merle tested individuals did not always correspond to expected phenotypes, sometimes with undesired health consequences in offspring. Two variants of *SILV* SINE, allelic to the wild type sequence, have been described so far - Mc and M.

Here we report a significantly larger portfolio of existing Merle alleles (Mc, Mc+, Ma, Ma+, M, Mh) in Merle dogs, which are associated with unique coat color features and stratified health impairment risk. The refinement of allelic identification was made possible by systematic, detailed observation of Merle phenotypes in a cohort of 181 dogs from known Merle breeds, by many breeders worldwide, and the use of advanced molecular technology enabling the discrimination of individual Merle alleles with significantly higher precision than previously available.

We also show that mosaicism of Merle alleles is an unexpectedly frequent phenomenon, which was identified in 30 out of 181 (16.6%). dogs in our study group. Importantly, not only major alleles, but also minor Merle alleles can be inherited by the offspring. Thus, mosaic findings cannot be neglected and must be reported to the breeder in their whole extent.

In light of negative health consequences that may be attributed to certain Merle breeding strategies, we strongly advocate implementation of the refined Merle allele testing for all dogs of Merle breeds to help the breeders in selection of suitable mating partners and production of healthy offspring.

## INTRODUCTION

In dogs, coat color is a polymorphic and quite complicated issue. Since Little ([1]1957) had described more than 20 loci affecting coat colors according to dog phenotypes, only a few genes have been recognized as being involved in the pigmentation – the most prominent being locus E (MC1R gene, [2], [3], [4]), locus K (CBD103 gene, [5]), locus A (ASIP gene, [6], [7], [8]), locus B (TYRP1 gene, [9]), locus D (MLPH gene, [10]), locus S (MITF gene, [11]) and M locus (*SILV* gene, [12]).

The Merle (M) locus was suggested by Little ([1]) as being responsible for Merle pattern, which is coat color where eumelanic regions are incompletely and irregularly diluted resulting in typical intensely pigmented patches. In 2006, the gene corresponding to the dominant allele of the M locus was finally recognized by Clark et al. [12]). Nevertheless, previous attempts were focused on factors, which are secreted primarily by keratinocytes (cells surrounding melanocytes) to stimulate the switch between phaeomelanin and eumelanin production ([13]). However, searching for the gene candidates responsible for Merle phenotypes excluded the genes involved in the melanogenic pathway, i.e., *micropthalmia transcription factor* (*MITF*), *tyrosinase* (*TYR*), *tyrosine related protein* (*TYRP1*), or *dopachrome tautomerase* (*DCT*) ([14]).

The Merle pattern can be seen in various breeds, such as the Australian Shepherd Dog, Australian Koolie, Border Collie, Dachshund, French Bulldog, Louisiana Catahoula, Labradoodle, Miniature American Shepherd, Miniature Australian Shepherd, Pyrenean Shepherd, Rough Collie, Shetland Sheepdog, Welsh Sheepdog, Cardigan Welsh Corgi, Chihuahua, Great Dane etc. Merle is thought to be inherited in an autosomal, incomplete dominant way. Dogs heterozygous for the M allele show a typical coat pattern, however, dogs homozygous for the M allele may also exhibit auditory and ophthalmologic impairments and abnormalities together with very pale or completely white phenotypes (Strain et al. 2009). Such negative health effects associated with the M locus encouraged the research to identify a particular gene responsible. Clark et al. ([12]) confirmed the *SILV* gene as being the cause of the Merle pattern - the short interspersed element (SINE) is inserted in the structure of the gene. Clark et al. ([12]) has mapped the SINE insertion of approximately 253 bp at the intron 10 /exon 11 border of the *SILV* gene. A typical SINE element is composed of a body and a poly-A tail of variable length. It has been assumed that it might be the extent of the poly-A stretch, which plays the peculiar biological role, visually recognized as different qualities of the Merle coat pattern. It has been suggested that the poly-A tail, being a monotonous genomic structure, is prone to replication errors, caused by a slippage of the cellular replication machinery in such a challenging genomic context. It has also been observed that SINE inserts of different lengths do exist. Shorter SINE was ascribed to the allele Mc (Cryptic Merle) which has no apparent effect on the dogs’ phenotype, while longer SINE inserts were found to be responsible for the individual Merle phenotypes ([12], [15]).

This discovery enabled commercial testing of the Merle gene by various laboratories in different breeds worldwide. However, the occurrence and combinations of the above alleles could not explain various irregularities in dog phenotypes and unexpected results of breeding. In 2011, the Catahoula Club EU in the Czech Republic started testing the Merle gene within the Czech population of the Louisiana Catahoula (LC) breed in cooperation with Biofocus laboratory (Germany) ([16]). LC is the North American breed famous for its Merle (leopard) pattern, where, initially, no regulations for breeding of Merle to Merle individuals had been applied in the breeding strategies, contrary to other Merle carrying breeds, especially in Europe. Therefore, many dogs carrying the Merle gene can be expected within the breeding pool. The pilot testing by Biofocus proved that many unusual solid-color in appearance Catahoulas carry Merle alleles, but of a different SINE size than those found by Clark et al. ([12]). The middle-sized SINE allele has been named “Atypical Merle” (Ma) in order to distinguish it from those already known ([16]). Further commercial testing, which started in 2015 and continues until now, has involved a wider LC population and later also other breeds and confirmed the Ma existence and its role in various specific and distinguishable Merle patterns ([17]).

Originally, only two alleles had been recognized by Clark et al. ([12], [15]). However, unusual or unpredictable results of breedings, which cause unexplainable phenotypes of offspring, have confirmed the need for further research in this field. The length of SINE seems to be a key factor for the explanation of different effects on the final coat pattern. However, various commercial laboratories use different methods and instruments for SINE length detection. This can strongly influence final results, their accuracy, and ability to distinguish different alleles. Some laboratories have already acknowledged the existence of the Ma allele and have started reporting it. Other laboratories are still waiting for scientific confirmation, because a relevant study on this subject has not been published as of yet. There is also only limited and/or biased choice of dogs and breeds available for testing in particular studies ([18], [19]). Thus, it might be difficult to estimate the rate of occurrence for different Merle alleles in a given breed without a thorough population breed study. It is usually beyond the ability of any commercial laboratory, especially without a close cooperation with dog breeders of particular breeds. Solely collecting data from the Merle testing is not sufficient to properly evaluate a relationship between genotypes and phenotypes without knowledge of particular dogs and their relationships.

Thus, the aim of our study was to evaluate the phenotype/genotype correlations between the individual Merle coat patterns and specific lengths of SINE in the *SILV* gene, i.e. to address genetically the extent of Merle alleles we have anticipated phenotypically in Lousiana Catahoula dogs, as well as in other Merle breeds (Australian Koolie, Australian Shepherd, Border Collie, French Bulldog and others).

Herein we confirm that in many Merle breeds, there exists a significantly higher number of Merle alleles than previously suggested. The individual Merle alleles can be distinguished according to their specific SINE lengths, corresponding unique Merle phenotypes and breeding results. To better understand the complexity of coat color genetics in Merle breeds, the *SILV* SINE gene genotyping has been complemented by a comprehensive Next Generation Sequencing genotyping of all known major coat color loci and their modifiers – a test called SuperColorLocus, which we have developed.

## MATERIALS AND METHODS

### Animals and sample collection

Biological material (buccal swab, hair, blood, sperm cells) was collected from 181 dogs (*Canis lupus familiaris*) of 14 breeds: Australian Shepherd Dog (ASD, 40), Australian Koolie (AK, 23), Border Collie (BC, 18), Dachshund (D, 9), French Bulldog (FB, 2), Louisiana Catahoula (LC, 73), Labradoodle (LD, 1), Miniature American Shepherd (MAS, 2), Miniature Australian Shepherd (MAUS, 2), Pyrenean Shepherd (PS, 1), Rough Collie (RC, 3), Shetland Sheepdog (SSD, 2), Welsh Sheepdog (WS, 3), and Mudi (M, 2).

The animals in the study were selected both for their visually detectable Merle phenotypes, and/or pedigree relationships with Merle-expressing individuals or individuals anticipated to carry the Merle allele. Dogs related by pedigree are listed in the Supplementary material (Table S1).

No dogs were housed for research purposes; all dogs were privately owned pets. The sampling of the biological material (buccal swab, hair, blood, sperm cells) was performed by veterinary practitioners or owners under written recommendations by the laboratory. All samplings were performed as a part of a routine commercial laboratory investigations for other routine laboratory genetical testing relevant for the breeding programs of the individual breeds recommended by the individual breeding clubs. All samples were non-invasive, causing no harm or pain to the animals and were contributed by the owners voluntarily as a part of a routine commercial laboratory testing for other purposes. No ethical commitee evaluation was applicable for this type of research. All owners have signed a written informed consent for the research, publication of results and pictures of their dogs.

### Isolation of DNA from biological samples

From buccal swab and sperm cells, DNA was isolated using QIAamp DNA Mini Kit (Qiagen, DE) according to the instruction of the manufacturer. For DNA isolation from hair, incubation in Lysis buffer was performed for 12 hours prior to proceeding with the isolation procedure.

### *SILV* SINE genotyping

A fragment of canine *SILV* gene with the potential SINE insertion ([12]) was PCR amplified using FAM-labelled forward primer (5′atcaaggtcatgtaaacccaggcc 3′) and reverse primer (5′ gtagccagtgggtgctaccacgtg 3′) with QIAGEN Multiplex PCR kit (Qiagen, DE), according to the instructions of the manufacturer. The PCR profile was the following: initial denaturation 95°C for 15 min, with 40 cycles 95°C/30 sec, 56°C/30 sec, 72°C/45 sec. PCR fragments were separated on ABI3500 genetic analyzer (ThermoFisher Scientific, USA) using fragment analysis module and the GeneMapper software, v5.0 plug-in.

### SuperColorLocus testing

SuperColorLocus (SCL) multiplex PCR product was prepared using QIAGEN Multiplex PCR kit (Qiagen, DE) and 200 nM each of forward and reverse primers to amplify the following genomic loci: Locus E - MC1R c.233G>T, MC1R c.250G>A, MC1R c.916C>T, MC1R c.790A>G; Locus K - CBD103 c.67_69del; Locus A - ASIP c.246G>T, ASIP c.250G>A, ASIP c.288C>T; Locus B - TYRP1 c.121T>A, TYRP1 c. 991C>T, TYRP1 c.1033_1036del; Locus D - MLPH c.-22G>A, PSMB7 c. 146T>G. Locus S (MITFinsSINE) and ASIPinsSINE were analyzed using fragment analysis analogically to *SILV* SINE testing. Genotyping primers are available upon request.

To build NGS libraries, the SCL multiplex PCR product was end-trimmed, adapters-ligated and barcoded using NEBNext^®^ Fast DNA Library Prep Set for Ion Torrent™ (NEB, USA) according to the instructions of the manufacturer. Built libraries were quantified using Ion Univ Library Quant Kit (ThermoFisher Scientific, USA) and subjected to emPCR overnight, using OT2 semi-automatic platform (LifeTechnologies). After bead enrichment (OT2 platform), NGS sequencing chip was loaded (typically v 314 or 316) and sequenced (PGM, LifeTechnologies). Raw data has been analyzed using in-house, ISO 17025 - validated bioinformatic software for SNP genotyping.

### Parentage testing

STR parentage analysis was performed using Canine ISAG STR Parentage Kit (ThermoFisher Scientific, USA) according to the instructions of the manufacturer. The system has been validated for forensic use in dog paternity testing and evaluates 21 ISAG-recommended polymorphic STR markers.

### Statistical data analysis

Data was obtained from 14 breeds, with various numbers of the individuals within each breed. Among the more frequent breeds, LC (73), ASD (40), AK (23), BC (18), and D (9) were used for a detailed analysis, while data from less frequent breeds, such as RC (3), WS (3), FB (2), M (2), MAS (2), MAUS (2), SSD (2), LD (1), PS (1), may have been used in overall calculations only (if not stated otherwise). Prevalence comparisons across all breeds may not have been performed in all cases because of the small subject numbers in some breeds. Therefore, we show here the qualitative differences found within the more populated breeds incuded in the study.

## RESULTS

### Refined *SILV* SINE genotyping

*SILV* SINE fragments were separated in denaturing polyacrylamide gel on ABI3500 Genetic Analyzer to obtain the highest resolution possible for fragment analysis technology. ABI3500 allowed us to discriminate individual alleles differing in 1 nucleotide in length even for long PCR products (approx 500 bp). Using this approach, we were able to confirm the presence of the already recognized Merle alleles Mc and M, as well as also identifying novel Merle alleles and have named them Mc+, Ma, Ma+ and Mh. Thus, we report here the presence of 7 individual types of Merle alleles and binned them as follows: The wild type allele bin was assigned a length of 171 bp, the bins for the other *SILV* SINE alleles are: Mc (200–230 bp), Mc+ (231–246 bp), Ma (247–254 bp), Ma+ (255–264 bp), M (265–268 bp), and Mh (269–280 bp).

**Fig 1.**
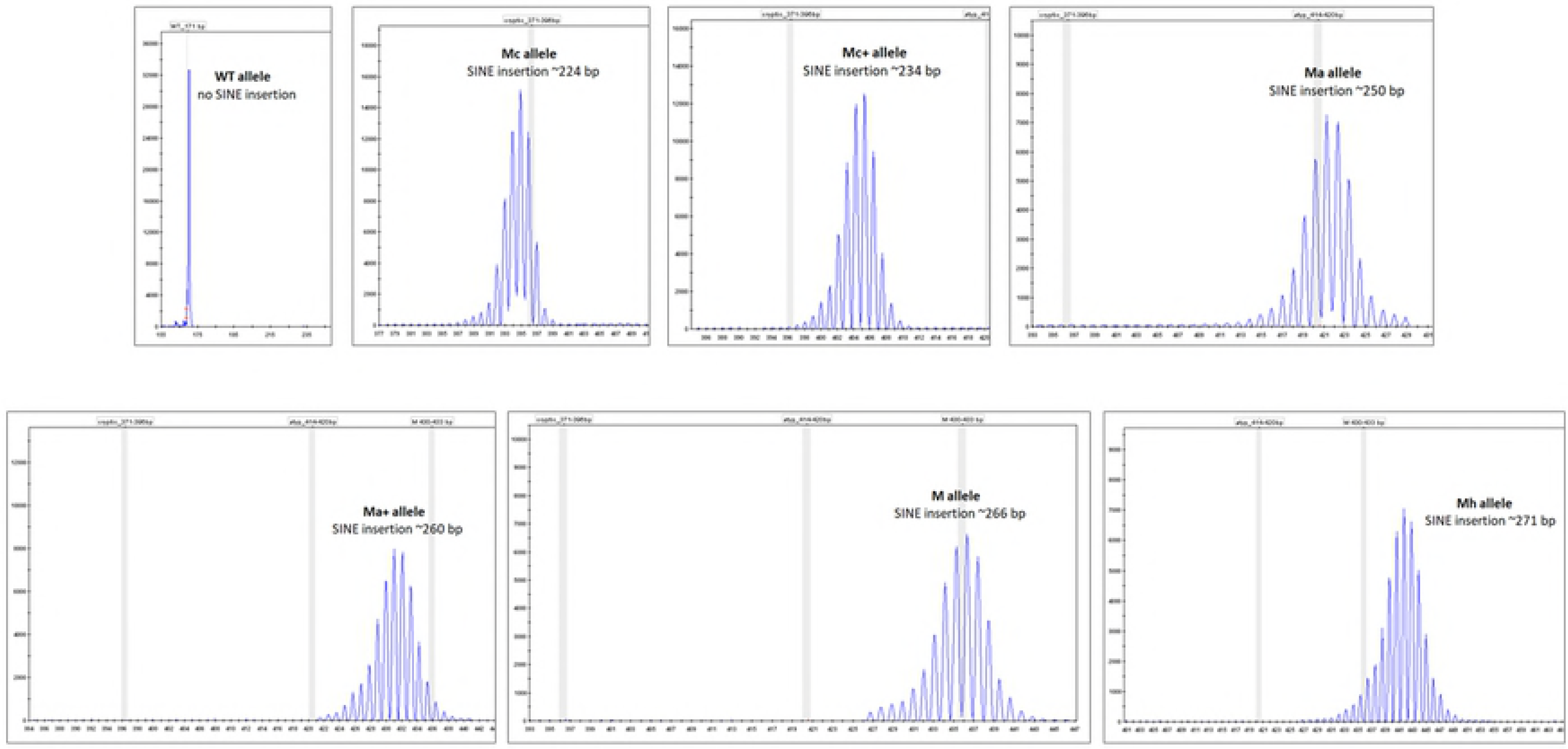
shows typical chromatograms of the individuall alleles as obtained using ABI3500. The whole porfolio of seven Merle alleles was found in three breeds with a larger number of dogs represented (LC, ASD, BC) (*n* ≥ 9, Table 1). Nevertheless, the length of particular Merle alleles did not differ significantly between all breeds tested.

**Table 1.** shows the typical size range (in bp) for the Merle alleles as identified in the individual breeds in the study. The typical size range (in bp) for the Merle alleles. ASD – Australian Shepherd Dog, AK – Australian Koolie, BC – Border Collie, D – Dachshund, FB – French Bulldog, LC – Louisiana Catahoula, LD – Labradoodle, MAS – Miniature American Shepherd, MAUS – Miniature Australian Shepherd, PS – Pyrenean Shepherd, RC – Rough Collie, SSD – Shetland Sheepdog, WS – Welsh Sheepdog, *n* – number of tested dogs. The size of the wild type allele is 171 bp.

| ALLELE | MAJOR PHENOTYPIC FEATURES |
| --- | --- |
| m | no Merle pattern - solid coat |
| Mc | no Merle pattern - solid coat |
| Mc+ | no Merle pattern - solid coat |
| Ma | no Merle pattern - diluted - brownish hue |
| Ma+ | muted, undefined, diluted - brownish hue |
| M | classic Merle |
| Mh | minimal Merle, areas deleted to white, tweed |

### Precision of the allelic size measurement

To assess the precision of allelic size measurement using ABI3500 fragment analysis, the length of paternal and maternal Merle alleles during their passage to the offspring generation were evaluated.

Figs 2A-2C show the size and inheritance of the obligatory parental Merle alleles across one generation in Dachshund (2A), Australian Shepherd (2B) and Catahoula (2C). The data shows a high degree of accuracy of the allelic size measurement and also preservation of the length of the poly-A tail between generations.

**Fig 2A.**
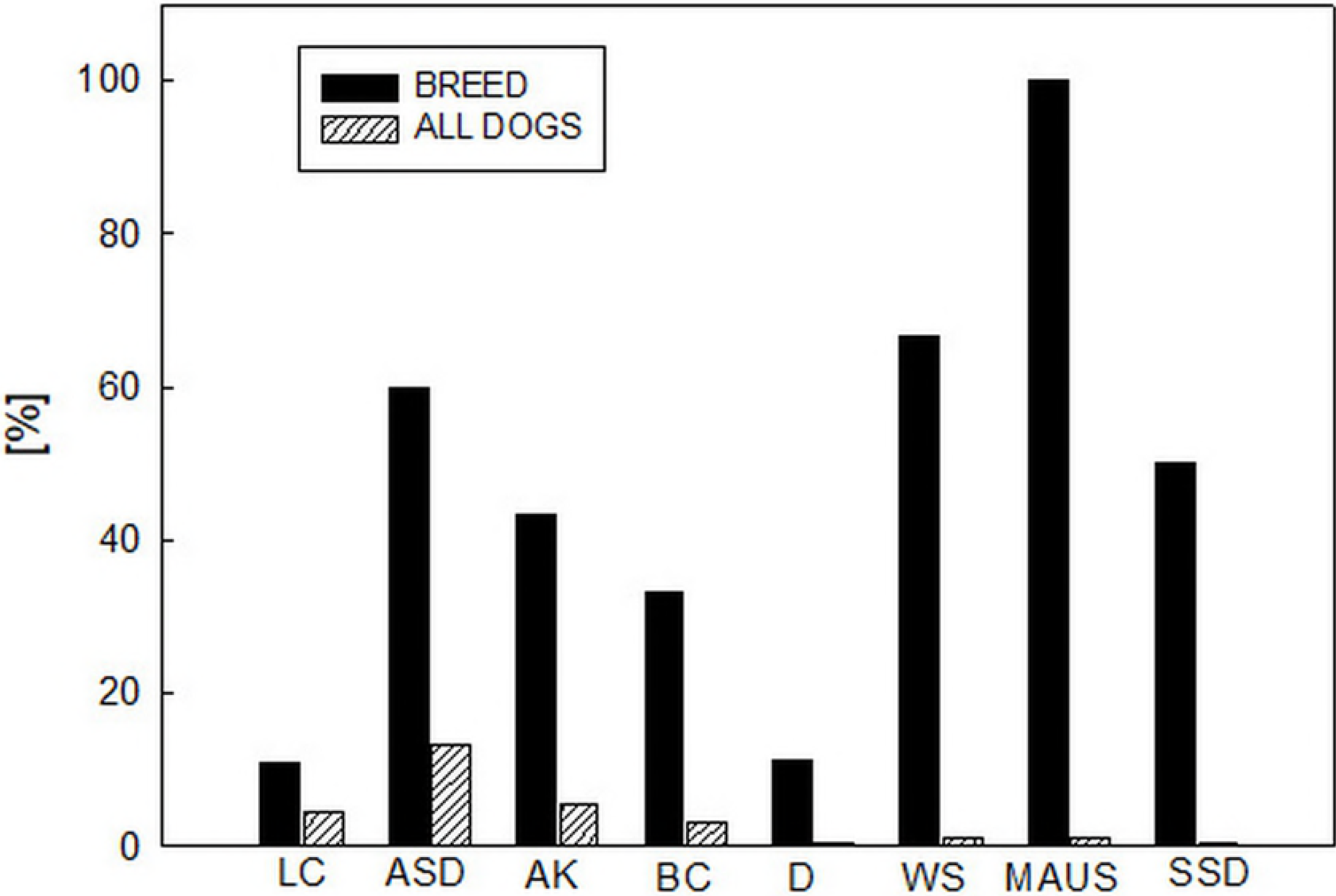

**Fig 2B.**
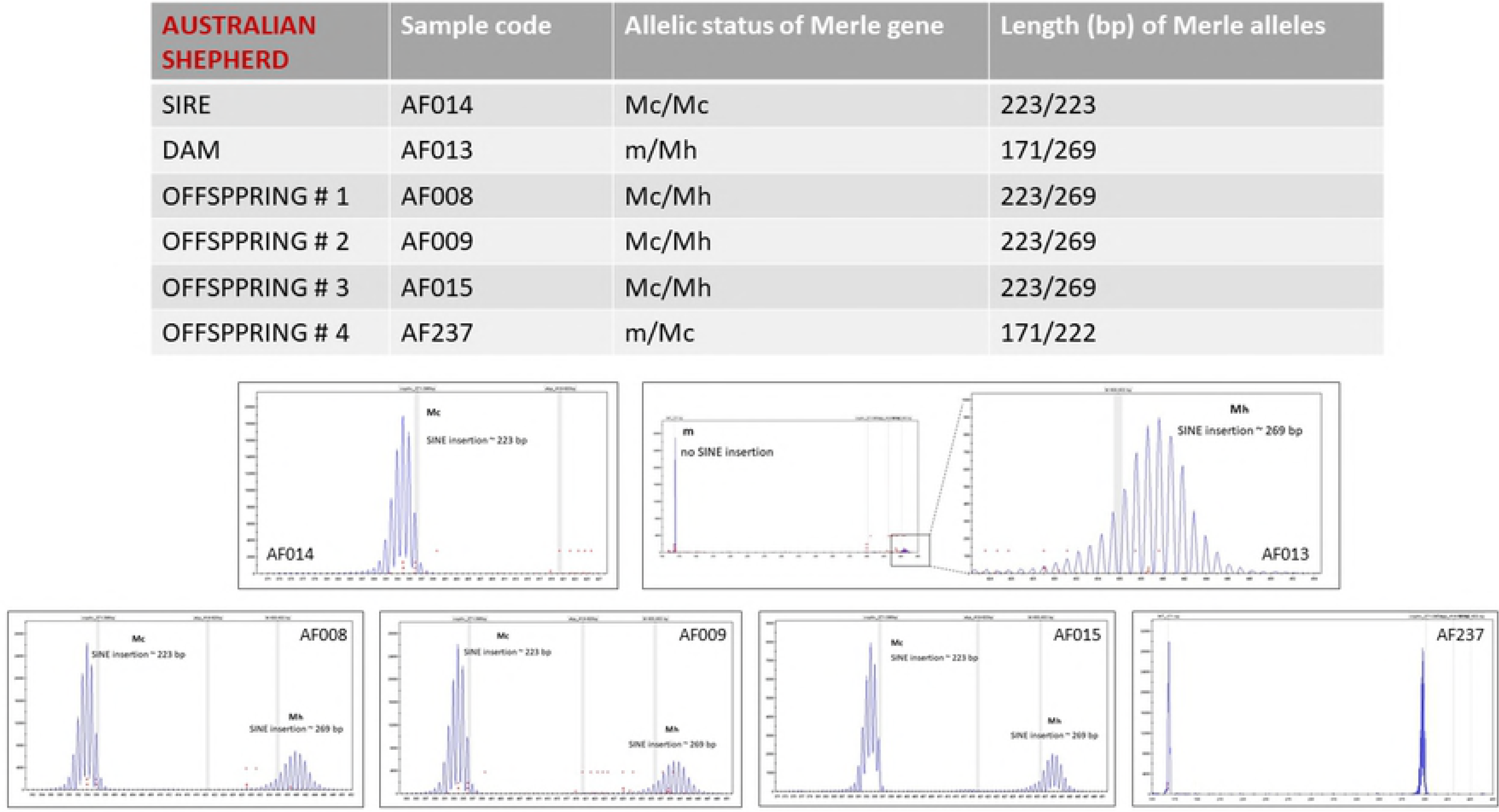

**Fig 2C.**
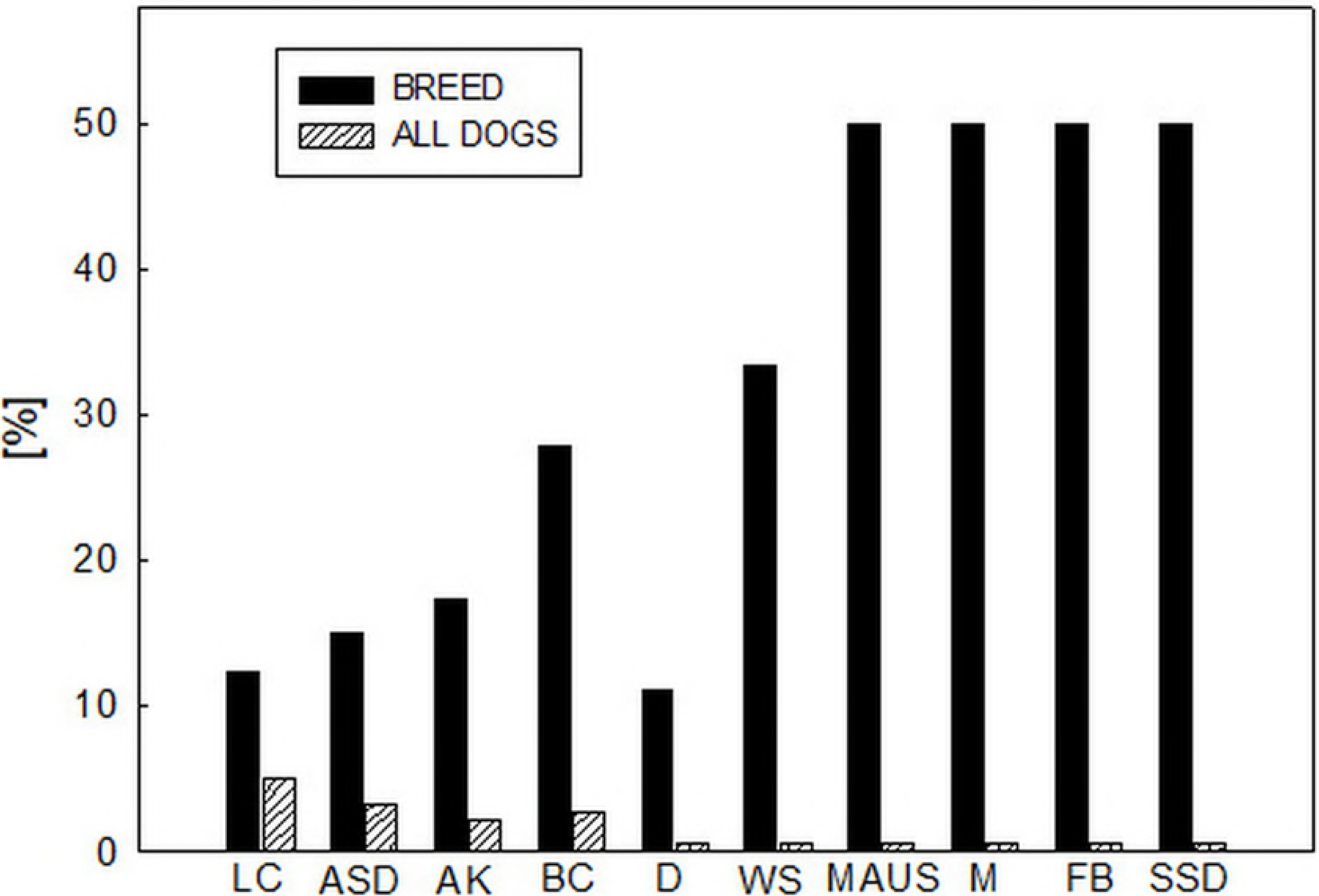

### Novel Merle alleles – phenotype/genotype correlations

As well as the already recognized m, Mc and M alleles, we have identified four novel Merle alleles and termed them Mc+, Ma, Ma+ and Mh. These alleles occur in classic heterozygous, but also in compound heterozygous combinations. All of them, classic heterozygous or compound heterozygous genotypes are associated with a typical, visually conspicuous phenotype. A detailed description thereof is given in detail in Figure 3, which, for educational reasons, has been separated into individual subfigures 3A-3X. All 7 Merle alleles, in theory, can make 28 individual allelic combinations (m/m, m/Mc, Mc/Mc, Mc/Mc+, m/Mc+, Mc+/Mc+, m/Ma, Mc/Ma, Mc+/Ma, Ma/Ma, m/Ma+, Mc/Ma+, Mc+/Ma+, Ma/Ma+, Ma+/Ma+, m/M, Mc/M, Mc+/M, Ma/M, Ma+/M, M/M, m/Mh, Mc/Mh, Mc+/Mh, Ma/Mh, Ma+/Mh, M/Mh, Mh/Mh). In our study, with the exception of 4 allelic combinations Mc/Mc+, Mc+/Ma, Ma/Ma+, Ma+/Mh we have observed all of them. Figs 3A-3X give a detailed description of the individual genotypes and their associated phenotypes. In the Supplementary material, Figs S3A-S3X show also the diagnostic chromatograms for the individual dogs.

**Fig 3A.**
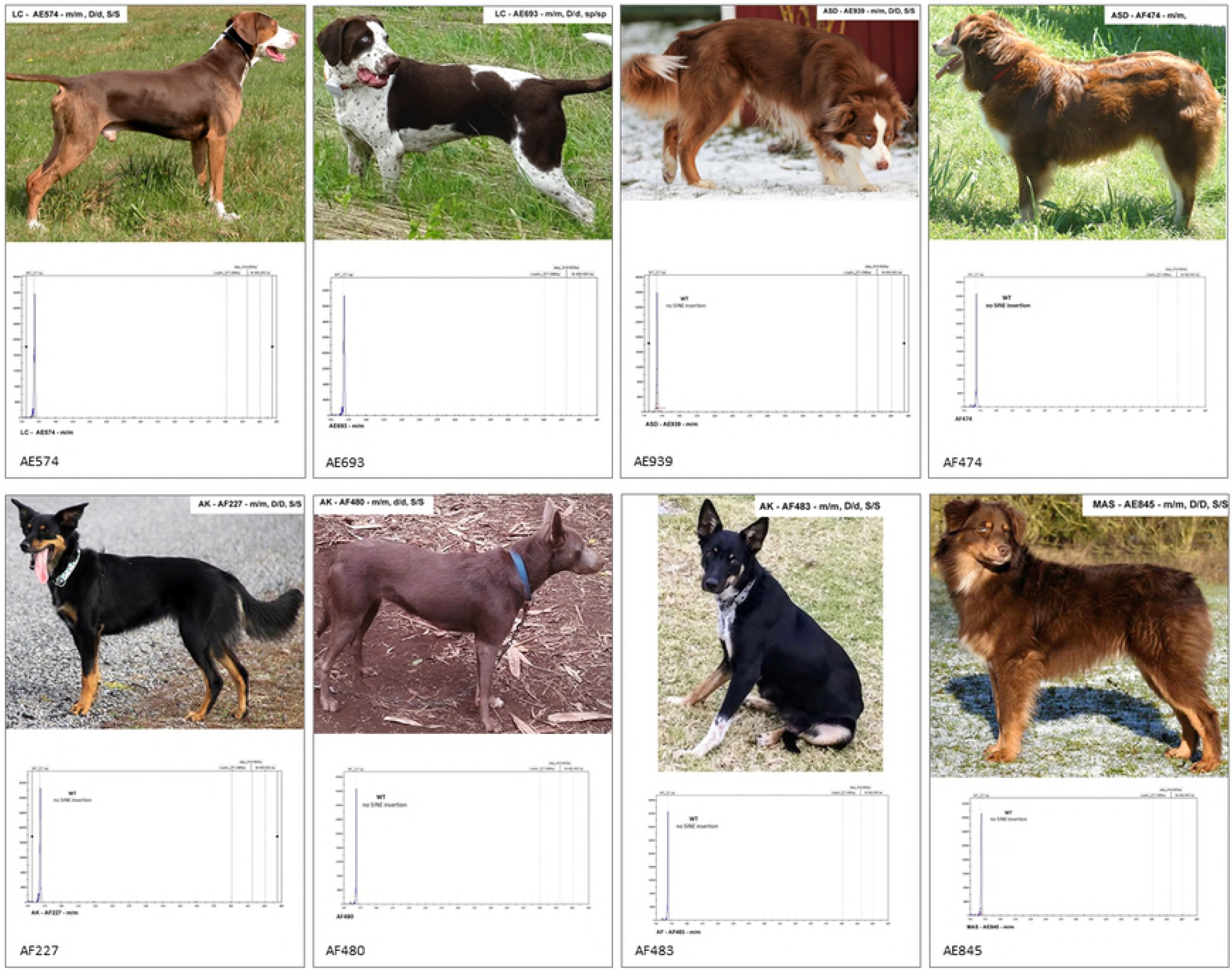
Genotype/phenotype correlations: m/m. Non-Merle, no *SILV* SINE insertion

**Fig 3B.**
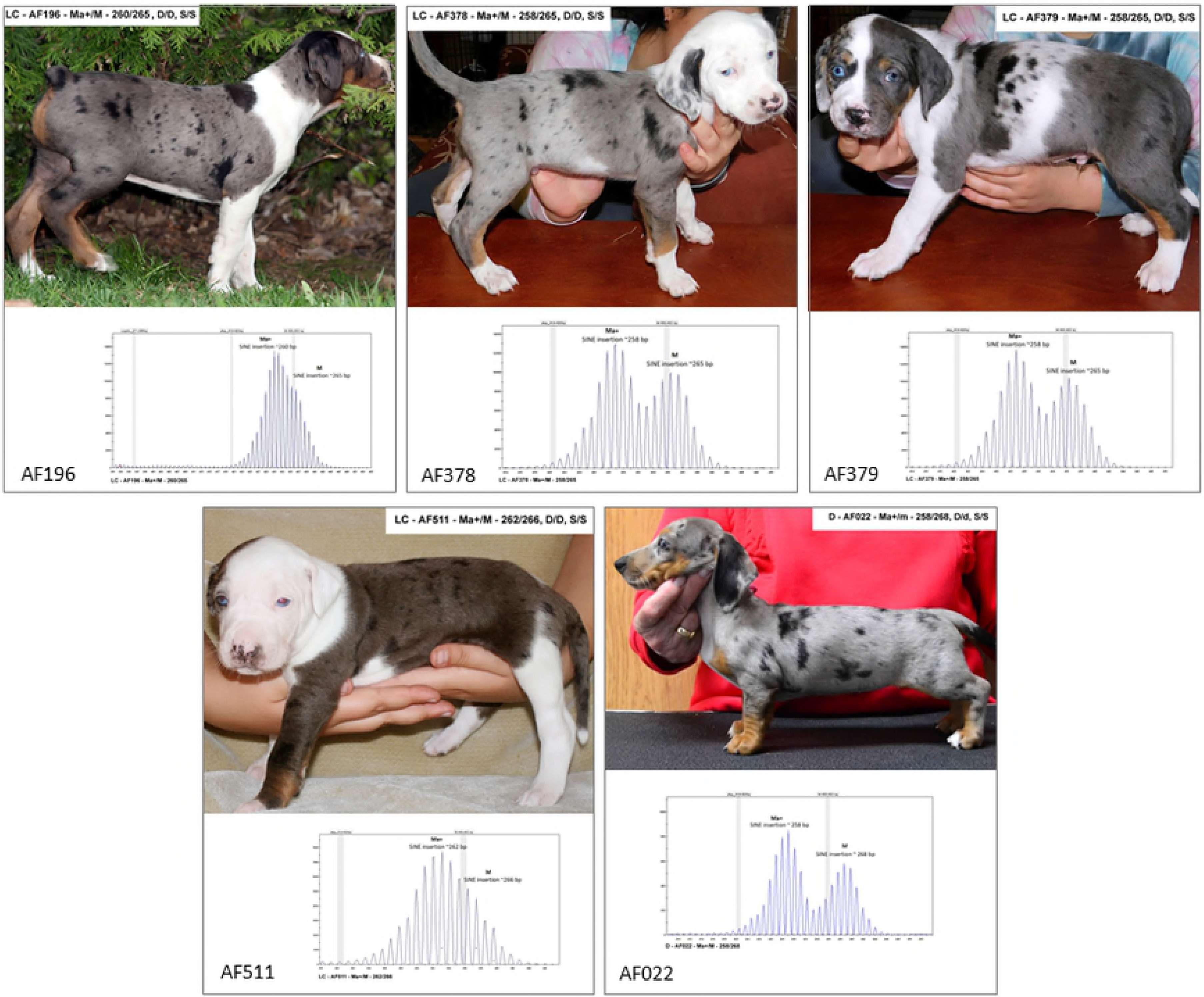
Genotype/phenotype correlations: m/Mc. No Merle pattern, no change to coat color or pigment shading. No eye color change. No pigment is deleted to white.

**Fig 3C.**
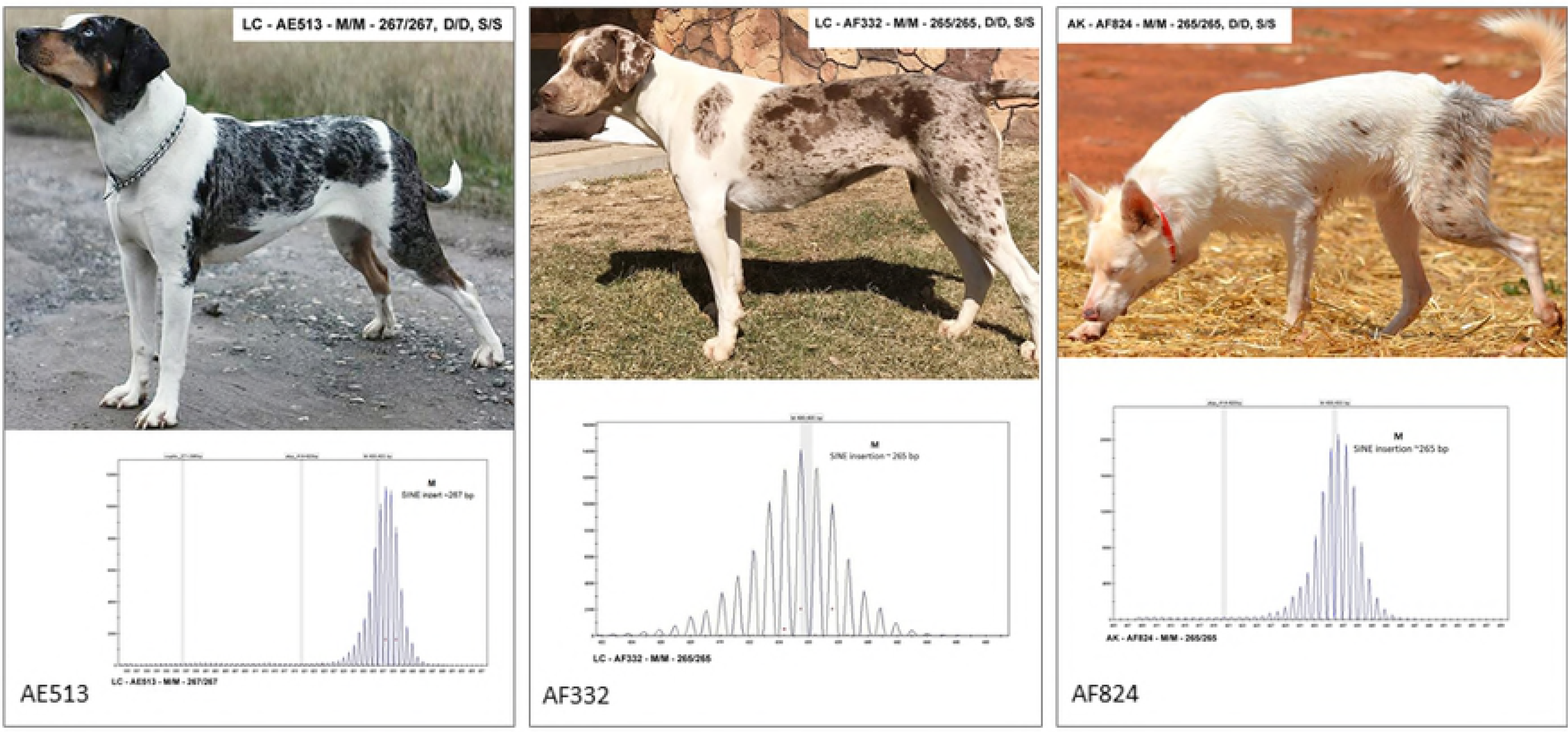
Genotype/phenotype correlations: Mc/Mc. No Merle pattern, may express as no change to coat color or pigment shading. Alternatively, there may be a slight change to coat color – pigment may express as faded or off-color or a slight brownish hue may express that is not related to b/b, especially for long coated breeds. No eye color change. No pigment is deleted to white.

**Fig 3D.**
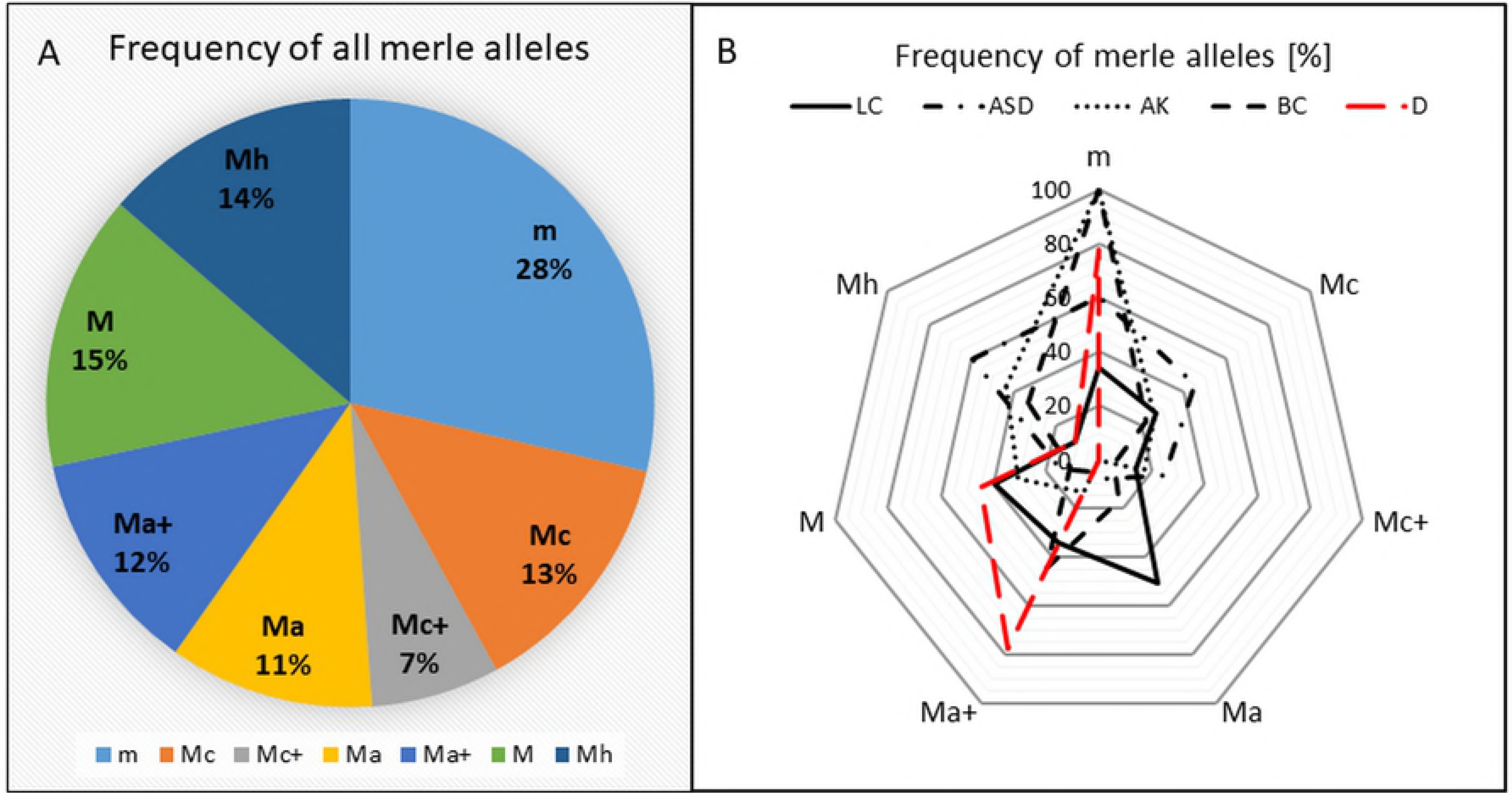
Genotype/phenotype correlations: m/Mc+. No Merle pattern, no change to coat color or pigment shading. No eye color change. No pigment is deleted to white.

**Fig 3E.**
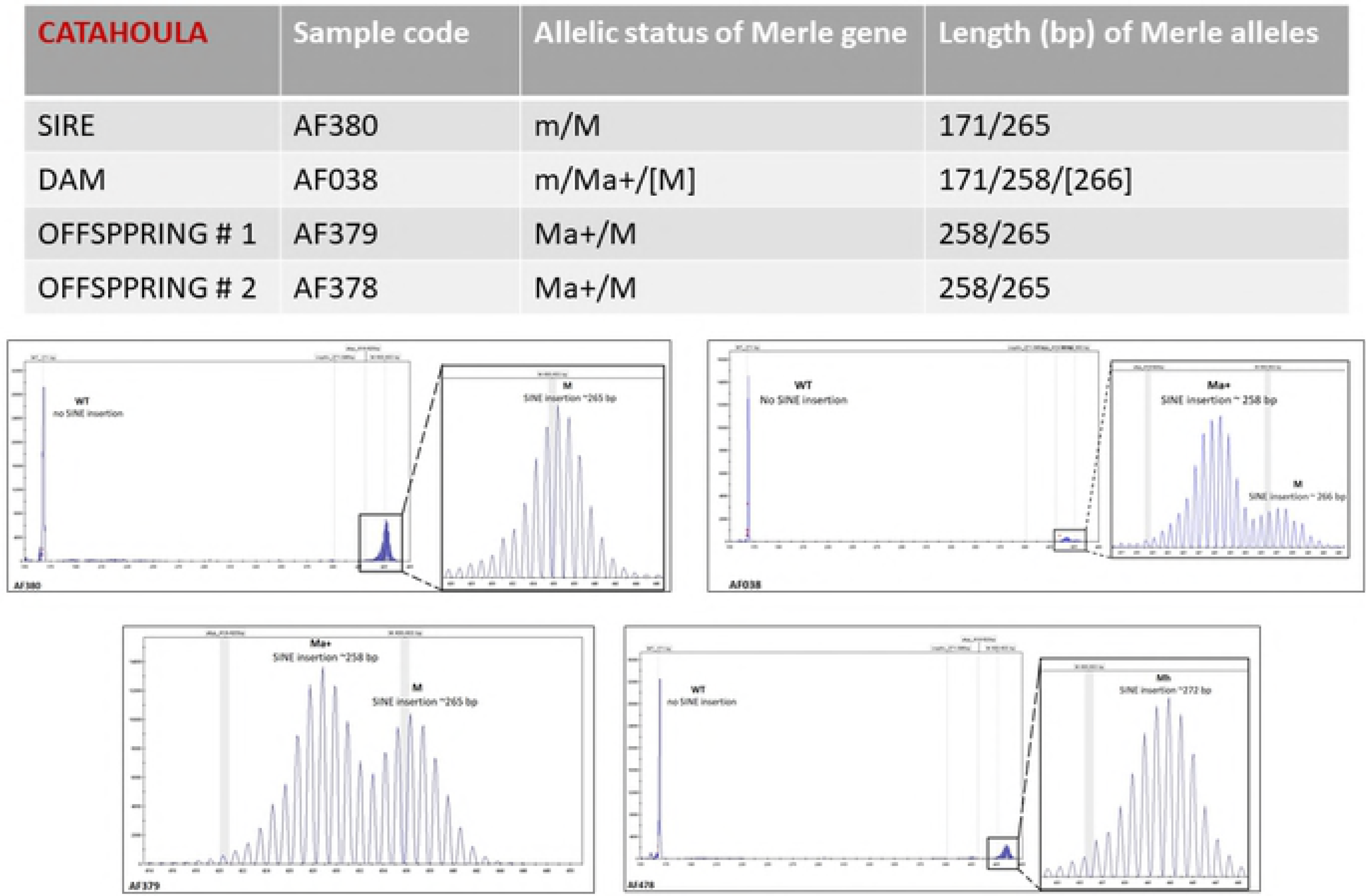
Genotype/phenotype correlations: Mc+/Mc+. No Merle pattern, may express as no change to coat color or pigment shading. Alternatively, there may be a slight change to coat color – pigment may express as faded or off-color or a brownish hue may express that is not related to b/b, especially for long coated breeds. No eye color change. No pigment is deleted to white.

**Fig 3F.**
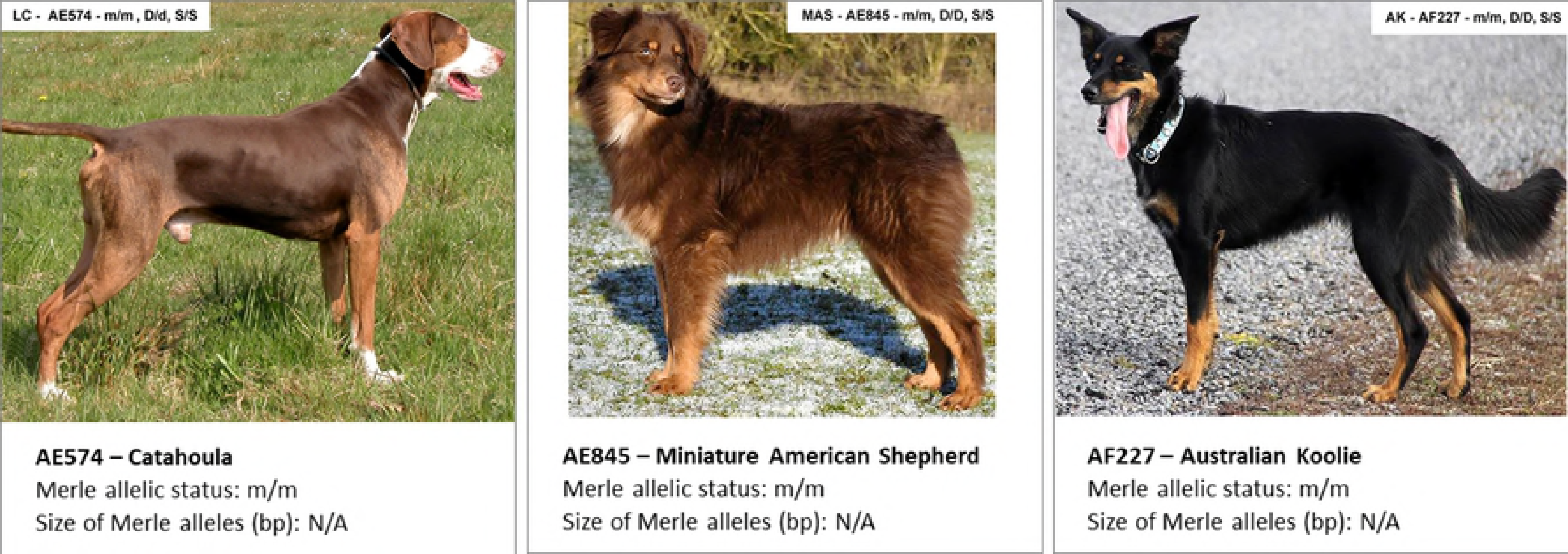
Genotypephenotype correlations: m/Ma. No Merle pattern, may express with no change to coat color or pigment shading. Alternatively, may show a diluted coat expression even when d/d is not present and/or a brownish hue may express that is not related to b/b. May express with a lighter undercoat especially on longer haired breeds. Lighter shaded areas may be visible on ears, neck, under tail and tail area. Blue eyes can be expressed. No pigment is deleted to white.

**Fig 3G.**
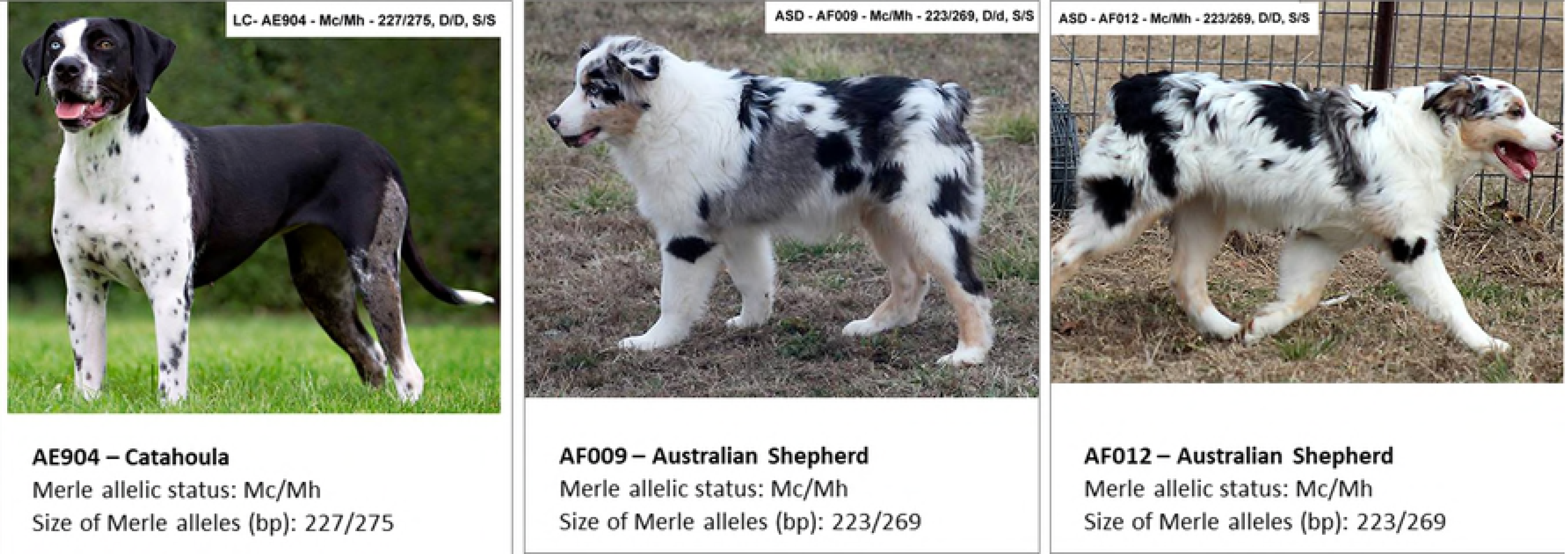
Genotypephenotype correlations: Mc/Ma. No Merle pattern, may express with no change to coat color or pigment shading. Alternatively, may show a diluted coat expression even when d/d is not present and/or a brownish hue may express that is not related to b/b or express with a lighter undercoat especially on longer haired breeds. Lighter shaded areas may be visible on ears, neck, under tail and tail area. Blue eyes can be expressed. No pigment is deleted to white.

**Fig 3H.**
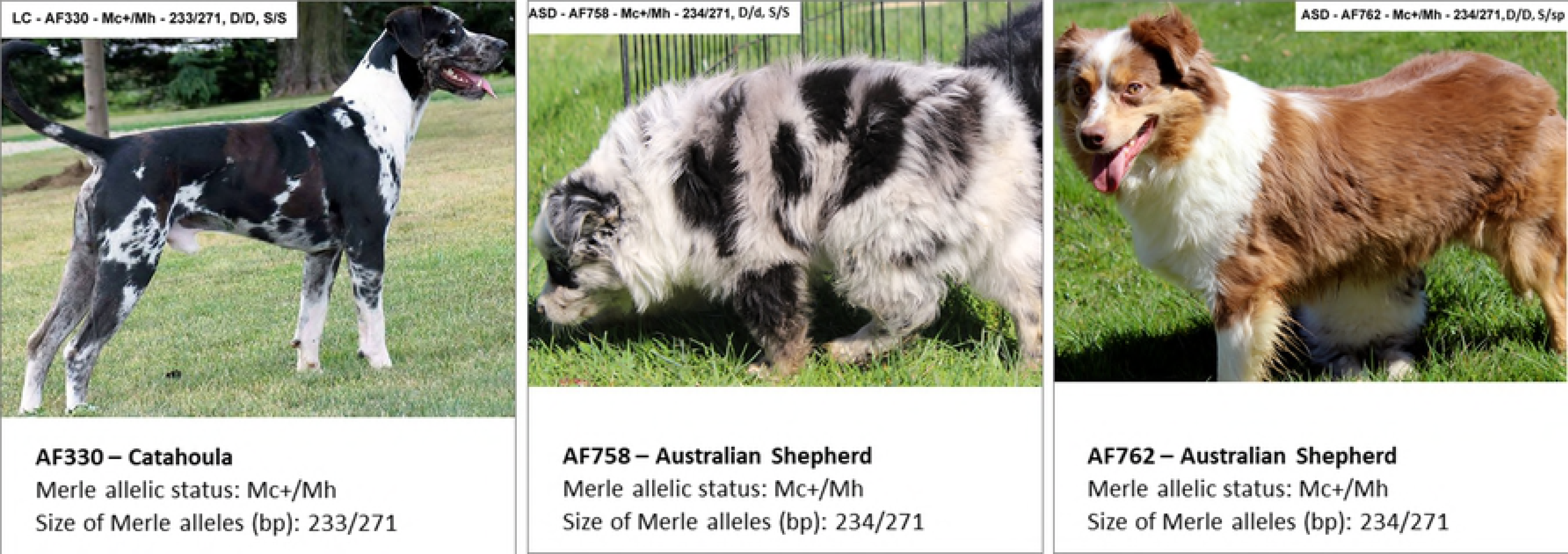
Genotype/phenotype correlations: Ma/Ma. Most often diluted in color even when d/d is not present and/or a brownish hue may express that is not related to b/b, more diluted background shading with smaller and fewer areas of darker spotting. Blue eyes can be expressed. No pigment is deleted to white.

**Fig 3I.**
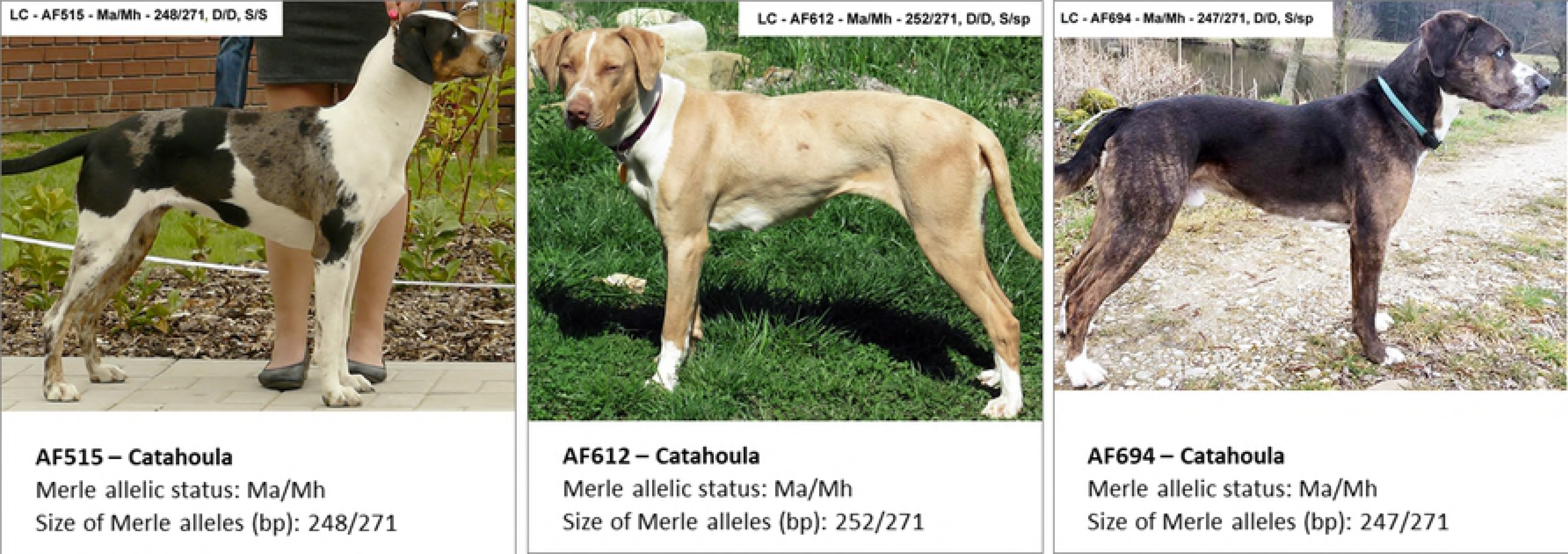
Genotype/phenotype correlations: m/Ma+. Merle pattern is muted, not crisp and clear or as well defined as some breed standards may require, most often diluted in color even when d/d is not present and/or a brownish hue may express that is not related to b/b. Alternatively, some dogs may express with no Merle pattern, no dilution and no change to coat color or pigment shading. Blue eyes can be expressed. No pigment is deleted to white

**Fig 3J.**
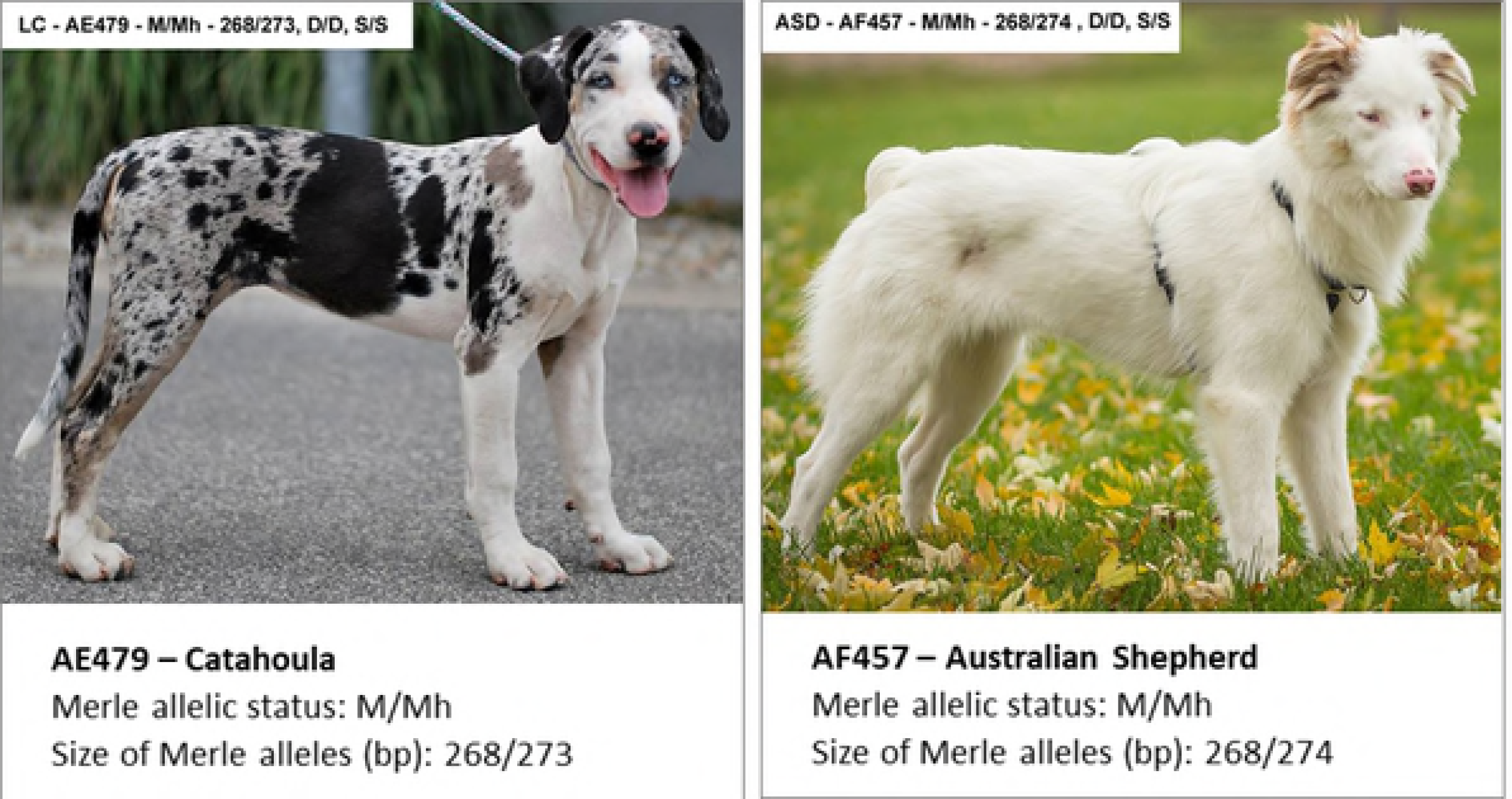
Genotype/phenotype correlations: Mc/Ma+. Merle pattern is muted, not crisp and clear or as well defined as some breed standards may require, most often diluted in color even when d/d is not present and/or a brownish hue may express that is not related to b/b. Alternatively, some dogs may express with no Merle pattern, no dilution and no change to coat color or pigment shading. Blue eyes can be expressed. No pigment is deleted to white.

**Fig 3K.**
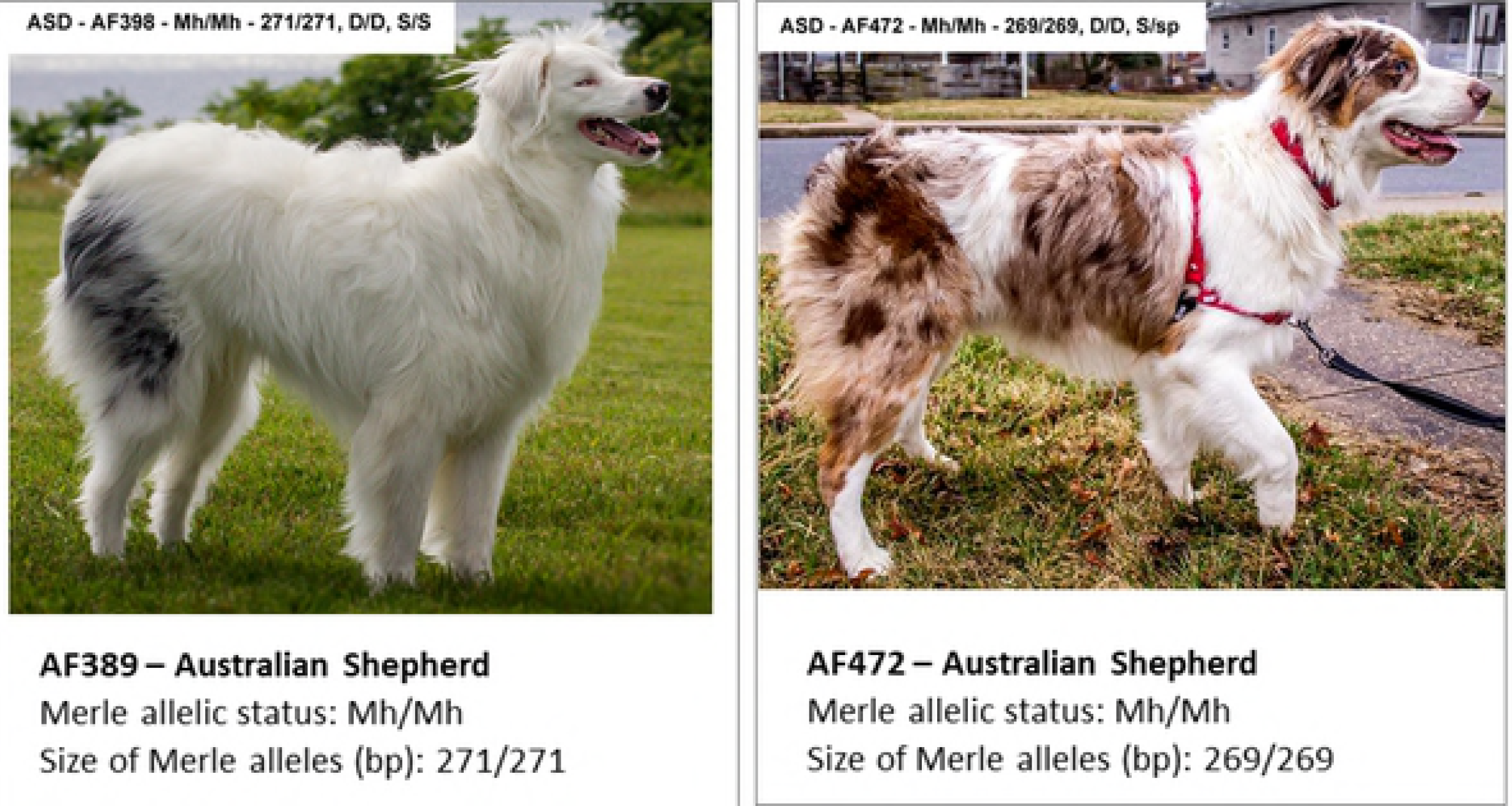
Genotype/phenotype correlations: Mc+/Ma+. Often diluted in color even when d/d is not present and/or a brownish hue may express that is not related to b/b, more diluted background shading with smaller and fewer areas of spotting. As the base pairs of Ma+ progress closer to M, a more noticeable Tweed patterning may be present, larger areas of solid pigment may show. Blue eyes can be expressed. Some pigment may be deleted to white as the base pair numbers of Ma+ progress closer to M.

**Fig 3L.**
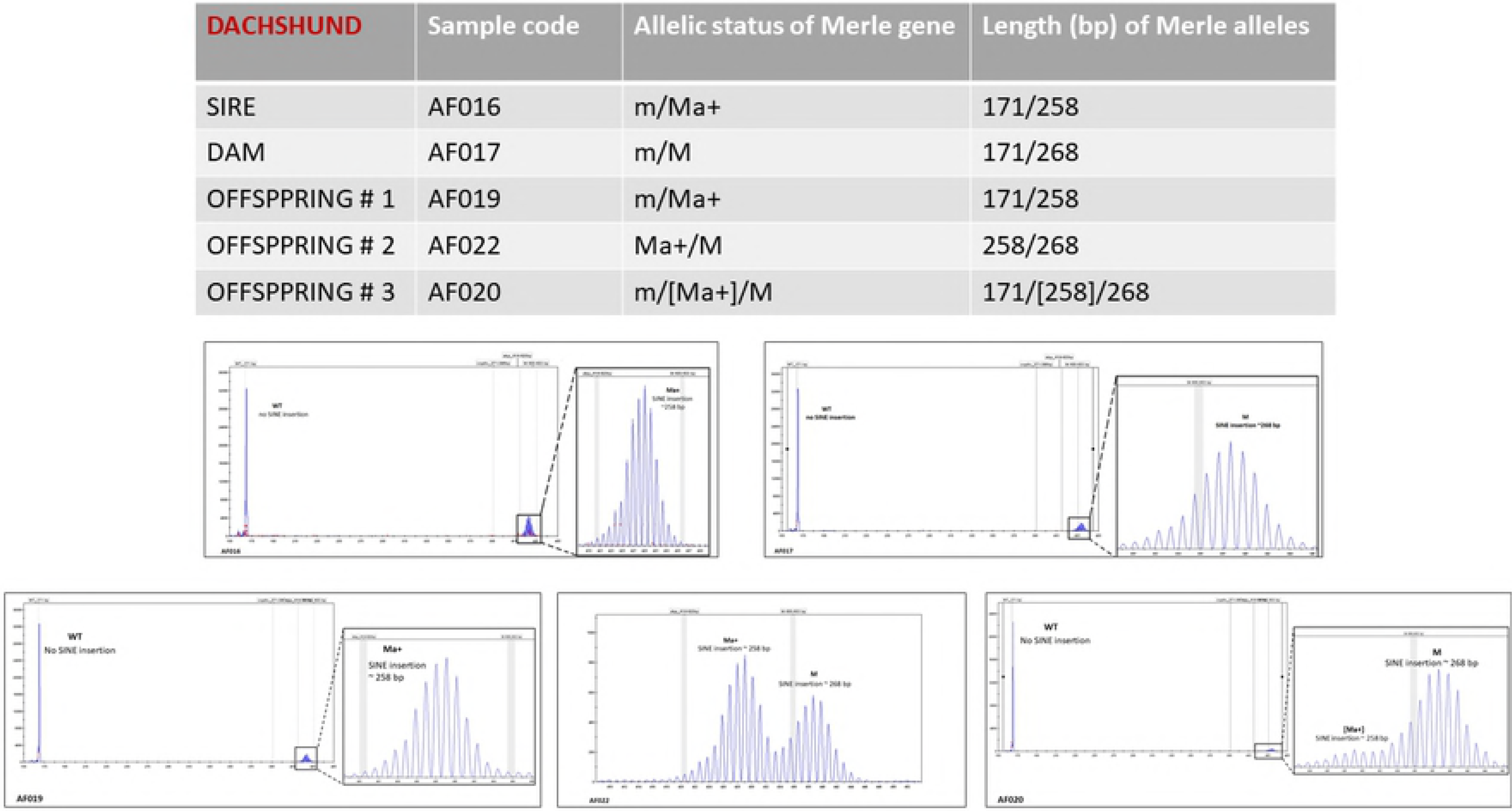
Genotype/phenotype correlations: Ma+/Ma+. Most often diluted in color even when d/d is not present and/or a brownish hue may express that is not related to b/b. More diluted background shading with smaller and fewer areas of spotting. Tweed patterning may be present. Blue eyes can be expressed. Pigment may be deleted to white.

**Fig 3M.**
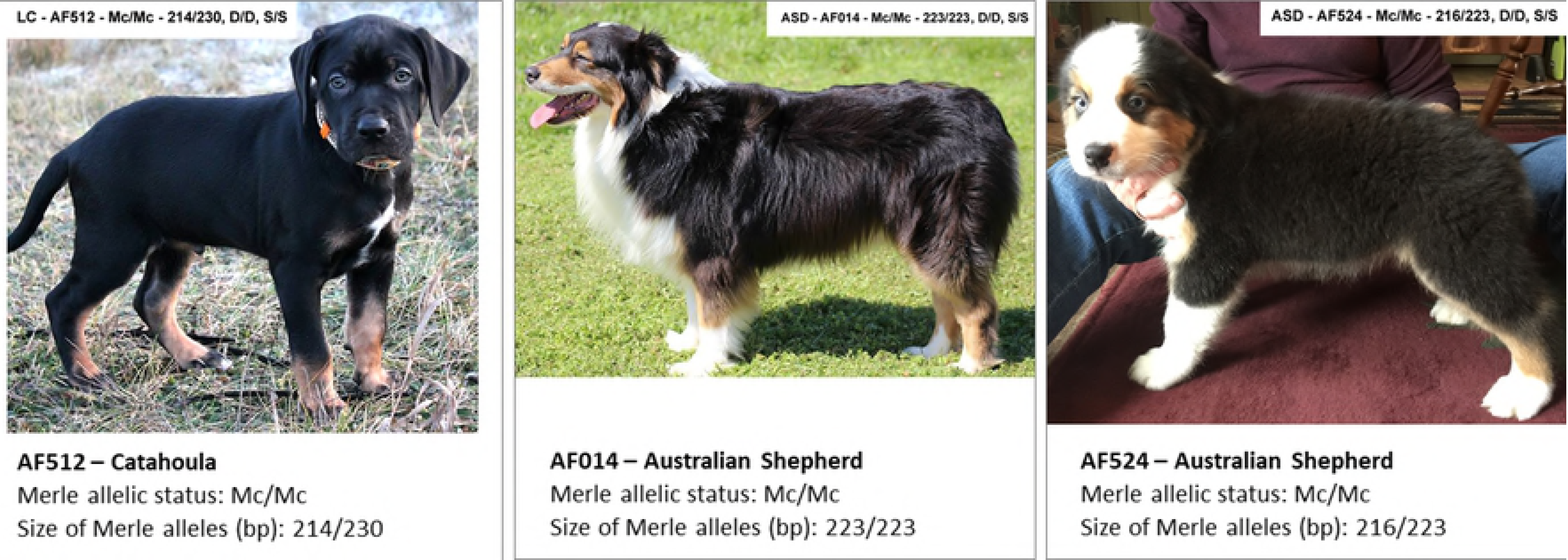
Genotype/phenotype correlations: m/M. Classic Merle pattern – random areas of the coat are diluted to a lighter pigment, creating a combination of areas consisting of a diluted color mixed with areas of full pigmentation. Blue eyes can be expressed. No pigment is deleted to white.

**Fig 3N.**
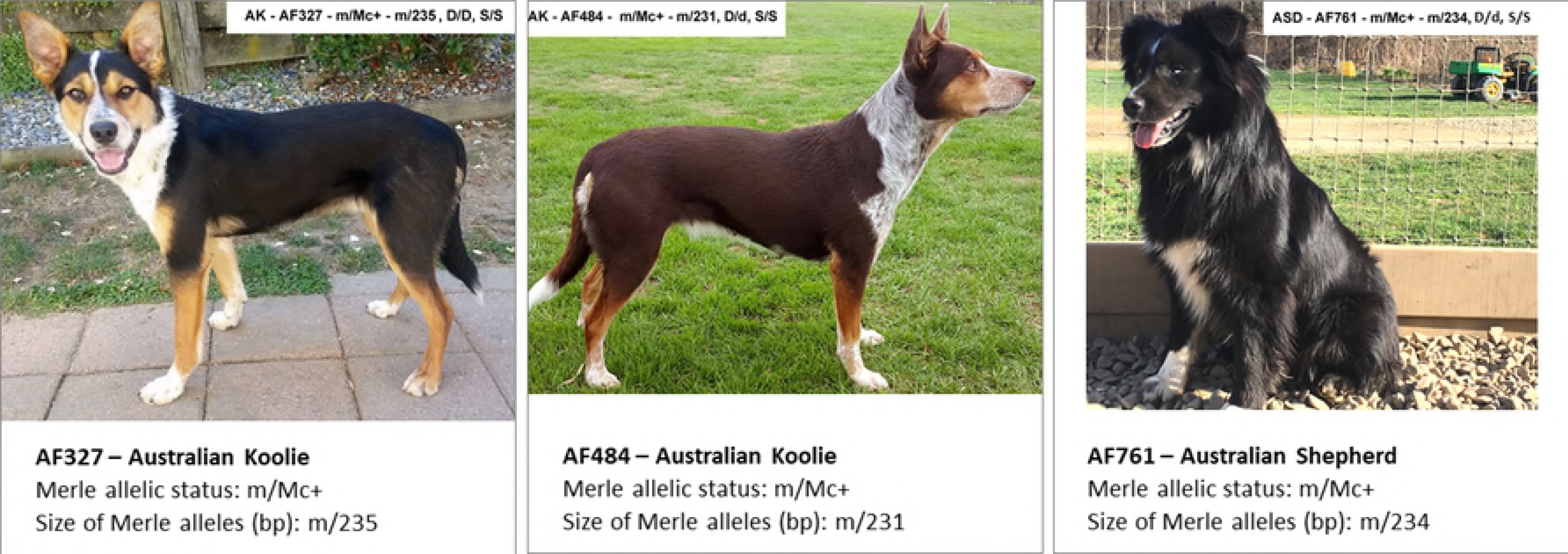
Genotype/phenotype correlations: Mc/M. Random areas of the coat are diluted to a lighter pigment, creating a combination of areas consisting of a diluted color mixed with areas of full pigmentation. Tweed patterning may express. Blue eyes can be expressed. No pigment is deleted to white.

**Fig 3O.**
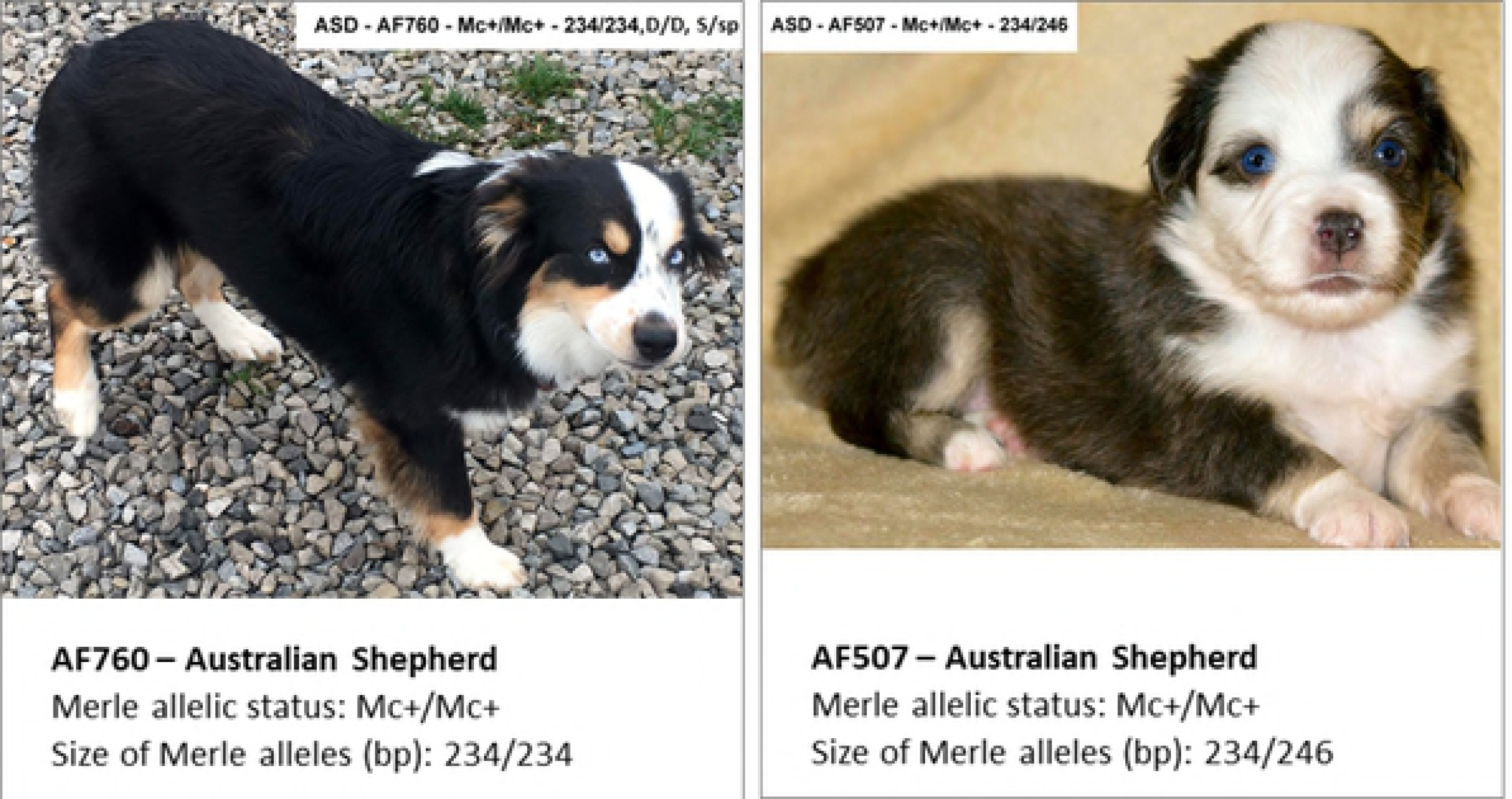
Genotype/phenotype correlations: Mc+/M. Random areas of the coat are diluted to a lighter pigment, creating a combination of areas consisting of a diluted color mixed with areas of full pigmentation. Tweed patterning may express. Blue eyes can be expressed. Some pigment may be deleted to white.

**Fig 3P.**
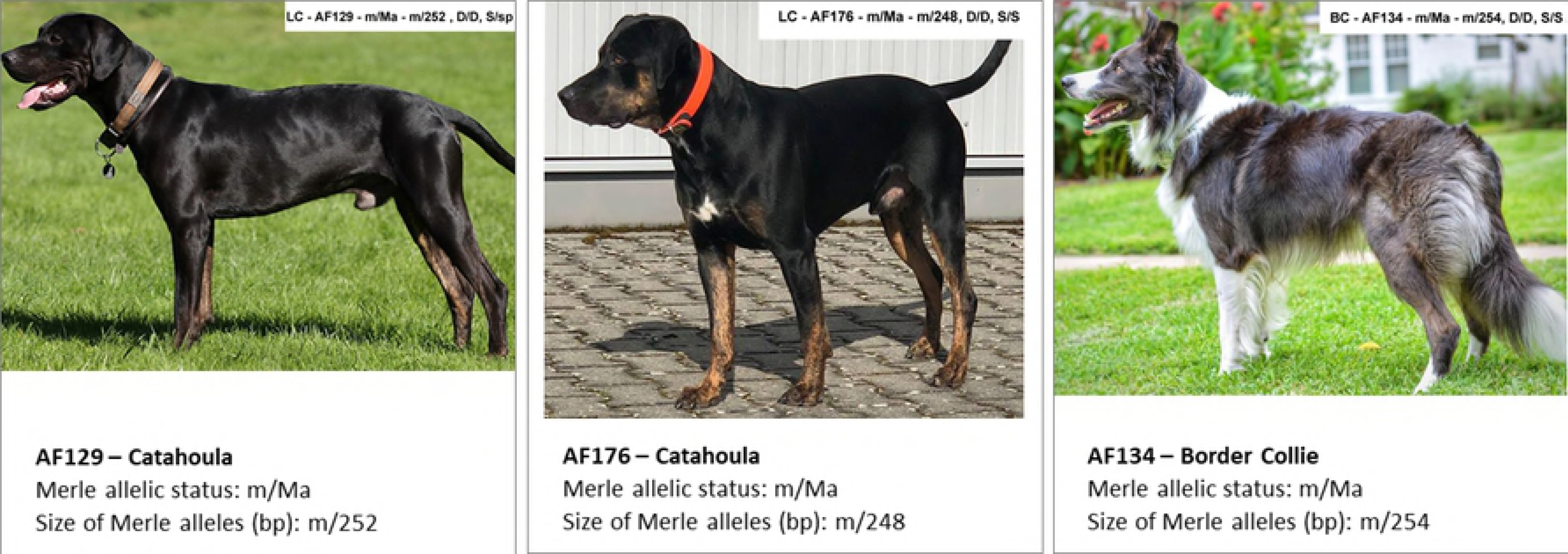
Genotype/phenotype correlations: Ma/M. Often referred to as “Patchwork” with large areas of solid pigment mixed with areas of more diluted background shading with smaller and fewer areas of darker spotting. Tweed patterning often expressed. Blue eyes can be expressed. Some pigment often deleted to white.

**Fig 3Q.**
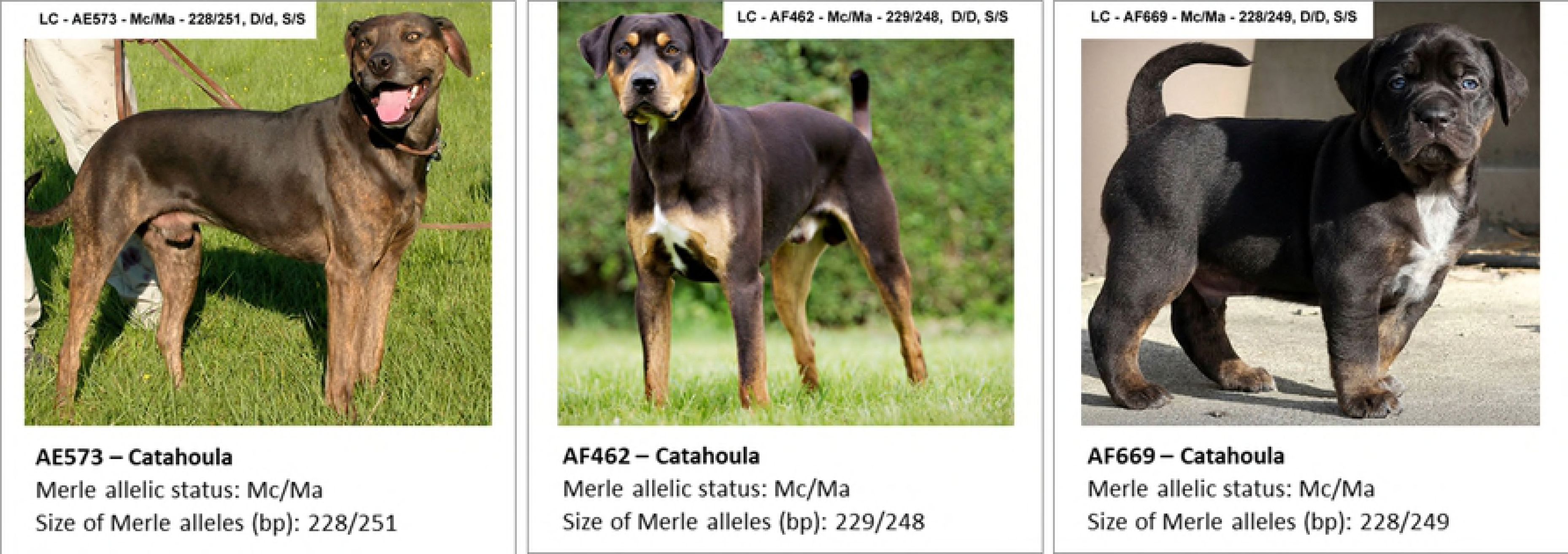
Genotype/phenotype correlations: Ma+/M. Most often diluted in color even when d/d is not present and/or a brownish hue may express that is not related to b/b. More diluted background shading with smaller and fewer areas of spotting. Extended white out of normal Irish Spotting pattern – up legs, past shoulders, white head often noted (not related to the “ white-head” gene). Blue eyes can be expressed. Pigment can be deleted to white.

**Fig 3R.**
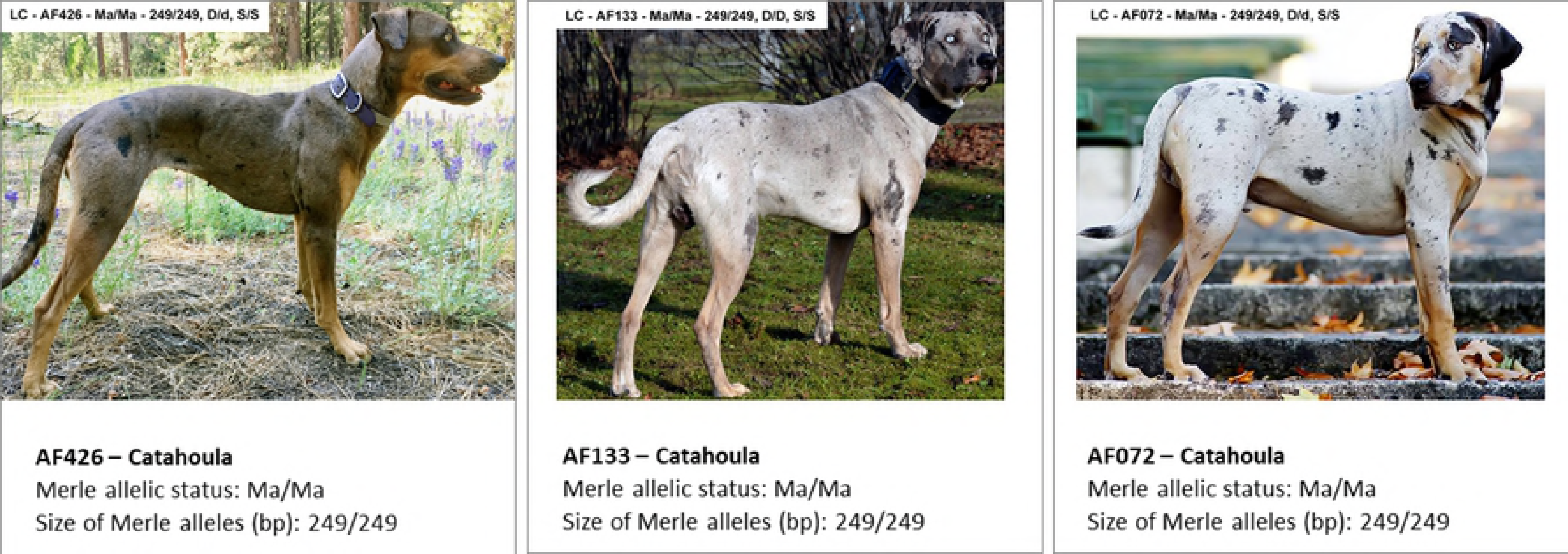
Genotype/phenotype correlations: M/M. Random areas of the coat are diluted to a lighter pigment, creating a combination of areas consisting of a diluted color mixed with areas of full pigmentation most often mixed with varying amounts of white. Blue eyes can be expressed. Pigment can be deleted to white.

### Genotype/phenotype correlations: Mh

The Mh allele has a broad range of phenotypes with 2 expressions that are very recognizable.

#1 - “Minimal Merle” - a large percentage of the body features solid colored pigment with only small random areas of Merle patterning. Individuals may also express extended white out of the normal area of the typical Irish Spotting pattern – this may include a large white collar, white up legs past the elbow, white past shoulders extending onto withers and white on the belly extending up the side. This extended white is sometimes associated with S/sp - (Piebald Carrier), however many m/Mh dogs with this type of white pattern have tested as S/S.

#2 - The more classic pattern that is often referred to as “Herding Harlequin” - Random diluted areas of Merle pigment are deleted to white, leaving solid patched areas that may be Tweed patterned including different shades. Some Merle areas may remain. The extended white patterning mentioned in description #1 may be present but is less noticeable due to the deleted white areas on the body.

#3 - Some dogs may express more as m/M, yet are still able to produce offspring with a phenotype as described above in example #1 and #2 - these offspring have inherited the same length of base pairs as the parent and yet express in either of the 3 ways presented here. See **Chart 1**.

**Chart 1:** In the following example, the dam - AF762 – is distinguished by a Mh phenotype as described in #1 - “Mininal Merle”. The two offspring – AF758 and AF759 both have a Mh phenotype as described in #2 - The more classic pattern that is often referred to as “Herding Harlequin” ”. Both dam and offspring have the same length of Mh - 271 bp.

Mc/Mh, Mc+/Mh, Ma/Mh, M/Mh and Mh/Mh allelic combinations are phenotypically indistinguishable and present one homogenous phenotypic group.

Of note, M/Mh and Mh/Mh may express with a greater percentage of white over the body.

**Fig 3S.**
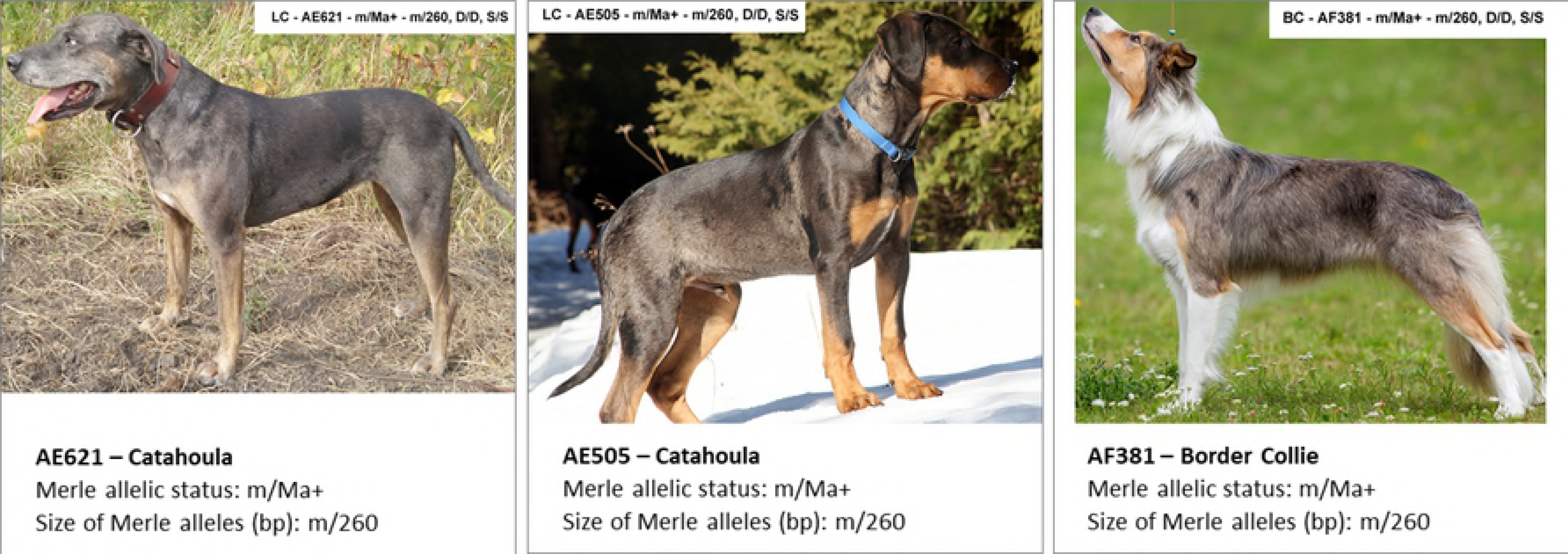
Genotype/phenotype correlations: m/Mh.

**Fig 3T.**
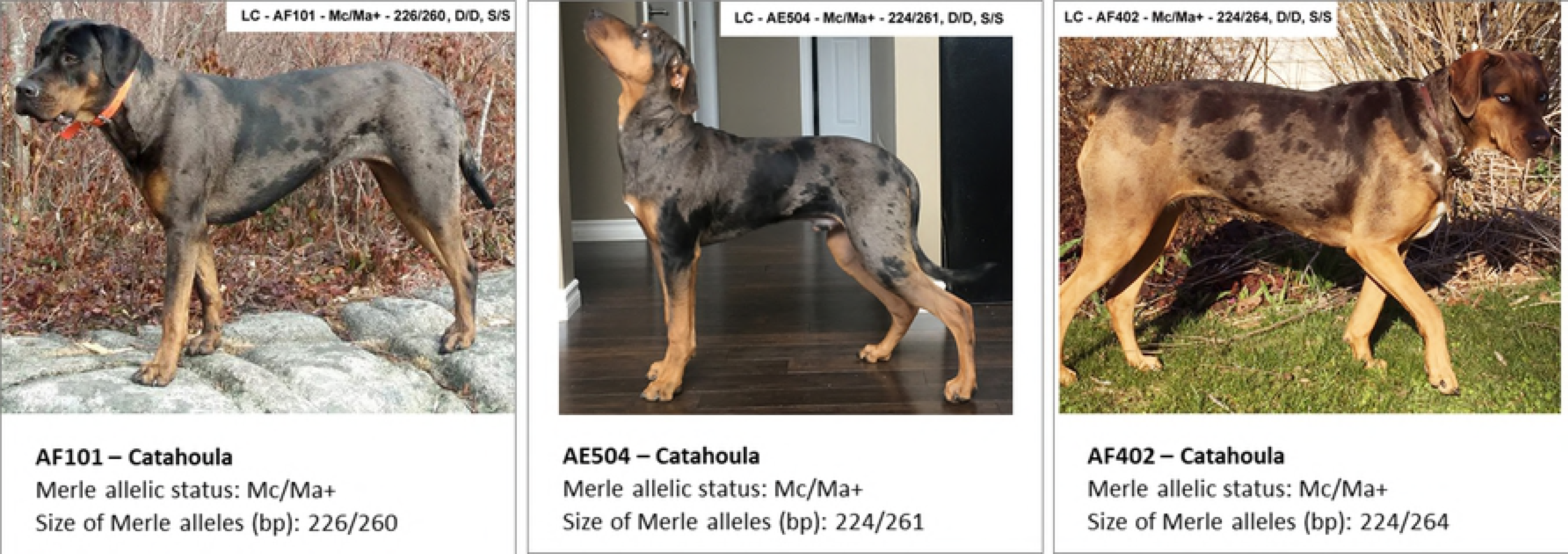
Genotype/phenotype correlations: Mc/Mh.

**Fig 3U.**
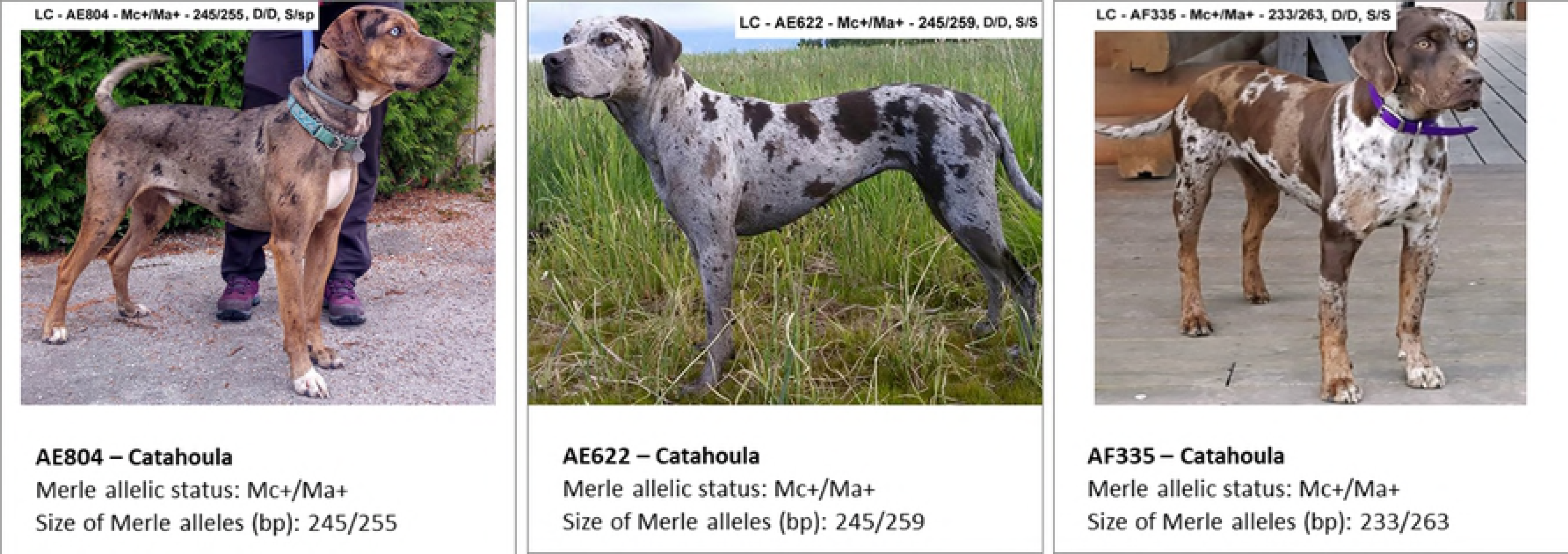
Genotype/phenotype correlations: Mc+/Mh.

**Fig 3V.**
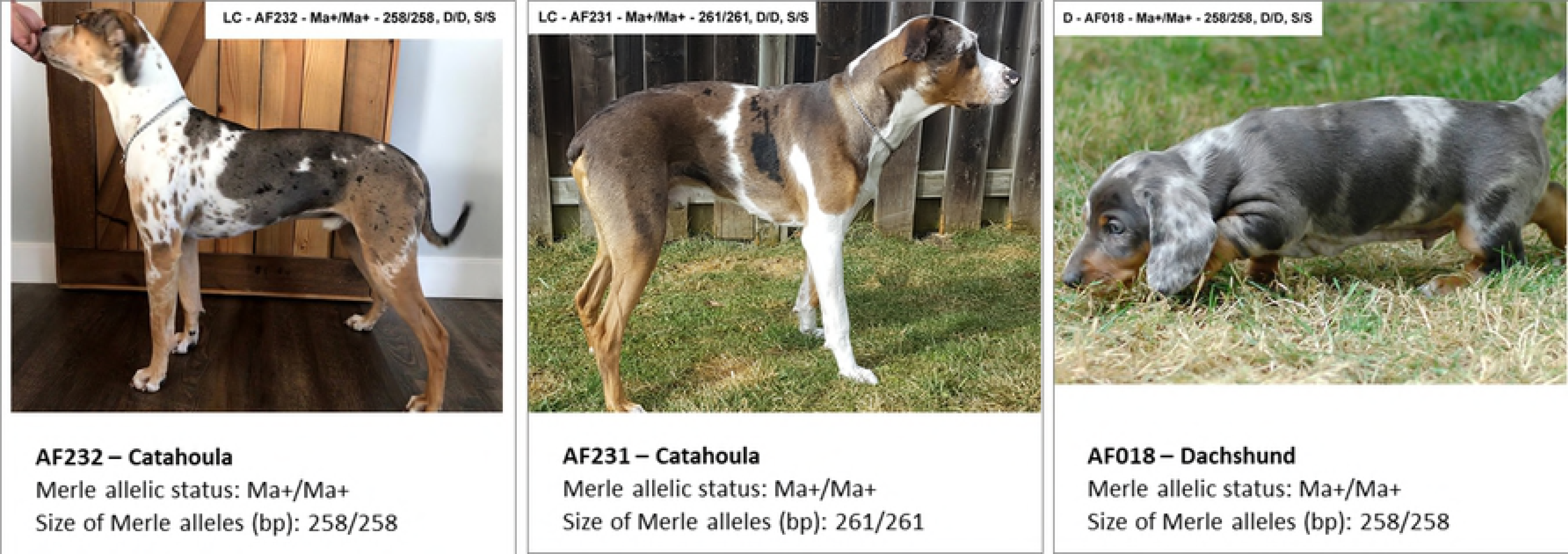
Genotype/phenotype correlations: Ma/Mh.

**Fig 3W.**
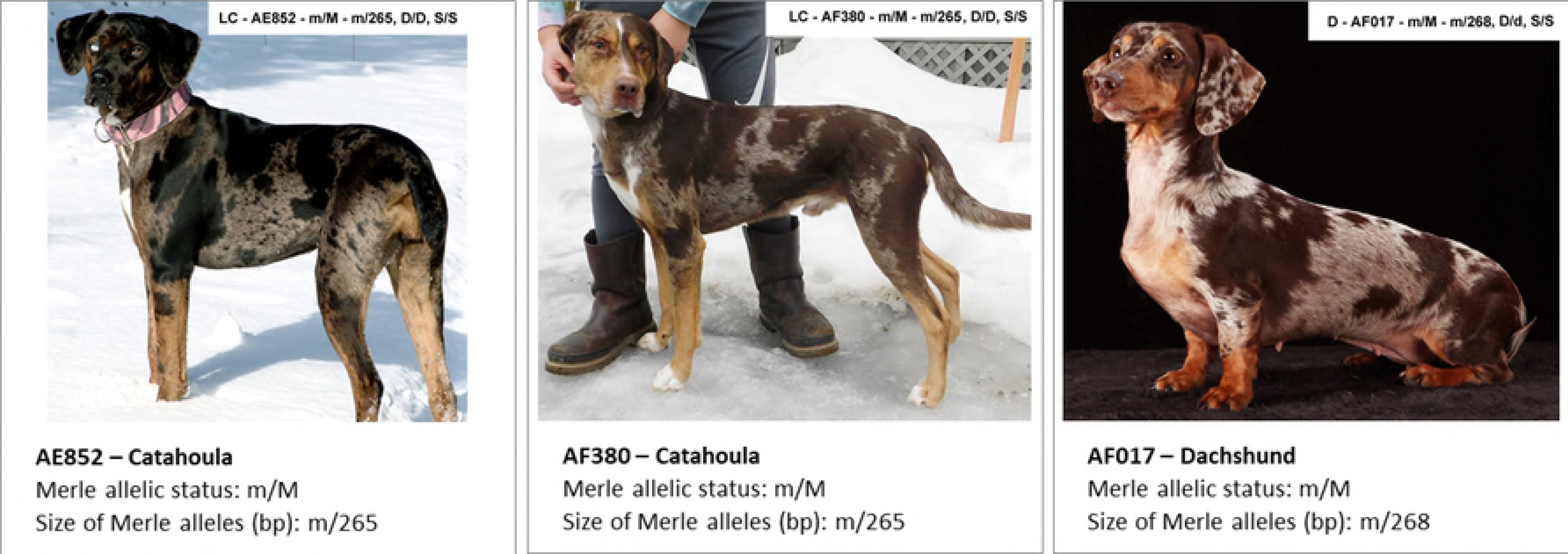
Genotype/phenotype correlations: M/Mh.

**Fig 3X.**
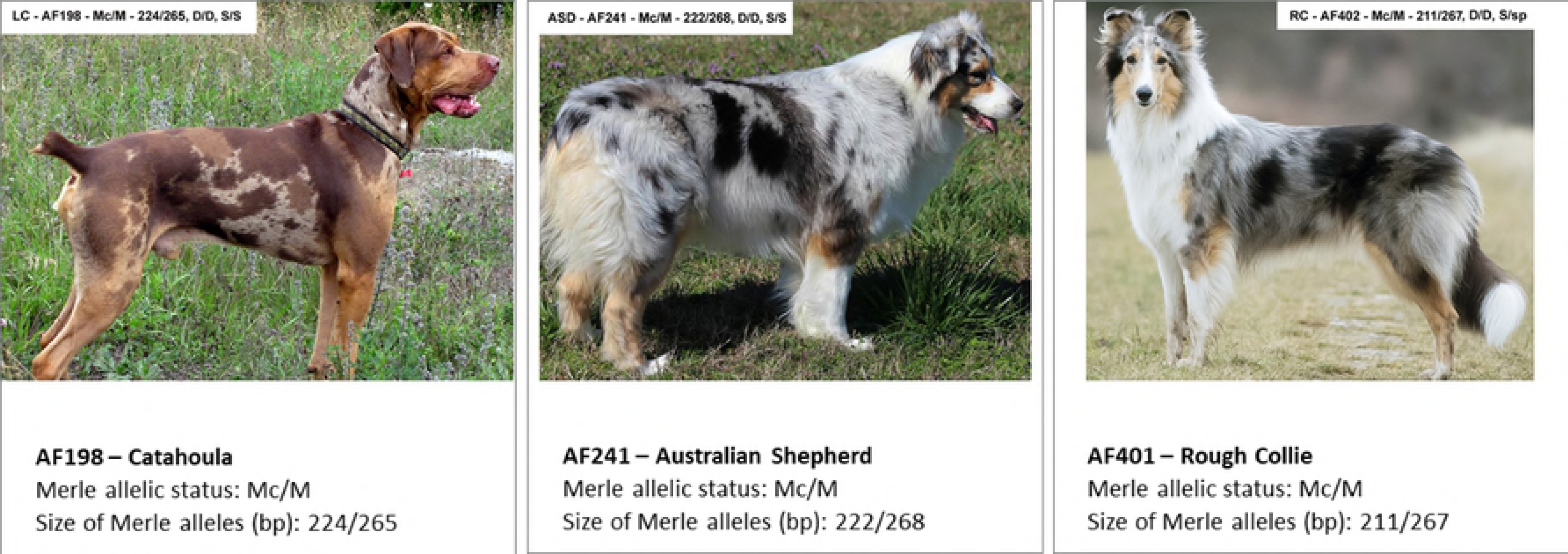
Genotype/phenotype correlations: Mh/Mh.

The following Table 2 summarizes the major phenotypic differences among individual Merle alleles.

**Table 2.** Major differences in phenotypes caused by individual Merle alleles.

| BREED | n | Mc | Mc+ | Ma | Ma+ | M | Mh |
| --- | --- | --- | --- | --- | --- | --- | --- |
| LC | 73 | 208–230 | 233–245 | 247–253 | 255–264 | 265–268 | 271–277 |
| ASD | 40 | 216–230 | 233–243 | 252–254 | 259–259 | 266–268 | 269–274 |
| AK | 23 | 213–230 | 231–235 |  | 262–264 | 265–268 | 269–278 |
| BC | 18 | 222–225 | 246 | 254 | 256–262 | 268 | 269–274 |
| D | 9 |  |  |  | 258–260 | 268 | 274 |
| RC | 3 | 211 |  |  | 260 | 267 |  |
| WS | 3 |  | 231 |  |  | 268 | 272–273 |
| MAUS | 2 | 217 |  |  | 262 |  | 269–273 |
| MAS | 2 |  |  |  |  | 267 |  |
| FB | 2 |  |  |  |  | 265 |  |
| SSD | 2 | 220–223 |  |  |  |  | 271 |
| M | 2 | 226 |  |  |  | 267 |  |
| PS | 1 |  |  |  |  | 265 |  |
| LD | 1 |  |  |  |  | 266 |  |
| Average range | 181 | 208–230 | 231–245 | 247–254 | 255–264 | 265–269 | 269–277 |

### Abundance of the individual Merle alleles

We have calculated the relative frequency of the individual Merle alleles in our cohort of 181 dogs. As shown in Fig 4, among all breeds tested in our study, dogs heterozygous for the wild type m allele were the most frequent (approximately one third of all individuals tested); M, Mh, and others followed (Fig 4A). However, significant differences in their abundance have been revealed in the five most populated breeds (Fig 4B). LC carried Ma, M, and Ma+ (50.7, 39.7, and 34% of tested dogs, respectively), while the m allele was present only in 34% of the individuals tested. In contrast, most of AK, BC, D, and ASD were heterozygous for the m allele, as 100, 100, 78, and 60% of the tested dogs, respectively, carried it. ASD showed besides m and Mh (60%) also Mc (45%) as the most frequent allele. In BC, Ma+ (44%) was the most frequent allele, followed by m, Mh (33%), and Mc (22%). In our cohort of dogs, D have been found free of Mc, Mc+, and Ma alleles, while Ma+ (78%) was the most abundant allele, followed with M (44%).

**Fig 4:**
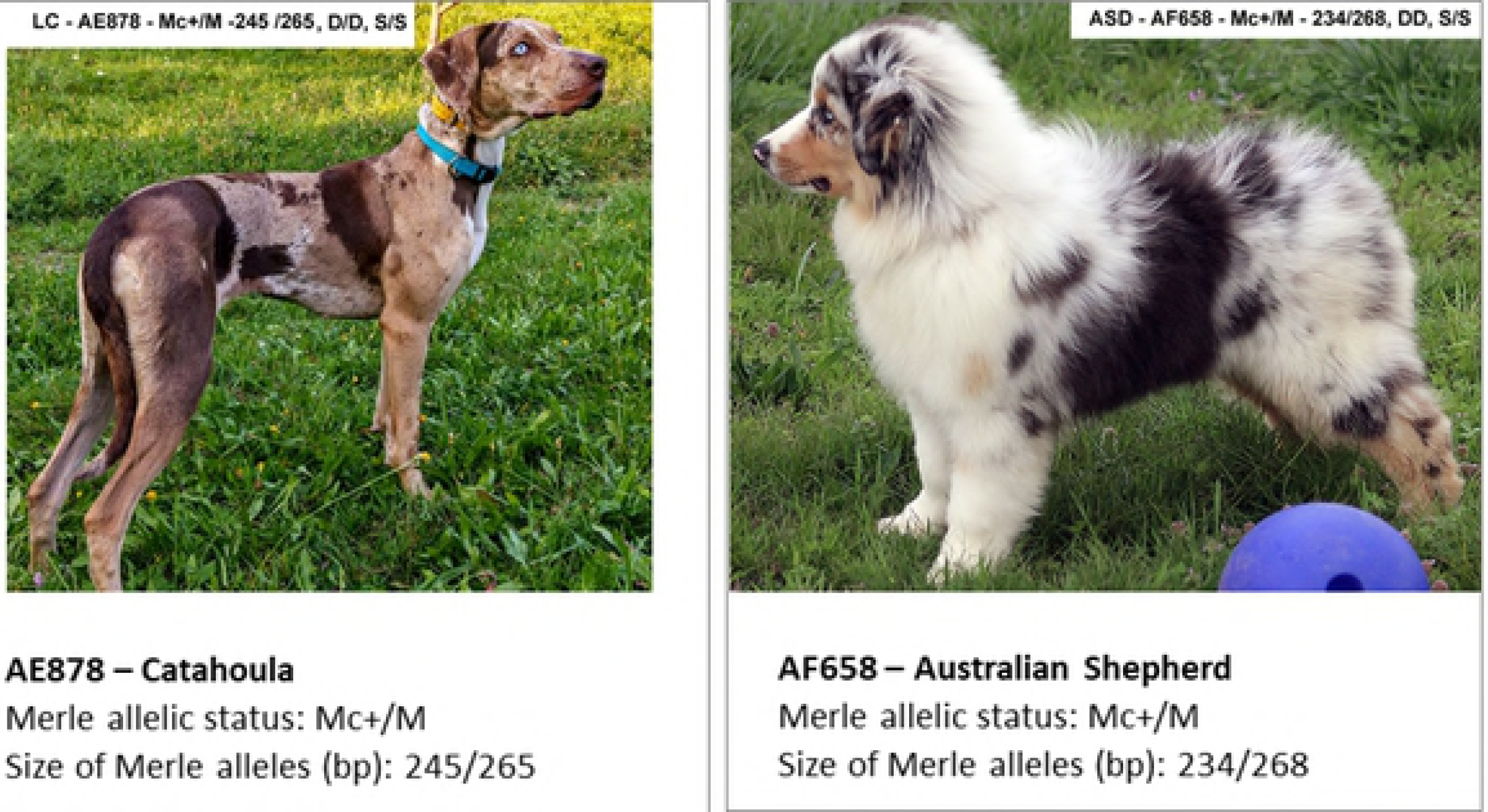
**A:** Relative frequency [%] of all merle alleles in all breeds tested. Percentage was calculated from all alleles found in all breeds involved in the study. **B:** Frequency [%] of all individual alleles in five most numerous breeds (with *n* ≥ 9), calculated as percent of all alleles found in the particular breed. ASD – Australian Shepherd Dog, AK – Australian Koolie, BC – Border Collie, D – Dachshund, LC – Louisiana Catahoula.

Frequency of all individual alleles in all breeds tested are summarized in Fig 5. The Mc allele was found in eight breeds, while the Mc+ allele was found in six breeds. The Ma allele was detected only in three breeds (Fig 4B), while the Ma+ allele was present in seven breeds (Table 1).

**Fig 5.**
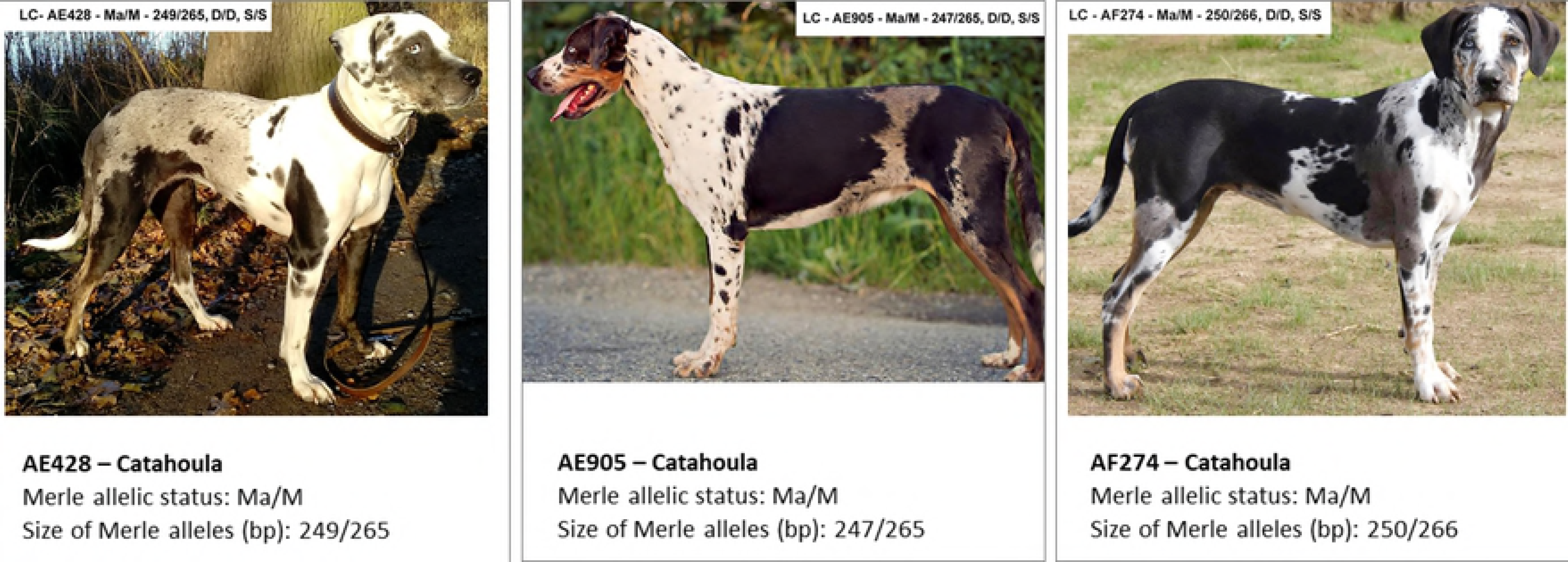
Relative frequency [%] of Mc (A), Mc+ (B), Ma (C), and Ma+ (D) alleles in individual breeds and in all dogs tested. ASD – Australian Shepherd Dog, AK – Australian Koolie, BC – Border Collie, D – Dachshund, LC – Louisiana Catahoula.

The Mh allele was found in 8 breeds out of 14 tested (Fig. 6). Beside MAUS, WS, and SSD (*n* = 2–3), where Mh was carried by 50–100% of tested dogs, the highest frequency of Mh was observed in ASD (60%), followed by AK (43.5%), and BC (33%). On the other hand, in LC, the most populated breed in our study, only 11% of the individuals tested carried Mh.

**Fig 6.**
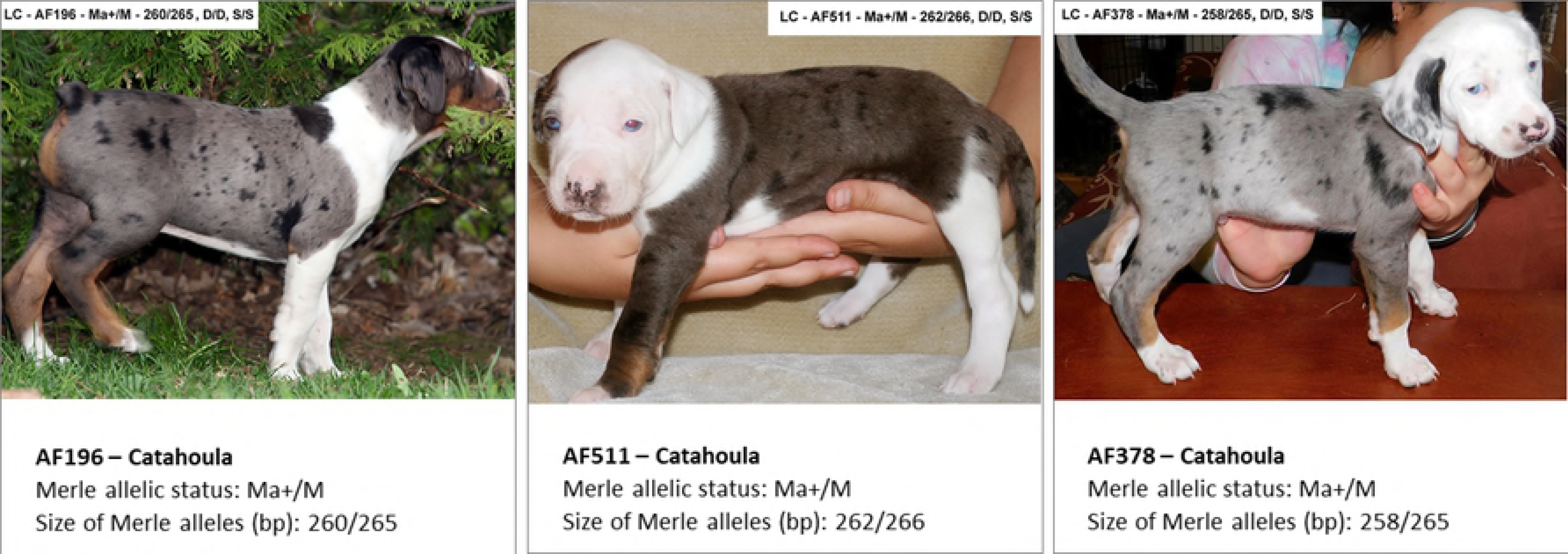
**Frequency of the Mh allele** within individual breeds, related to the occurence of the Mh allele in all dogs under study (*n* = 181). ASD – Australian Shepherd Dog (*n* = 40), AK – Australian Koolie (*n* = 23), BC – Border Collie (*n* = 18), D – Dachshund (*n* = 9), LC – Louisiana Catahoula (*n* = 73), MAUS – Miniature Australian Shepherd (*n* = 2), SSD – Shetland Sheepdog (*n* = 2), WS – Welsh Sheepdog (*n* = 3).

### Merle mosaicism

Mosaicism of the Merle gene seems to be quite wide-spread among the Merle breeds. It is characterized by a presence of more than two M Locus alleles in tested samples.

We have found that 16.6% of all dogs tested show Merle mosaicism, harboring 3 or more different M locus alleles in the tested sample. The ABI3500 semi-quantitive fragment analysis technology allows us to discriminate between a minor and major fraction of the summary peak signal for a given allele, and to use terms “minor” and “major” allele based on the height of the individual allelic peaks. Square brackets, [], are used to denote the minor allele/s.

Some mosaic results may explain an unusual phenotype that does not express as the two major alleles would be expected to. Fig 7 shows three examples documenting this phenomenon.

**Fig 7.**
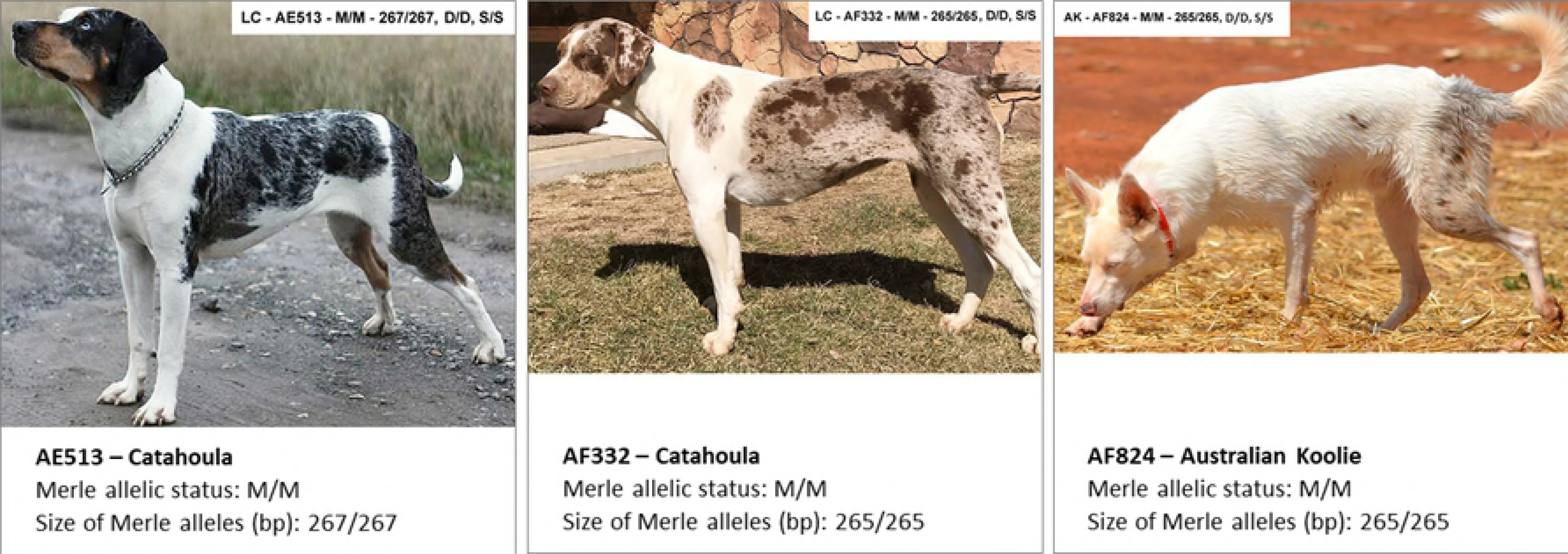
We have identified shortening of Merle in 27 dogs from our cohort of 181 animals tested (Table 3). We suggest that mosaicism represents the result of shorthening of major longer Merle alleles.

**Table 3:** Shortening of Merle alleles resulting in mosaicism.

| BREED | SAMPLE CODE | MERLE ALLELIC STATUS | MAJOR ALLELE/S | MINOR ALLELE/S | SHORTENED MERLE ALLELES |
| --- | --- | --- | --- | --- | --- |
| Catahoula | AE512 | [Mc]/[Mc+]/Ma/Mh<br>[218]/[238]/250/277 | 250/277 | 218/238 | Mh and/or Ma to Mc/Mc+ |
| Australian Koolie | AF521 | m/[Mc]/Mh<br>m/[225]/269 | 269 | 225 | Mh to Mc |
| Australian Shepherd | AF509 | [Mc]/[Mc]/ Mh/Mh<br>[220]/[230] /269/269 | 269/269 | 220/230 | Mh to Mc |
| Border Collie | AF459 | m/[Mc]/Mh<br>m/[225]/271 | 271 | 225 | Mh to Mc |
| Australian Shepherd | AE877 | m/[Mc]/Mh<br>m/[230]/270 | 270 | 230 | Mh to Mc |
| Australian Koolie | AF508 | [Mc]/[Mc]/[Mc]/[Mc+]/Mh/Mh<br>[213]/[220]/[230]/[235]/269/269 | 269/269 | 213/220/<br>230/235 | Mh to Mc and/or Mc+ |
| Shetland Sheepdog | AE982 | m/[Mc]/Mh<br>m/[220]/271 | 271 | 220 | Mh to Mc |
| Australian Koolie | AF513 | m/[Mc+]/Mh<br>m/[231]/278 | 278 | 231 | Mh to Mc+ |
| Australian Shepherd | AF074 | m/[Ma]/Mh<br>m/[254]/270 | 270 | 254 | Mh to Ma |
| Miniature Australian Shepherd | AE820 | m/[Ma+]/Mh<br>m/[262]/273 | 273 | 262 | Mh to Ma+ |
| Catahoula | AE870 | m/[Mc]/M<br>m/[208]/265 | 265 | 208 | M to Mc |
| French Bulldog | AE941 | m/[Mc+]/M<br>m/[241]/265 | 265 | 241 | M to Mc+ |
| Catahoula | AE787 | [Ma]/M/M<br>[251]/265/265 | 265/265 | 251 | M to Ma |
| Catahoula | AF132 | m/[Ma+]/M<br>m/[257]/266 | 266 | 257 | M to Ma+ |
| Border Collie | AF163 | m/[Ma+]/M<br>m/[260]/268 | 268 | 260 | M to Ma+ |
| Dachshund | AF020 | m/[Ma+]/M<br>m/[258]/268 | 268 | 258 | M to Ma+ |
| Border Collie | AF163 | m/[Ma+]/M<br>m/[260]/268 | 268 | 260 | M to Ma+ |
| Catahoula | AE956 | [Mc]/Ma/M<br>[225]/251/265 | 251/265 | 225 | M and/or Ma to Mc |
| Catahoula | AE515 | [Mc]/Ma/M<br>[221]/252/267 | 252/267 | 221 | M or Ma to Mc |
| Border Collie | AF514 | m/[Mc]/[Mc+]/Ma+<br>m/[222]/[246]/262 | 262 | 222/246 | Ma+ to Mc+ and/or Mc |
| Australian Koolie | AF225 | m/[Mc]/Ma+<br>m/[215]/262 | 262 | 215 | Ma+ to Mc |
| Australian Shepherd | AE786 | m/[Mc+]/Ma<br>m/[243]/252 | 252 | 243 | Ma to Mc+ |
| Catahoula | AF265 | m/[Mc]/Mc+<br>m/[222]/239 | 239 | 222 | Mc+ to Mc |
| Wesh Sheepdog | AF613 | m/[Mc+]/Mh<br>m/[231]/273 | 273 | 231 | Mh to Mc+ |
| Australian Shepherd | AF652 | m/[Mc]/M<br>m/[226]/266 | 266 | 226 | M to Mc |
| Border Collie | AF781 | m/[Ma+]/M<br>m/[261]/267 | 267 | 261 | M to Ma+ |
| Border Collie | AF782 | m [Ma+]/Mh<br>m [257] 270 | 270 | 257 | Mh to Ma+ |
| Mudi | AF650 | m/[Mc]/M<br>m/[226]/267 | 267 | 226 | M to Mc |

Interestingly, in three examples (AF038, AE943 and AF273) we observed that the longer Merle allele represented the minor allelic population, with the shorter allelic variant being the major allele. Nevertheless, we consider this more of an artifact caused by suboptimal quality of isolated DNA (random PCR bias leading to the amplification of shorter Merle fragment) rather than a real finding.

To test the mode of inheritance of these multiple allelic variants, we have tested STR – proven pedigrees and have shown that mosaic alleles presented in the area of reproduction of egg and sperm from the parents can be distributed towards the offspring and can be identified based on their size. Fig 8 shows a typical pedigree exhibiting this phenomenon.

**Fig 8.**
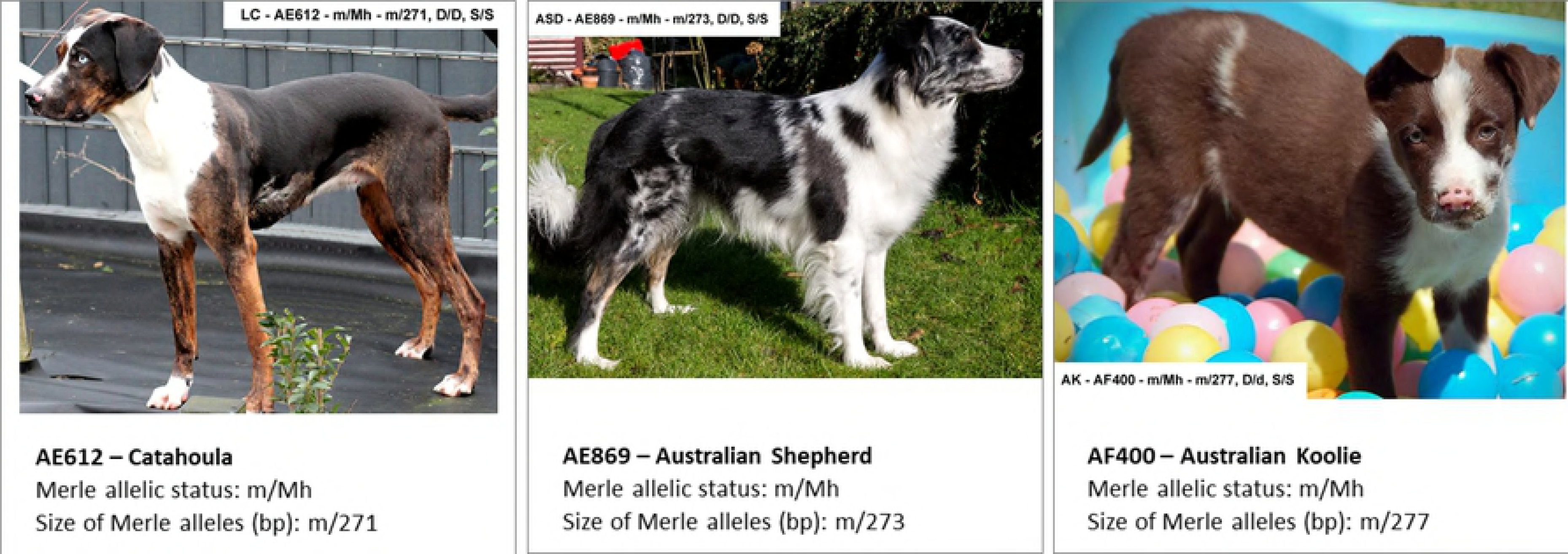
Typical pedigree of offspring carrying minor mosaic alleles.

Interestingly, the Merle allelic status of mosaic animals can differ between biological materials tested, and, as shown in Fig 9, the findings in buccal swab (terminally differentiated epidermal derivative) can dramatically difer from the finding in sperm cells (germinal cell line). In this specific example, the proband, sire AF174, has been routinely tested using buccal swab and found out to carry [Mc]/Mh/Mh; ([212]/269/269), with the Mc allele representing the minor fraction. The dam, AF484, was genotyped as m/Mc+; (m/231). Rather unexpectedly, their offspring, AF327, exhibited the following Merle genetic status: m/Mc+; (m/235), with the 235 allele seemingly not to originate from either of the alleged parents. To exclude any parental mismatch, paternity has been confirmed using STR genotyping. Thus, in searching for the source of the Mc+ allele of 235 bp, Merle genotyping of the sperm cells of the sire, AF174, was performed. Rather interestingly, the Merle allelic genotype of the semen revealed a mosaicism of [Mc]/[Mc]/[Mc]/Mc+]/Mh/Mh - ([213]/[220]/[230]/[235]/269/269). From this extensive Merle allelic pool identified in the sperm cells of the sire, the searched for Mc+ allele of 235 bp, has been passed to the offspring AF327.

Together with the phenomenon of consistent shortening of major Merle alleles in Merle mosaic animals observed in 27 out of 181 animals in our cohort (Table 3), the finding of semen as a source of a significant pool of minor Merle alleles, has led us to a hypothesis we herein propose – unexpected Merle genotypes found in the offspring, not corresponding to the Merle genotypes obtained on other than germline cells in the parents, can be explained by the hidden pool of Merle alleles in the germline, undetectable in buccal swab or other differentiated tissues, rather than by the original theory of prolongation of shortened Merle alleles.

**Fig 9.**
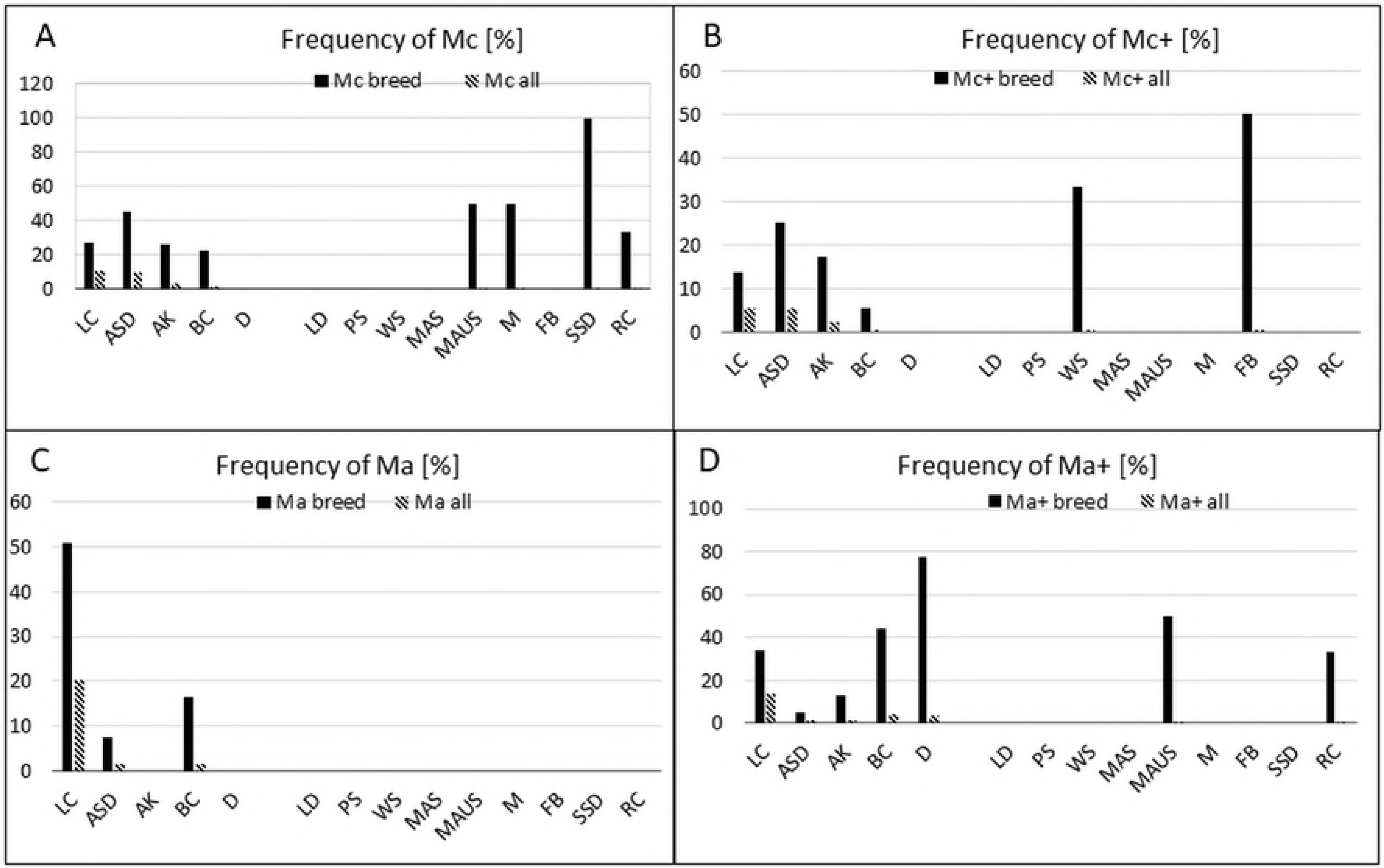
Different genotypes found in buccal swab (terminally differentiated epidermal derivative) and sperm cells (germinal cell line).

As discussed above, Merle allelic status might differ dramatically among the biological materials investigated. The following Fig 10 shows the difference in Merle genotypes in the buccal swab (**A**) m/[Mc]/Mh – (m/[230]/270), with the Mc allele being the minor one, and the hair (**B**) Mh/Mh – (271/271) of a proband AE877. Since the hair was sampled from the Merle-exhibiting areas, the genotype Mh/Mh could have been expected. Still, the buccal swab has shown a wider portfolio of Merle alleles - m/[Mc]/Mh. Given the fact that the proband, AE877, is a female, it was not possible to analyze the Merle allelic status of her germline – oocytes cannot be obtained non-invasively and we respect the animals.

**Fig 10.**
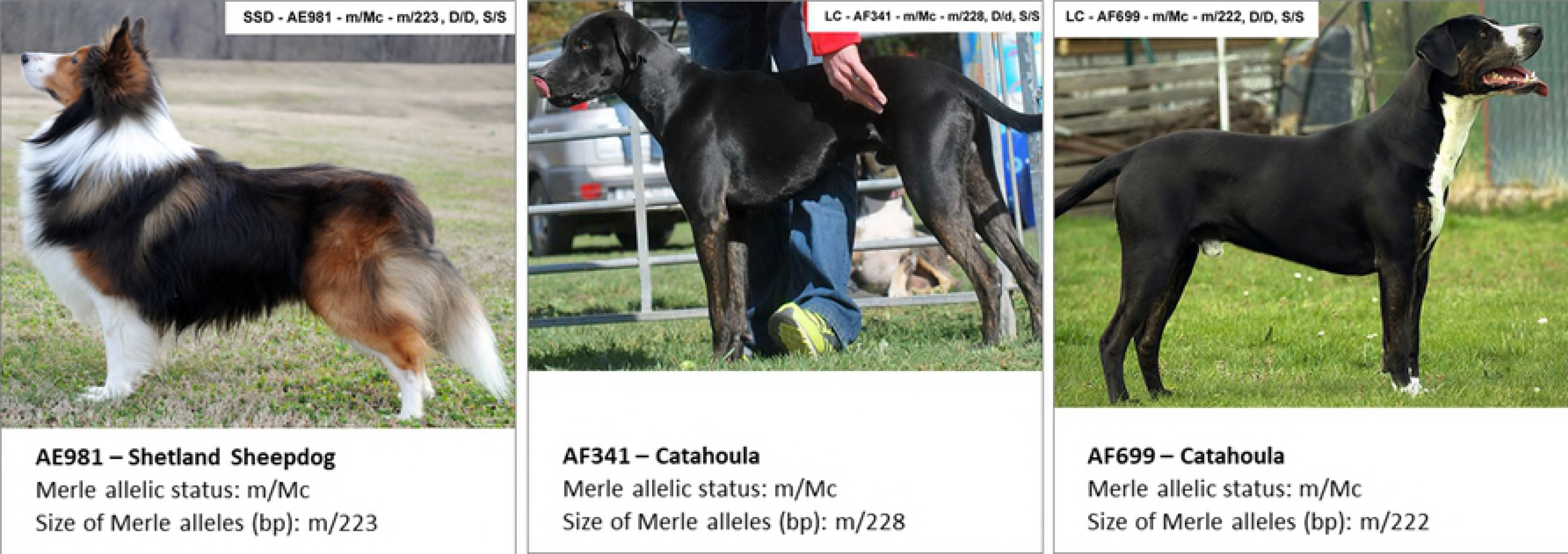
Difference in Merle allelic status between biological materials tested, proband AE877, a female ASD.

Fig 11 shows another example of the same theme - in proband AF132, the routine testing of peripheral blood (**A**) has shown a significantly different Merle profile when compared to hair from Merle patches, and sperm cells. Similar to the case depicted in Fig 10, the reason for semen testing in proband AF132 was the offspring that exhibited Merle allelic status incompatible with the obligatory parental alleles identified using buccal swab in both sire and dam. For more details, see also the parentage Merle details of the proband AF132 given in Fig 8.

**Fig 11.**
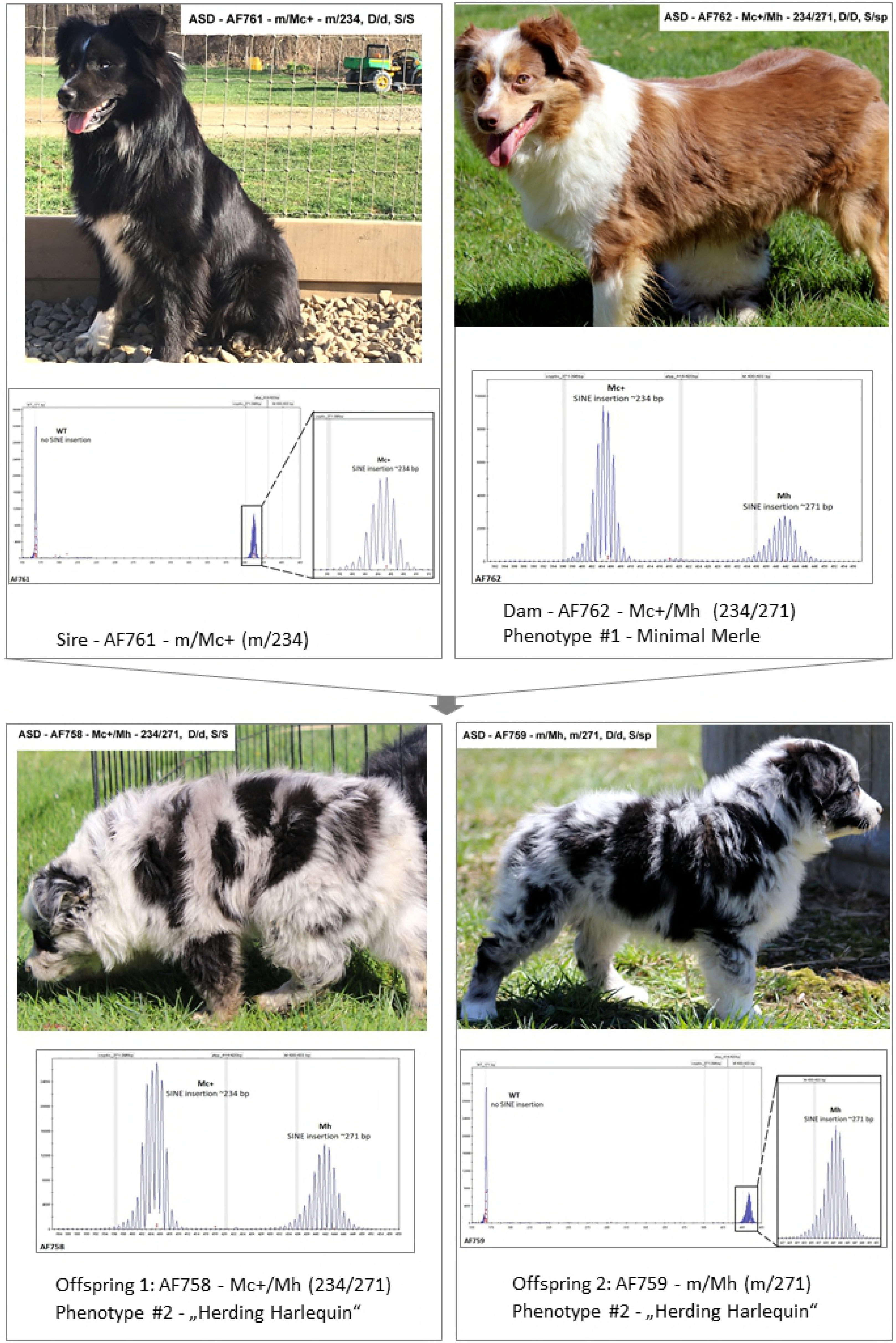
Difference in Merle allelic status among biological materials tested, proband AF132, a male LC.

**Table 4.** shows the details of Merle allelic mosaicism in 30 individuals.

| Sample code | Allelic status of Merle alleles | Size of Merle alleles in bp | Breed | Biological material | SuperColorLocus reference (Table 5) |
| --- | --- | --- | --- | --- | --- |
| AE512 | [Mc]/[Mc+]/Ma/Mh | [218]/[238]/250/277 | Catahoula | buccal swab |  |
| AE956 | [Mc]/Ma/M | [225]/251/265 | Catahoula | buccal swab |  |
| AE787 | [Ma]/M/M | [251]/265/265 | Catahoula | buccal swab |  |
| AE515 | [Mc]/Ma/M | [221]/252/267 | Catahoula | peripheral blood |  |
| AE786 | m/[Mc+]/Ma | m/[243]/252 | Australian Shepherd | hair |  |
| AE820 | m/[Ma+]/Mh | m/[262]/273 | Miniature Australian Shepherd | peripheral blood |  |
| AE982 | m/[Mc]/Mh | m/[220]/271 | Shetland Sheepdog | buccal swab |  |
| AF074 | m/[Ma]/Mh | m/[254]/270 | Australian Shepherd | buccal swab |  |
| AF104 | m/[Mc+]/M | m/[241]/265 | French Bulldog | hair |  |
| AF163 | m/[Ma+]/M | m/[260]/268 | Border Collie | buccal swab |  |
| AF038 | m/Ma+/[M] | m/258/[266] | Catahoula | buccal swab |  |
| AF508 | [Mc]/[Mc]/[Mc]/Mc+/Mh/Mh | [213]/[220]/[230]/[235]/269/269 | Australian Koolie | semen | AF174 - buccal swab |
| AF225 | m/[Mc]/Ma+ | m/[215]/262 | Australian Koolie | buccal swab |  |
| AE132 | m/[Ma+]/M | m/[257]/266 | Catahoula | hair | AE802 - buccal swab |
| AE870 | m/[Mc]/M | m/[208]/265 | Catahoula | peripheral blood |  |
| AF265 | m/[Mc]/Mc+ | m/[222]/239 | Catahoula | hair |  |
| AF020 | m/[Ma+]/M | m/[258]/268 | Dachshund | buccal swab |  |
| AE943 | m/Mc+/[M] | m/233/[267] | Australian Shepherd | semen |  |
| AE877 | m/[Mc]/Mh | m/[230]/270 | Australian Shepherd | buccal swab |  |
| AF273 | m/Ma/[M] | m/250/[266] | Catahoula | peripheral blood |  |
| AF459 | m/[Mc]/Mh | m/[225]/271 | Border Collie | buccal swab |  |
| AF513 | m/[Mc+]/Mh | m/[231]/278 | Australian Koolie | hair |  |
| AF514 | m/[Mc]/[Mc+]/Ma+ | m/[222]/[246]/262 | Border Collie | buccal swab |  |
| AF509 | [Mc]/[Mc]/ Mh/Mh | [220]/[230] /269 269 | Australian Shepherd | hair |  |
| AF521 | m/[Mc]/Mh | m/[225]/269 | Australian Koolie | buccal swab |  |
| AF613 | m/[Mc+]/Mh | m/[231]/273 | Welsh Sheepdog | hair |  |
| AF652 | m/[Mc]/ M | m/[226]/266 | Australian Shepherd | hair |  |
| AF781 | m/[Ma+]/M | m/[261]/267 | Border Collie | buccal swab |  |
| AF782 | m/[Ma+]/Mh | m/[257]/270 | Border Collie | buccal swab |  |
| AF650 | m/[Mc]/M | m/[226]/267 | Mudi | buccal swab |  |

**Fig 12.**
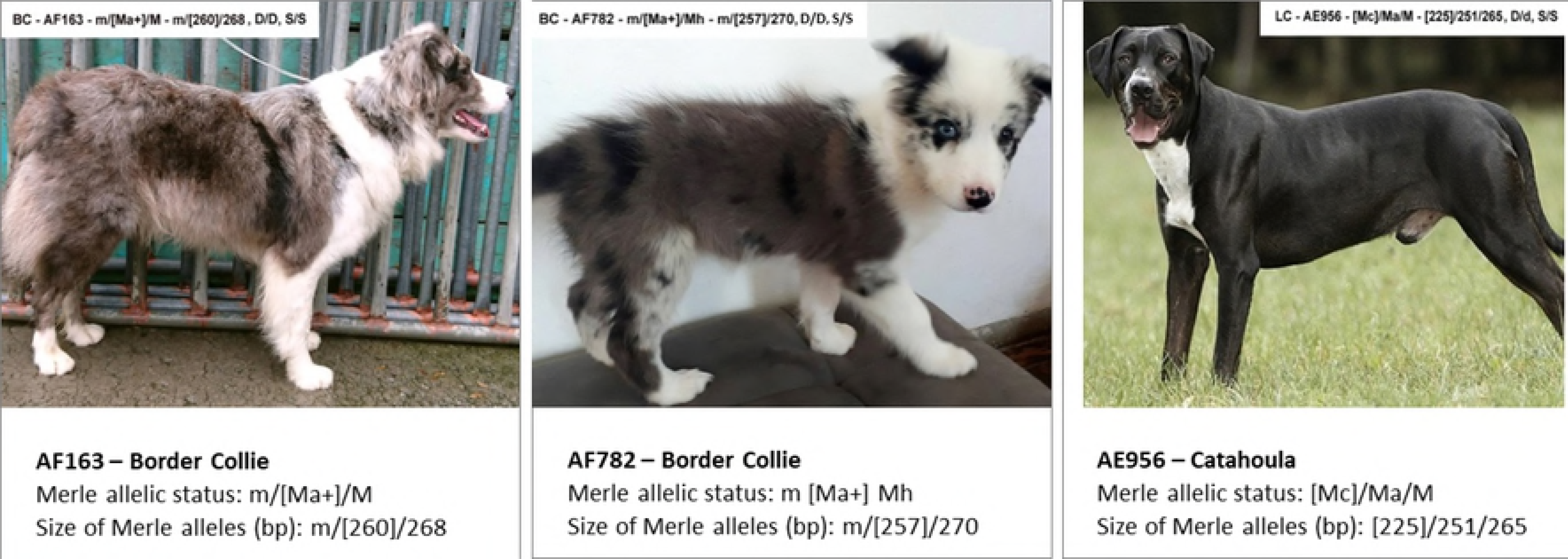
shows photographs and Merle chromatograms of the individuals given in Table 4.

Mosaicism of the Merle gene was observed in 10 breeds out of 14 tested (Fig 13). No mosaicism was found in LD, MAS, and RC, whereas its frequency was approximately 50% in other breeds with a low number of tested dogs (WS, MAUS, M, FB, and SSD). The mosaicism was the most widespread in BC (27.8% of tested individuals), followed by AK, ASD, LC, and D with 17.4,15, 12.3, and 11% of tested dogs. Moreover, BC, AK, and ASD (and also WS, MAUS, and SSD) also are the breeds where the Mh allele occur more frequently, contrary to LC and D (Fig 6).

**Fig 13:**
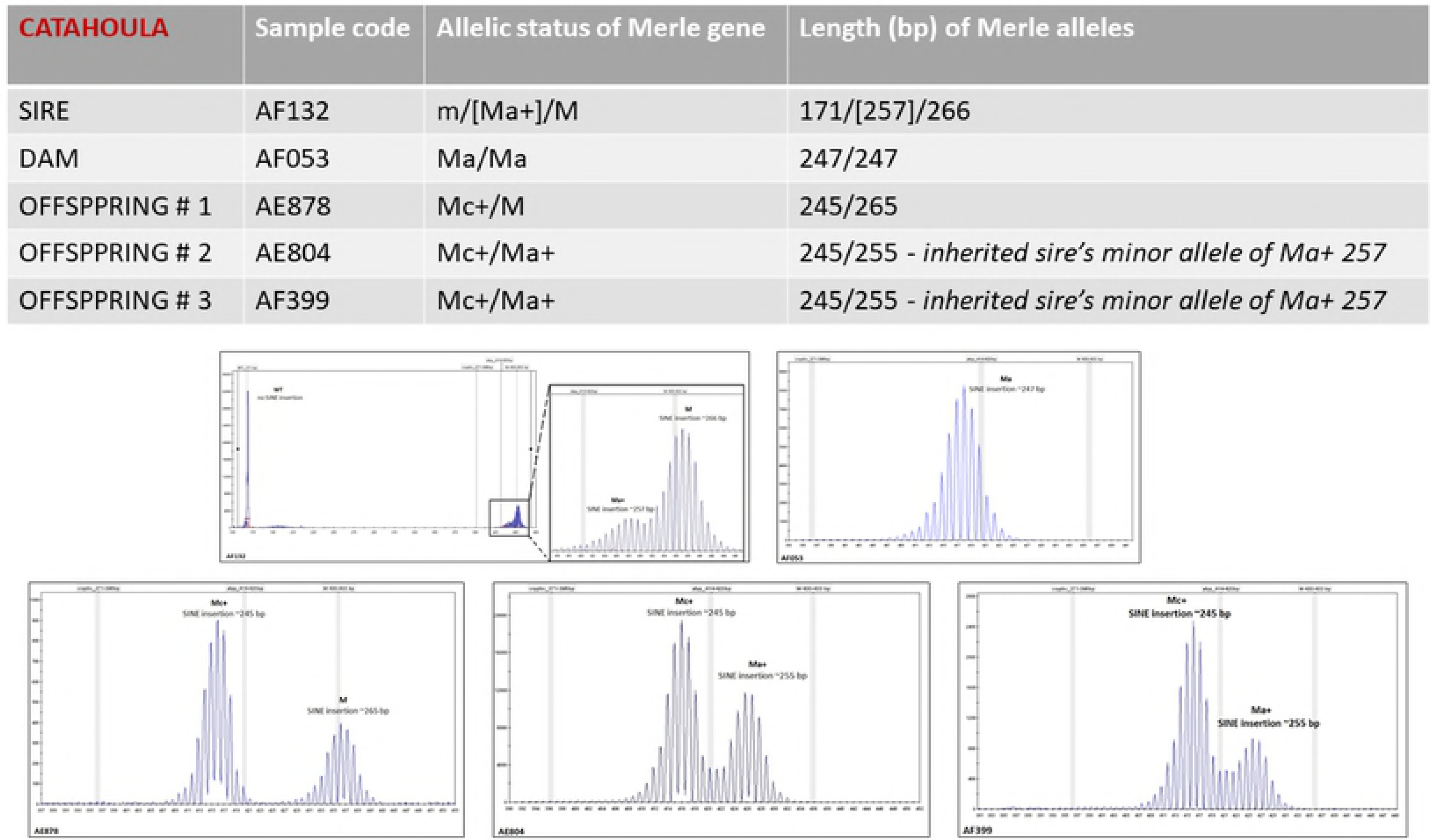
Occurrence of mosaic in different breeds. Values represent the percentage of total number of dogs in each breed with Merle mosaicism detected [ASD – Australian Shepherd Dog (*n* = 40), AK – Australian Koolie (*n* = 23), BC – Border Collie (*n* = 18), D – Dachshund (*n* = 9), LC – Louisiana Catahoula (*n* = 73), MAUS – Miniature Australian Shepherd (*n* = 2), SSD – Shetland Sheepdog (*n* = 2), FB – French Bulldog (*n* = 2), WS – Welsh Sheepdog (*n* = 3), M – Mudi (*n* = 2)] or all tested dogs (*n* = 181).

The finding of Merle mosaicism can further be supported by significant differences between the occurence of major or minor alleles found in the animals tested (Fig 14). The major alleles found in our study included m (30%), M (15%) and Mh (15%), followed by Mc, Ma and Ma+ alleles in a few cases. In contrast, Mc (50%), Mc+ (19.4%), and Ma+ (16.7%) represented the most abundant minor alleles.

**Fig 14:**
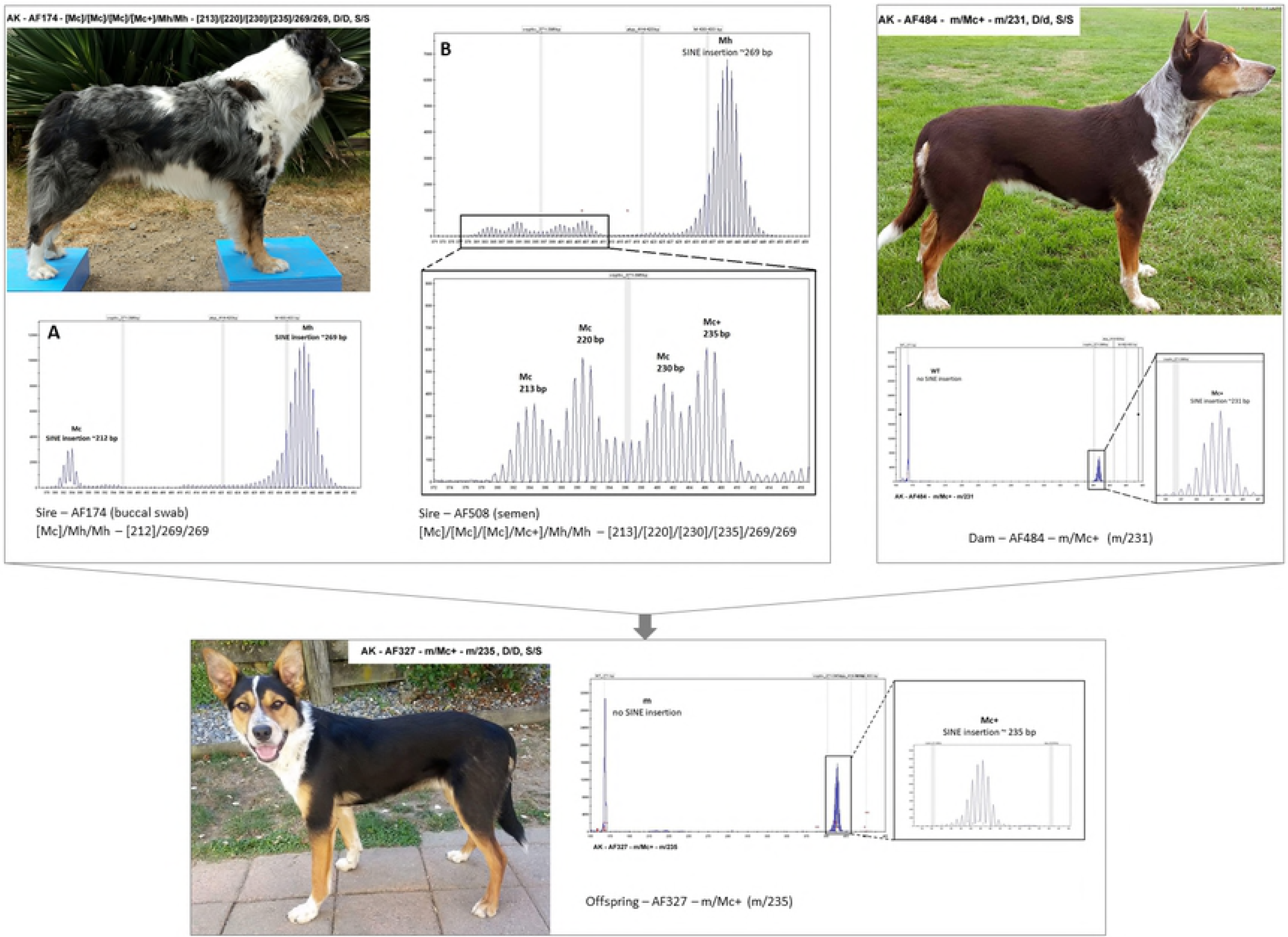
Relative frequency of individual alleles as percentage of all alleles found as major or minor ones in mosaic dogs, calculated for all breeds together.

### SuperColorLocus testing and Merle – phenotypic relationships

Using the SuperColorLocus analysis we tested our cohort of dogs also for the major coat color loci (locus E, K, A) and their modifiers (locus B, D, S) to explain phenotypic features of the dogs in the study and also to evaluate a possible relation to the phenotype of dogs carrying Merle alleles. Table 5 **(The SuperColorLocus table)** shows the coat color genotypes of all dogs in our study; all respective photographs of the animals can be found in Fig S3A-S3X in the Supplement section. We also evaluated the influence of S and D loci on the final phenotype of the Merle carrying dogs and found that the S locus does not seem to be a strong modifier of the resulting Merle phenotype. S locus genotype also seems to have minimal, if any, impact on health consequences connected with Merle phenomenon. Genotyping of the other coat color modifying loci – E, K, A, B and D has shown an expected correlation with the actual phenotypes of the animals – for comparison of coat color genotypes and phenotypes please refer to Table 5 and Figs S3A-S3X.

**Table 5.** shows the genotype summary of the so far recognized coat color loci and their modifiers relevant for the study.

SuperColorLocus – GENOTYPE SUMMARY
| Sample code | Breed | Locus E<br>(Alleles E, e) | Locus E<br>(Allele Em) | Locus E<br>(allele Eg) | Locus E<br>(allele Eh) | Locus K<br>(alleles K, k*) | Locus A<br>(alleles Ay, aw, at, a) | Locus B<br>(allele bc) | Locus B<br>(allele bs) | Locus B<br>(allele bd) | Locus D | Locus S | Locus H<br>(Harlequin - PSMB7) |
| --- | --- | --- | --- | --- | --- | --- | --- | --- | --- | --- | --- | --- | --- |
| AE428 | Catahoula | E/E | wt/wt | wt/wt | wt/wt | K/k* | at/at | B/B | B/bs | B/B | D/D | S/S | h/h |
| AE463 | Catahoula | E/E | Em/Em | wt/wt | wt/wt | K/k* | at/at | B/B | B/B | B/B | D/d | S/S | h/h |
| AE479 | Catahoula | E/e | wt/wt | wt/wt | wt/wt | K/k* | at/at | B/B | B/B | B/B | D/D | S/S | h/h |
| AE487 | Catahoula | E/E | Em/wt | wt/wt | wt/wt | K/k* | Ay/aw | B/B | B/B | B/B | D/D | S/S | h/h |
| AE488 | Catahoula | E/E | wt/wt | wt/wt | wt/wt | K/k* | at/at | B/B | B/bs | B/B | D/d | S/sp | h/h |
| AE504 | Catahoula | E/e | wt/wt | wt/wt | wt/wt | k*/k* | at/a | B/B | B/bs | B/B | D/D | S/S | h/h |
| AE505 | Catahoula | E/E | Em/wt | wt/wt | wt/wt | k*/k* | at/at | B/B | B/B | B/B | D/D | S/S | h/h |
| AE512 | Catahoula | E/E | Em/Em | wt/wt | wt/wt | K/k* | at/a | B/B | B/B | B/B | D/D | S/S | h/h |
| AE513 | Catahoula | E/E | wt/wt | wt/wt | wt/wt | k*/k* | at/at | B/B | B/B | B/B | D/D | S/S | h/h |
| AE514 | Catahoula | E/E | wt/wt | wt/wt | wt/wt | k*/k* | at/at | B/B | bs/bs | B/B | D/d | S/S | h/h |
| AE515 | Catahoula | E/E | Em/wt | wt/wt | wt/wt | K/k* | at/at | B/B | bs/bs | B/B | D/D | S/sp | h/h |
| AE535 | Catahoula | E/E | Em/Em | wt/wt | wt/wt | K/k* | at/at | B/B | B/B | B/B | D/D | S/S | h/h |
| AE572 | Catahoula | E/E | wt/wt | wt/wt | wt/wt | k*/k* | at/at | B/B | bs/bs | B/B | D/D | S/S | h/h |
| AE573 | Catahoula | E/E | wt/wt | wt/wt | wt/wt | K/k* | at/at | B/B | B/B | B/B | D/d | S/S | h/h |
| AE574 | Catahoula | E/E | Em/wt | wt/wt | wt/wt | K/k* | at/at | B/bc | B/bs | B/bd | D/d | S/S | h/h |
| AE612 | Catahoula | E/E | Em/wt | wt/wt | wt/wt | K/k* | at/a | B/B | B/bs | B/B | D/D | S/S | h/h |
| AE621 | Catahoula | E/E | Em/Em | wt/wt | wt/wt | K/k* | at/at | B/B | B/bs | B/B | D/D | S/S | h/h |
| AE622 | Catahoula | E/E | Em/wt | wt/wt | wt/wt | K/k* | at/at | B/B | B/bs | B/B | D/D | S/S | h/h |
| AE693 | Catahoula | E/E | wt/wt | wt/wt | wt/wt | K/k* | at/at | B/B | bs/bs | B/B | D/d | sp/sp | h/h |
| AE740 | Catahoula | E/E | Em/Em | wt/wt | wt/wt | K/k* | at/at | B/B | bs/bs | B/B | D/d | S/S | h/h |
| AE786 | Australian Shepherd | E/E | Em/wt | wt/wt | wt/wt | k*/k* | at/at | B/B | B/bs | B/B | D/D | S/S | h/h |
| AE787 | Catahoula | E/E | Em/wt | wt/wt | wt/wt | K/k* | at/at | B/bc | B/bs | B/bd | D/D | S/sp | h/h |
| AE802 | Catahoula | E/E | Em/Em | wt/wt | wt/wt | k*/k* | at/at | B/B | bs/bs | B/B | D/D | S/sp | h/h |
| AE803 | Australian Shepherd | E/E | wt/wt | wt/wt | wt/wt | k*/k* | at/at | B/B | B/bs | B/B | D/D | S/S | h/h |
| AE804 | Catahoula | E/E | Em/wt | wt/wt | wt/wt | K/k* | at/at | B/B | B/bs | B/B | D/D | S/sp | h/h |
| AE817 | Border Collie | E/e | Em/wt | wt/wt | wt/wt | K/K | aw/at | B/B | B/B | B/B | D/d | S/S | h/h |
| AE818 | Border Collie | E/E | Em/wt | wt/wt | wt/wt | K/k* | at/at | B/B | B/bs | B/B | D/d | S/sp | h/h |
| AE819 | Border Collie | E/E | Em/Em | wt/wt | wt/wt | K/k* | at/at | B/B | B/B | B/B | D/d | S/S | h/h |
| AE820 | Miniature Australian Shepherd | E/E | wt/wt | wt/wt | wt/wt | k*/k* | at/a | B/B | B/B | B/B | D/D | S/S | h/h |
| AE845 | Miniature American Shepherd | E/e | wt/wt | wt/wt | wt/wt | k*/k* | at/a | B/bc | B/bs | B/bd | D/D | S/S | h/h |
| AE846 | Miniature American Shepherd | E/E | wt/wt | wt/wt | wt/wt | k*/k* | at/at | B/B | B/bs | B/B | D/D | S/S | h/h |
| AE852 | Catahoula | E/E | Em/Em | wt/wt | wt/wt | K/k* | at/a | B/B | B/bs | B/B | D/D | S/S | h/h |
| AE868 | Australian Shepherd | E/E | wt/wt | wt/wt | wt/wt | k*/k* | at/at | B/B | bs/bs | B/B | D/D | S/S | h/h |
| AE869 | Australian Shepherd | E/E | wt/wt | wt/wt | wt/wt | k*/k* | at/at | B/B | B/bs | B/B | D/D | S/S | h/h |
| AE870 | Catahoula | E/E | Em/wt | wt/wt | wt/wt | K/k* | at/a | B/B | bs/bs | B/B | D/D | S/S | h/h |
| AE877 | Australian Shepherd | E/E | wt/wt | wt/wt | wt/wt | k*/k* | at/at | B/B | B/bs | B/B | D/D | S/S | h/h |
| AE878 | Catahoula | E/E | Em/wt | wt/wt | wt/wt | K/k* | at/at | B/B | bs/bs | B/B | D/D | S/S | h/h |
| AE903 | Australian Shepherd | E/E | wt/wt | wt/wt | wt/wt | k*/k* | at/at | B/B | bs/bs | B/B | D/D | S/S | h/h |
| AE904 | Catahoula | E/E | Em/Em | wt/wt | wt/wt | K/k* | at/a | B/B | B/bs | B/B | D/D | S/S | h/h |
| AE905 | Catahoula | E/E | Em/wt | wt/wt | wt/wt | k*/k* | at/at | B/B | B/bs | B/B | D/D | S/S | h/h |
| AE939 | Australian Shepherd | E/E | wt/wt | wt/wt | wt/wt | k*/k* | at/at | B/B | B/bs | B/B | D/D | S/S | h/h |
| AE941 | French Bulldog | E/E | Em/wt | wt/wt | wt/wt | K/k* | Ay/a | B/B | B/B | B/B | D/d | S/S | h/h |
| AE942 | French Bulldog | E/E | Em/Em | wt/wt | wt/wt | K/k* | Ay/a | B/B | B/B | B/B | d/d | S/sp | h/h |
| AE943 | Australian Shepherd | E/e | Em/wt | wt/wt | wt/wt | k*/k* | at/at | B/B | bs/bs | B/B | D/D | S/S | h/h |
| AE956 | Catahoula | E/E | Em/wt | wt/wt | wt/wt | K/k* | at/at | B/B | B/B | B/B | D/d | S/S | h/h |
| AE957 | Australian Shepherd | E/E | Em/wt | wt/wt | wt/wt | k*/k* | at/at | B/B | B/bs | B/B | D/D | S/S | h/h |
| AE981 | Shetland Sheepdog | E/E | wt/wt | wt/wt | wt/wt | k*/k* | Ay/aw | B/B | B/B | B/B | D/D | S/S | h/h |
| AE982 | Shetland Sheepdog | E/E | wt/wt | wt/wt | wt/wt | k*/k* | at/a | B/B | B/B | B/B | D/D | S/S | h/h |
| AF008 | Australian Shepherd | E/E | wt/wt | wt/wt | wt/wt | k*/k* | at/at | B/bc | B/B | B/bd | D/D | S/S | h/h |
| AF009 | Australian Shepherd | E/E | wt/wt | wt/wt | wt/wt | k*/k* | at/at | B/bc | B/B | B/bd | D/d | S/S | h/h |
| AF010 | Australian Shepherd | E/E | wt/wt | wt/wt | wt/wt | k*/k* | at/at | B/bc | B/B | B/bd | D/d | S/S | h/h |
| AF011 | Australian Shepherd | E/E | Em/wt | wt/wt | wt/wt | k*/k* | at/at | B/bc | B/B | B/bd | D/D | S/S | h/h |
| AF012 | Australian Shepherd | E/E | wt/wt | wt/wt | wt/wt | k*/k* | at/at | B/bc | B/B | B/bd | D/D | S/S | h/h |
| AF013 | Australian Shepherd | E/E | wt/wt | wt/wt | wt/wt | k*/k* | at/at | bc/bc | B/B | B/bd | D/d | S/S | h/h |
| AF014 | Australian Shepherd | E/E | Em/wt | wt/wt | wt/wt | k*/k* | at/at | B/B | B/B | B/B | D/D | S/S | h/h |
| AF015 | Australian Shepherd | E/E | Em/wt | wt/wt | wt/wt | k*/k* | at/at | B/bc | B/B | B/bd | D/D | S/S | h/h |
| AF016 | Dachshund | E/E | wt/wt | wt/wt | wt/wt | k*/k* | at/at | B/B | B/B | B/B | D/D | S/S | h/h |
| AF017 | Dachshund | E/E | wt/wt | wt/wt | wt/wt | k*/k* | at/at | bc/bc | B/B | B/B | D/d | S/S | h/h |
| AF018 | Dachshund | E/E | wt/wt | wt/wt | wt/wt | k*/k* | at/at | B/B | B/B | B/B | D/D | S/S | h/h |
| AF019 | Dachshund | E/E | wt/wt | wt/wt | wt/wt | k*/k* | at/at | B/bc | B/B | B/B | D/D | S/S | h/h |
| AF020 | Dachshund | E/E | wt/wt | wt/wt | wt/wt | k*/k* | at/at | B/bc | B/B | B/B | D/D | S/S | h/h |
| AF022 | Dachshund | E/E | wt/wt | wt/wt | wt/wt | k*/k* | at/at | B/bc | B/B | B/B | D/d | S/S | h/h |
| AF023 | Dachshund | E/E | wt/wt | wt/wt | wt/wt | k*/k* | at/at | B/bc | B/B | B/B | D/D | S/S | h/h |
| AF038 | Catahoula | E/E | Em/Em | wt/wt | wt/wt | K/k* | aw/at | B/B | B/B | B/B | D/D | S/S | h/h |
| AF039 | Catahoula | E/E | Em/Em | wt/wt | wt/wt | K/k* | Ay/a | B/B | B/B | B/B | D/D | S/S | h/h |
| AF050 | Australian Shepherd | E/E | Em/wt | wt/wt | wt/wt | k*/k* | at/at | B/bc | B/B | B/bd | D/D | S/sp | h/h |
| AF052 | Australian Shepherd | E/E | wt/wt | wt/wt | wt/wt | k*/k* | at/at | B/bc | B/B | B/bd | D/D | S/sp | h/h |
| AF053 | Catahoula | E/E | wt/wt | wt/wt | wt/wt | K/k* | aw/at | B/B | B/bs | B/B | D/D | S/S | h/h |
| AF056 | Border Collie | E/E | wt/wt | wt/wt | wt/wt | k*/k* | at/at | B/B | B/B | B/B | D/d | S/S | h/h |
| AF072 | Catahoula | E/E | Em/w<br>t | wt/wt | wt/wt | k*/k* | at/at | B/B | B/bs | B/B | D/d | S/S | h/h |
| AF073 | Rough Collie | E/E | wt/wt | wt/wt | wt/wt | k*/k* | at/at | B/B | B/B | B/B | D/D | S/S | h/h |
| AF074 | Australian Shepherd | E/E | wt/wt | wt/wt | wt/wt | K/k* | at/at | B/B | B/bs | B/B | D/D | S/S | h/h |
| AF096 | Australian Shepherd | E/E | wt/wt | wt/wt | wt/wt | k*/k* | at/at | B/B | B/bs | B/B | D/D | S/sp | h/h |
| AF099 | Catahoula | E/E | Em/E<br>m | wt/wt | wt/wt | k*/k* | Ay/at | B/B | B/bs | B/B | D/D | S/S | h/h |
| AF101 | Catahoula | E/E | Em/E<br>m | wt/wt | wt/wt | K/k* | at/at | B/B | B/bs | B/B | D/D | S/S | h/h |
| AF104 | French Bulldog | E/E | Em/w<br>t | wt/wt | wt/wt | K/k* | Ay/a | B/B | B/B | B/B | D/d | S/S | h/h |
| AF129 | Catahoula | E/E | Em/E<br>m | wt/wt | wt/wt | K/k* | at/at | B/B | B/bs | B/B | D/D | S/sp | h/h |
| AF133 | Catahoula | E/E | Em/E<br>m | wt/wt | wt/wt | K/k* | at/at | B/B | B/bs | B/B | D/D | S/S | h/h |
| AF134 | Border Collie | E/e | Em/w<br>t | wt/wt | wt/wt | K/K | aw/at | B/B | B/bs | B/B | D/D | S/S | h/h |
| AF135 | Dachshund | E/E | wt/wt | wt/wt | wt/wt | k*/k* | Ay/at | B/B | B/B | B/B | D/D | S/sp | h/h |
| AF143 | Australian Koolie | E/E | Em/E<br>m | wt/wt | wt/wt | K/k* | aw/at | B/B | bs/bs | B/B | D/D | S/S | h/h |
| AF146 | Catahoula | E/E | Em/w<br>t | wt/wt | wt/wt | K/k* | at/at | B/B | B/bs | B/B | D/D | S/sp | h/h |
| AF161 | Catahoula | E/E | Em/E<br>m | wt/wt | wt/wt | K/k* | at/at | B/B | bs/bs | B/B | D/d | S/sp | h/h |
| AF163 | Border Collie | E/E | Em/w<br>t | wt/wt | wt/wt | K/k* | at/at | B/B | B/B | B/B | D/D | S/S | h/h |
| AF174 | Koolie | E/E | Em/w<br>t | wt/wt | wt/wt | K/k* | at/at | B/B | B/B | B/B | D/D | S/S | h/h |
| AF176 | Catahoula | E/E | Em/E<br>m | wt/wt | wt/wt | K/k* | at/at | B/B | B/B | B/B | D/D | S/S | h/h |
| AF185 | Catahoula | E/E | Em/E<br>m | wt/wt | wt/wt | k*/k* | at/at | B/B | B/B | B/B | D/D | S/S | h/h |
| AF195 | Rough Collie | E/E | wt/wt | wt/wt | wt/wt | k*/k* | at/at | B/B | B/B | B/B | D/D | S/sp | h/h |
| AF196 | Catahoula | E/E | Em/w<br>t | wt/wt | wt/wt | k*/k* | at/at | B/B | B/bs | B/B | D/D | S/S | h/h |
| AF198 | Catahoula | E/E | Em/w<br>t | wt/wt | wt/wt | k*/k* | at/at | B/B | bs/bs | B/B | D/D | S/S | h/h |
| AF223 | Australian Koolie | E/E | wt/wt | wt/wt | wt/wt | k*/k* | at/at | B/B | bs/bs | B/B | D/d | S/S | h/h |
| AF225 | Australian Koolie | E/E | wt/wt | wt/wt | wt/wt | K/K | at/at | B/B | B/bs | B/B | D/D | S/S | h/h |
| AF227 | Australian Koolie | E/E | Em/w<br>t | wt/wt | wt/wt | k*/k* | at/at | B/B | B/bs | B/B | D/D | S/S | h/h |
| AF231 | Catahoula | E/e | Em/w<br>t | wt/wt | wt/wt | k*/k* | at/a | B/B | B/bs | B/B | D/D | S/S | h/h |
| AF232 | Catahoula | E/E | Em/E<br>m | wt/wt | wt/wt | k*/k* | at/at | B/B | B/B | B/B | D/D | S/S | h/h |
| AF237 | Australian Shepherd | E/E | Em/w<br>t | wt/wt | wt/wt | k*/k* | at/at | B/bc | B/B | B/bd | D/D | S/S | h/h |
| AF239 | Australian Shepherd | E/E | Em/w<br>t | wt/wt | wt/wt | k*/k* | at/at | B/B | B/bs | B/B | D/D | S/S | h/h |
| AF241 | Australian Shepherd | E/E | Em/E<br>m | wt/wt | wt/wt | k*/k* | at/at | B/B | B/B | B/B | D/D | S/S | h/h |
| AF265 | Catahoula | E/E | Em/E<br>m | wt/wt | wt/wt | k*/k* | at/at | B/B | B/bs | B/B | D/D | S/S | h/h |
| AF267 | Miniature Australian Shepherd | E/E | wt/wt | wt/wt | wt/wt | k*/k* | at/at | B/bc | B/B | B/bd | D/D | S/S | h/h |
| AF270 | Labradoodle | E/e | Em/w<br>t | wt/wt | wt/wt | K/k* | Ay/at | B/bc | B/B | B/bd | D/D | S/S | h/h |
| AF273 | Catahoula | E/E | Em/w<br>t | wt/wt | wt/wt | K/k* | at/at | B/B | bs/bs | B/B | D/D | S/sp | h/h |
| AF274 | Catahoula | E/E | wt/wt | wt/wt | wt/wt | k*/k* | at/at | B/B | B/bs | B/B | D/D | S/S | h/h |
| AF327 | Australian Koolie | E/E | Em/w<br>t | wt/wt | wt/wt | k*/k* | at/at | B/B | B/bs | B/B | D/D | S/S | h/h |
| AF330 | Catahoula | E/E | Em/w<br>t | wt/wt | wt/wt | K/k* | at/at | B/B | B/bs | B/B | D/D | S/S | h/h |
| AF331 | Catahoula | E/E | Em/w<br>t | wt/wt | wt/wt | K/k* | at/at | B/B | bs/bs | B/B | D/D | S/S | h/h |
| AF332 | Catahoula | E/E | wt/wt | wt/wt | wt/wt | K/k* | at/at | B/B | bs/bs | B/B | D/D | S/S | h/h |
| AF333 | Catahoula | E/E | wt/wt | wt/wt | wt/wt | K/k* | at/at | B/B | B/bs | B/B | D/D | S/S | h/h |
| AF334 | Catahoula | E/E | wt/wt | wt/wt | wt/wt | k*/k* | at/at | B/B | bs/bs | B/B | D/D | S/S | h/h |
| AF335 | Catahoula | E/E | wt/wt | wt/wt | wt/wt | K/k* | at/at | B/B | bs/bs | B/B | D/D | S/S | h/h |
| AF341 | Catahoula | E/E | wt/wt | wt/wt | wt/wt | K/k* | at/at | B/B | B/bs | B/B | D/d | S/S | h/h |
| AF350 | Australian Koolie | E/E | wt/wt | wt/wt | wt/wt | k*/k* | aw/at | B/B | B/bs | B/B | D/d | S/sp | h/h |
| AF351 | Australian Koolie | E/E | Em/w<br>t | wt/wt | wt/wt | K/k* | at/at | B/B | bs/bs | B/B | D/d | S/sp | h/h |
| AF352 | Catahoula | E/E | wt/wt | wt/wt | wt/wt | K/k* | aw/at | B/B | B/B | B/B | D/D | S/S | h/h |
| AF378 | Catahoula | E/E | Em/w<br>t | wt/wt | wt/wt | k*/k* | at/at | B/B | B/bs | B/B | D/D | S/S | h/h |
| AF379 | Catahoula | E/E | Em/w<br>t | wt/wt | wt/wt | k*/k* | at/at | B/B | B/bs | B/B | D/D | S/S | h/h |
| AF380 | Catahoula | E/E | wt/wt | wt/wt | wt/wt | k*/k* | at/at | B/B | bs/bs | B/B | D/D | S/S | h/h |
| AF381 | Border Collie | E/E | Em/w<br>t | wt/wt | wt/wt | k*/k* | at/at | B/B | B/bs | B/B | D/D | S/S | h/h |
| AF382 | Border Collie | E/E | Em/E<br>m | wt/wt | wt/wt | K/k* | aw/at | B/B | B/B | B/B | D/d | S/S | h/h |
| AF398 | Australian Shepherd | E/E | Em/E<br>m | wt/wt | wt/wt | k*/k* | aw/at | B/bc | B/B | B/bd | D/D | S/S | h/h |
| AF399 | Catahoula | E/E | Em/w<br>t | wt/wt | wt/wt | K/k* | at/at | B/B | bs/bs | B/B | D/D | S/sp | h/h |
| AF400 | Australian Koolie | E/E | Em/w<br>t | wt/wt | wt/wt | K/k* | aw/at | B/B | bs/bs | B/B | D/d | S/S | h/h |
| AF401 | Rough Collie | E/E | wt/wt | wt/wt | wt/wt | k*/k* | at/at | B/B | B/B | B/B | D/D | S/sp | h/h |
| AF402 | Catahoula | E/e | wt/wt | wt/wt | wt/wt | k*/k* | at/a | B/B | B/bs | B/B | D/D | S/S | h/h |
| AF425 | Catahoula | E/E | Em/w<br>t | wt/wt | wt/wt | k*/k* | at/at | B/B | B/B | B/B | D/D | S/S | h/h |
| AF426 | Catahoula | E/E | Em/E<br>m | wt/wt | wt/wt | k*/k* | at/at | B/B | B/bs | B/B | D/d | S/S | h/h |
| AF431 | Border Collie | E/E | wt/wt | wt/wt | wt/wt | K/k* | at/at | B/B | bs/bs | B/B | D/D | S/S | h/h |
| AF432 | Australian Koolie | E/E | Em/w<br>t | wt/wt | wt/wt | K/k* | Ay/at | B/bc | B/bs | B/bd | D/D | S/S | h/h |
| AF457 | Australian Shepherd | E/E | Em/w<br>t | wt/wt | wt/wt | k*/k* | at/at | B/B | bs/bs | B/B | D/D | S/S | h/h |
| AF459 | Border Collie | E/E | wt/wt | wt/wt | wt/wt | K/K | at/at | B/B | bs/bs | B/B | D/d | S/S | h/h |
| AF460 | Australian Koolie | E/E | Em/w<br>t | wt/wt | wt/wt | K/K | at/at | B/B | B/bs | B/B | D/D | S/S | h/h |
| AF462 | Catahoula | E/E | Em/w<br>t | wt/wt | wt/wt | K/k* | at/at | B/B | B/bs | B/B | D/D | S/S | h/h |
| AF472 | Australian Shepherd | E/E | Em/w<br>t | wt/wt | wt/wt | k*/k* | at/at | B/bc | B/B | B/bd | D/D | S/sp | h/h |
| AF473 | Border Collie | E/E | Em/E<br>m | wt/wt | wt/wt | K/k* | aw/at | B/B | B/bs | B/B | D/D | S/S | h/h |
| AF474 | Australian Shepherd | E/E | wt/wt | wt/wt | wt/wt | k*/k* | at/a | B/bc | B/bs | B/bd | D/D | S/S | h/h |
| AF475 | Australian Shepherd | E/E | Em/w<br>t | wt/wt | wt/wt | K/k* | at/at | B/bc | B/B | B/bd | D/D | S/S | h/h |
| AF480 | Australian Koolie | E/E | wt/wt | wt/wt | wt/wt | K/k* | at/at | B/bc | B/bs | B/bd | d/d | S/S | h/h |
| AF481 | Australian Koolie | E/E | wt/wt | wt/wt | wt/wt | K/k* | at/at | B/B | bs/bs | B/B | D/d | S/S | h/h |
| AF482 | Australian Koolie | E/E | Em/w<br>t | wt/wt | wt/wt | K/k* | aw/at | B/B | B/bs | B/B | d/d | S/S | h/h |
| AF483 | Australian Koolie | E/E | Em/w<br>t | wt/wt | wt/wt | k*/k* | at/at | B/B | B/bs | B/B | D/d | S/S | h/h |
| AF484 | Australian Koolie | E/E | Em/w<br>t | wt/wt | wt/wt | k*/k* | at/at | B/B | bs/bs | B/B | D/d | S/S | h/h |
| AF485 | Australian Koolie | E/E | Em/w<br>t | wt/wt | wt/wt | k*/k* | at/at | B/B | B/bs | B/B | D/D | S/S | h/h |
| AF509 | Australian Shepherd | E/E | Em/w<br>t | wt/wt | wt/wt | k*/k* | at/at | B/B | B/bs | B/B | D/D | S/sp | h/h |
| AF510 | Catahoula | E/E | Em/w<br>t | wt/wt | wt/wt | k*/k* | at/a | B/B | bs/bs | B/B | D/D | S/S | h/h |
| AF511 | Catahoula | E/E | Em/w<br>t | wt/wt | wt/wt | k*/k* | at/a | B/B | B/bs | B/B | D/D | S/S | h/h |
| AF512 | Catahoula | E/E | Em/E<br>m | wt/wt | wt/wt | k*/k* | at/a | B/B | B/B | B/B | D/D | S/S | h/h |
| AF513 | Australian Koolie | E/E | Em/E<br>m | wt/wt | wt/wt | K/k* | at/at | B/B | B/bs | B/B | D/d | S/S | h/h |
| AF514 | Border Collie | E/E | Em/w<br>t | wt/wt | wt/wt | K/k* | at/at | B/B | B/B | B/B | D/D | S/S | h/h |
| AF515 | Catahoula | E/E | wt/wt | wt/wt | wt/wt | K/k* | at/at | B/B | B/bs | B/B | D/D | S/S | h/h |
| AF521 | Australian Koolie | E/E | Em/E<br>m | wt/wt | wt/wt | K/k* | at/at | B/B | B/bs | B/B | D/d | S/S | h/h |
| AF522 | Border Collie | E/E | wt/wt | wt/wt | wt/wt | K/K | aw/at | B/bc | B/bs | B/bd | D/[d]/<br>d | S/sp | h/h |
| AF523 | Pyrenean Shepherd | E/E | Em/E<br>m | wt/wt | wt/wt | K/k* | aw/aw | B/B | B/B | B/B | D/D | S/S | h/h |
| AF524 | Australian Shepherd | E/e | wt/wt | wt/wt | wt/wt | k*/k* | at/at | B/B | B/B | B/B | D/D | S/S | h/h |
| AF525 | Catahoula | E/E | Em/w<br>t | wt/wt | wt/wt | k*/k* | at/at | B/B | B/bs | B/B | D/d | S/S | h/h |
| AF578 | Dachshund | E/E | wt/wt | wt/wt | wt/wt | k*/k* | at/at | B/B | bs/bs | B/B | D/D | S/S | h/h |
| AF579 | Border Collie | E/E | wt/wt | wt/wt | wt/wt | K/k* | at/at | B/B | B/bs | B/B | D/D | S/S | h/h |
| AF580 | Australian Koolie | E/E | Em/w<br>t | wt/wt | wt/wt | K/k* | at/at | B/B | B/bs | B/B | D/D | S/sp | h/h |
| AF612 | Catahoula | E/E | wt/wt | wt/wt | wt/wt | K/k* | at/at | B/B | bs/bs | B/B | D/D | S/sp | h/h |
| AF613 | Welsh Sheepdog | E/E | Em/w<br>t | wt/wt | wt/wt | k*/k* | Ay/at | B/B | B/B | B/B | D/D | S/sp | h/h |
| AF614 | Welsh Sheepdog | E/E | Em/w<br>t | wt/wt | wt/wt | k*/k* | Ay/at | B/B | B/B | B/B | D/D | S/S | h/h |
| AF615 | Welsh Sheepdog | E/E | Em/w<br>t | wt/wt | wt/wt | K/k* | Ay/Ay | B/B | B/B | B/B | D/D | S/sp | h/h |
| AF649 | Mudi | E/e | wt/wt | wt/wt | wt/wt | K/k* | a/a | B/B | B/B | B/B | D/D | S/S | h/h |
| AF650 | Mudi | e/e | wt/wt | wt/wt | wt/wt | K/k* | a/a | B/B | B/B | B/B | D/D | S/S | h/h |
| AF652 | Australian Shepherd | E/E | wt/wt | wt/wt | wt/wt | k*/k* | aw/at | B/bc | B/B | B/bd | D/D | S/S | h/h |
| AF658 | Australian Shepherd | E/E | wt/wt | wt/wt | wt/wt | k*/k* | at/at | B/bc | B/B | B/bd | D/D | S/S | h/h |
| AF668 | Australian Shepherd | E/E | wt/wt | wt/wt | wt/wt | k*/k* | at/at | B/bc | B/B | B/bd | D/D | S/S | h/h |
| AF669 | Catahoula | E/E | Em/E<br>m | wt/wt | wt/wt | K/k* | at/a | B/B | B/B | B/B | D/D | S/S | h/h |
| AF694 | Catahoula | E/E | Em/w<br>t | wt/wt | wt/wt | K/k* | at/at | B/B | B/B | B/B | D/D | S/sp | h/h |
| AF696 | Australian Koolie | E/E | Em/E<br>m | wt/wt | wt/wt | K/k* | at/at | B/B | B/bs | B/B | D/D | S/S | h/h |
| AF697 | Australian Koolie | E/E | Em/w<br>t | wt/wt | wt/wt | K/K | at/at | B/B | B/bs | B/B | D/D | S/S | h/h |
| AF699 | Catahoula | E/E | wt/wt | wt/wt | wt/wt | K/k* | at/at | B/B | B/B | B/B | D/D | S/S | h/h |
| AF758 | Australina Shepherd | E/E | wt/wt | wt/wt | wt/wt | K/k* | at/at | B/bc | B/B | B/bd | D/d | S/S | h/h |
| AF759 | Australian Shepherd | E/E | wt/wt | wt/wt | wt/wt | K/k* | at/at | B/B | B/bs | B/B | D/d | S/sp | h/h |
| AF760 | Australian Shepherd | E/E | wt/wt | wt/wt | wt/wt | k*/k* | at/at | B/bc | B/B | B/bd | D/D | S/sp | h/h |
| AF761 | Australian Shepherd | E/E | wt/wt | wt/wt | wt/wt | K/k* | at/at | B/B | B/B | B/B | D/d | S/S | h/h |
| AF762 | Australian Shepherd | E/E | wt/wt | wt/wt | wt/wt | k*/k* | at/at | B/bc | B/bs | B/bd | D/D | S/sp | h/h |
| AF778 | Border Collie | E/E | Em/w<br>t | wt/wt | wt/wt | K/k* | at/a | B/B | B/bs | B/B | D/D | S/sp | h/h |
| AF781 | Border Collie | E/e | wt/wt | wt/wt | wt/wt | K/k* | at/at | B/B | B/bs | B/B | D/D | S/S | h/h |
| AF782 | Border Collie | E/E | wt/wt | wt/wt | wt/wt | K/K | at/at | B/B | B/bs | B/B | D/D | S/S | h/h |
| AF824 | Australian Koolie | E/E | wt/wt | wt/wt | wt/wt | K/k* | at/at | B/B | bs/bs | B/B | D/D | S/S | h/h |

Interestingly, during the genotyping process, we came across some irregularities, especially concerning the A locus in some Border Collies and Shetland Sheepdogs, where the SINE insert in the ASIP gene has been regularly missed. Similar phenomenon has been observed for B locus testing in French Bulldogs. To elucidate the genetic background of both recurrent genotyping errors, further research is warranted.

## DISCUSSION

The Merle coat pattern has long been a fascinating feature for many breeders. It has attracted attention not only because of its visual uniqueness, but also for its negative health consequences, which have been observed rather frequently – vision and hearing impairment of the affected animals. In 2006, Clark et al. [12] mapped the genomic abnormality to the *SILV* gene and identified a SINE element inserted at the vicinity of exon 10. SINEs (Short Interspersed Elements) are retroviral relics that parasitized the genome of vertebrates millions of years ago ([20], [21], [22]). SINEs are thought to be a great source of genome evolution, but have also been shown to be involved in many human genetic disorders and cancer, in animals also in peculiar phenotypic features, such as Locus A variants (alleles “a” and ,, “at”).

A typical SINE element contains a poly-A tail, which harbors a stretch of homogeneous A nucleotide repetitions. This genomic feature poses a rather challenging situation for the cellular replication machinery and leads to a replication slippage and uneven replication products. This eventually leads to the senescence of the parasitic retroelement – shortening of the poly tail under a critical level buries the element at its original site with no transposition activity left. As shown for *SILV* SINE, shortening of the primary poly-A tail leads also to the atenuation of the local effect of the SINE insert on the surrounding genomic environment, with visible phenotypic consequences.

It was originally assumed that the poly-A tail of SINE inserted in the *SILV* gene might not only shorten in time, but also extend its length. This information has previously been theorized based only on phenotype and not genotype. We have studied the pedigree of related dogs in our cohort as reported in the Supplementary data (Table S1) and measured exactly the length of the obligatory alleles (within 1-2 bp size difference) and their passage from parent to the offspring, and siblings and cannot confirm this theory. We suggest, for the first time, that parental alleles are conserved in length between generations and it is highly probable that the alleged phenomenon of the poly-A extension does not exist. This finding is of special practical importance for breeders, as animals harboring critically shortened Merle alleles would not pose any danger in terms of health issues to their offspring and might safely be bred with full Merle individuals. Further research focused on precise monitoring of the length of Merle alleles is needed to clarify the poly-A dynamics of *SILV* SINE.

Another extremely interesting output of our study is the identification of a high degree of mosaicism in a high proportion of the animals tested. In 30 out of 181 dogs analyzed, more than one allelic type was identified. This finding was made possible by leveraging the high-resolution fragment analysis technology that allowed us to discriminate 7 allelic variants of the Merle gene, 4 of them (Mc+, Ma, Ma+ and Mh) previously unknown. It is highly probably that the existence of various allelic variants mirrors the complex genomic situation during the early embryonic development, when populations of cells with different SINE allelic versions may arise. Importantly, the Merle genotype pattern, as we have found, m may differ among the individual compartments tested. Sperm cells are the most heterogeneous tissue in terms of Merle allelic variability, with the possilibity of many minor Merle alleles idenfied, which can be passed to the next generation in a Mendelian fashion. Buccal swab, generally used for routine genotyping, may not mirror the germline status, be it a terminally differentiated ectodermal tissue. This phenomenon must be taken into consideration especially in those cases, when buccal swab-genotyped animals produce an phenotypically unexpected offspring. In those instances it is strongly recommended to test sperm cells of the sire for possible mosaicism that would explain the unexpected breeding results. For males often used for breeding, we strongly recommnend to test their sperm cells first, to reveal Merle genetic background in full context and thus, address the possibility of passing minor Merle alleles to offspring. With females, the situation is complicated by the impossibility to obtain germinal cells. Nevertheless, to reveal possible mosaicism, hair from Merle looking coat areas can be tested in parallel with buccal swab.

As already mentioned, the length of the SINE poly-A seems to be crucial as to the biological effects and phenotype correlations (pigment cells ontogenesis). Thus, critically shortened Merle alleles, exhibiting no visual pigment defects, such as Mc, Mc+ or Ma may also be expected to lose the biological effectsin terms of hearing or vision impairment development. Our preliminary data support this hypothesis, though more research is needed to elucidate this most important biological aspect of Merle biology. In line with it,(Moreover?) SINE inserts with a long poly-A tail, such as M or Mh may pose a greater risk of auditory or ophthalmic irregularities. It has been widely accepted that the most pronounced negative health consequences (both auditory and ophthalmic) are connected with the M/M genotype. Interestingly, our preliminary data show that even a heterozygous combination of m/Mh may exert the same effects – in our cohort of 181 dogs we identified 10 dogs with the following genotypes with associated hearing and/or vision impairments (Fig 15):

**Fig 15:**
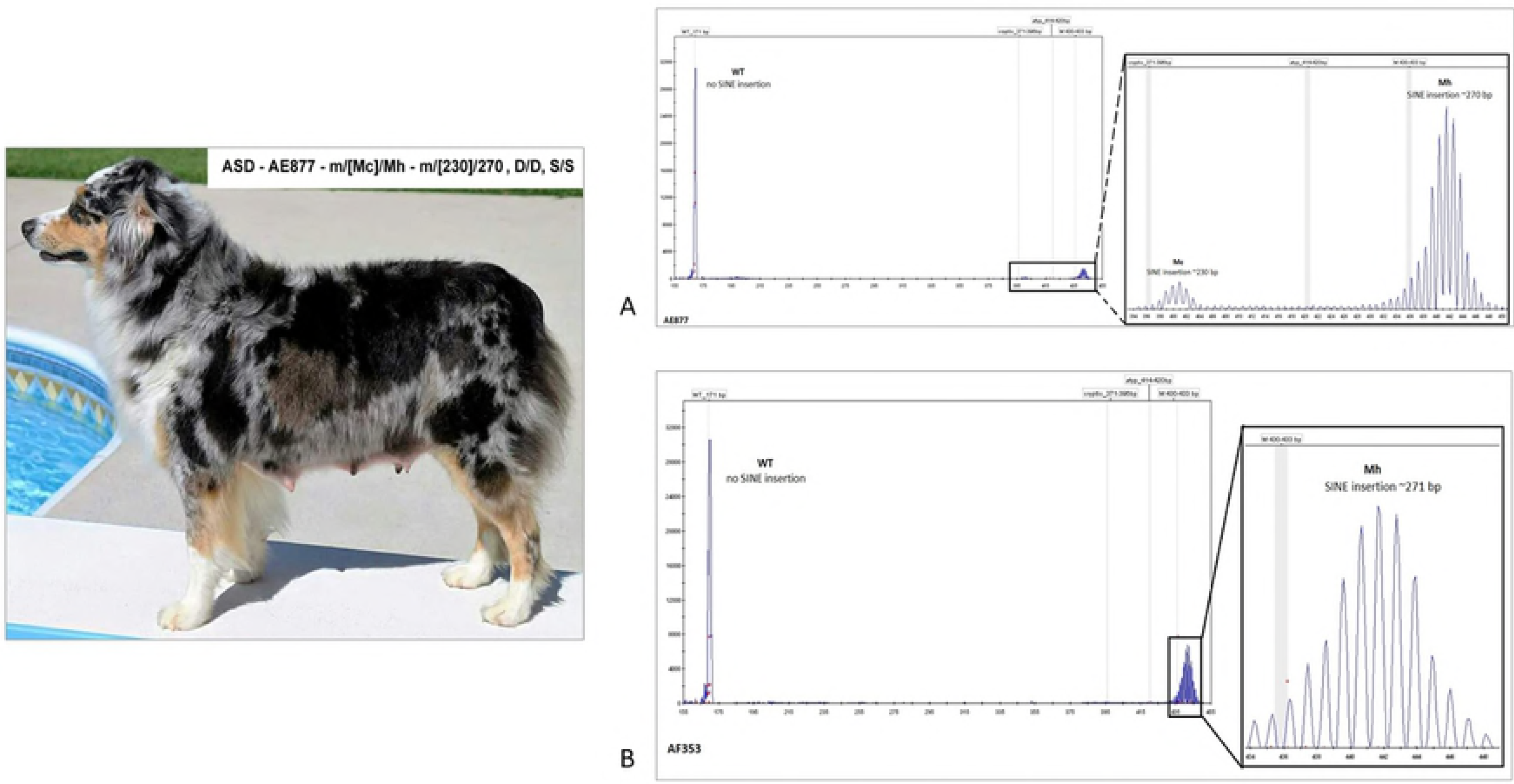
Merle allelic status and associated hearing and vision impairment

Harlequin Merle (Mh) also seems to explain more white in heterozygous Merle dogs.

An additional finding in our study considers the Tw – Tweed locus, that has been theorized to be a modifier of Merle. Named after a Merle pattern that expresses with random shaded-in areas, usually with two or three distinguishable shades. Based upon results from our study, we challenge the existence of the Tweed allele and rather ascribe the Tweed phenotype to different combinations of *SILV* SINE Merle alleles, see Figs 3A - 3X

In conclusion, our study has brought to light novel findings in the field of Merle research and challenged some theorized data, which might exert undesirable effects on breeding strategies in Merle dogs. The results from our study are hoped to be instrumental not only to researchers working in the field of coat color genetics in dogs, but especially for breeders in developing their breeding strategies in Merle breeds – to preserve the health of their animals to the fullest extent thenatural laws of population genetics allow.

## ACKNOWLEDGEMENTS

ML, HS, TJ and SP conceived the work, analyzed the data and prepared the manuscript. SP and TJ collected the laboratory data. ML and HS were responsible for contacting breeders and obtaining photographs of animals under study.

We would like to thank all the dog owners who enabled this research by contributing samples and pictures of their dogs. Our special thanks go to the following enthusiastic breeders and friends:

## Karen Smiley Combs - KC Aussies, Georgia, USA

For tirelessly educating many in the Australian Shepherd community. For guiding owners through the process of testing, for explaining results and helping breeders to understand the oddities they are seeing in whelping boxes.

## Tanita Bruce - President of the Australian Koolie Association, Australia

For coordinating the testing of all Australian Koolies represented and funding the majority of those 32 tests. For her dedication to educating those in all breeds.

**Dr. Marcela Françoso - Veterinarian and Border Collie enthusiast, Brazil.** As a vet, Marcela gives seminars on color, health issues and breeding strategies. Recognizing the different types of phenotypes in Merle Border Collies she arranged for testing on a large group of dogs from Brazil.

## Candice Jankuta – Cardi Catahoulas, Edmonton, Alberta

Candice was instrumental in helping Mary Langevin to identify the phenotype of Ma+, having wondered for many years why so many of her dogs had such an unusually subtle Merle pattern. They would spend countless hours back and forth over pedigrees and litter results, identifying Ma+.

**Alena Líšková** – being a hobby breeder and enthusiast, for her invaluable assistance with educating Merle dog breeders in Slovakia.

**Sally Young –** for her assistance with proof-reading

---

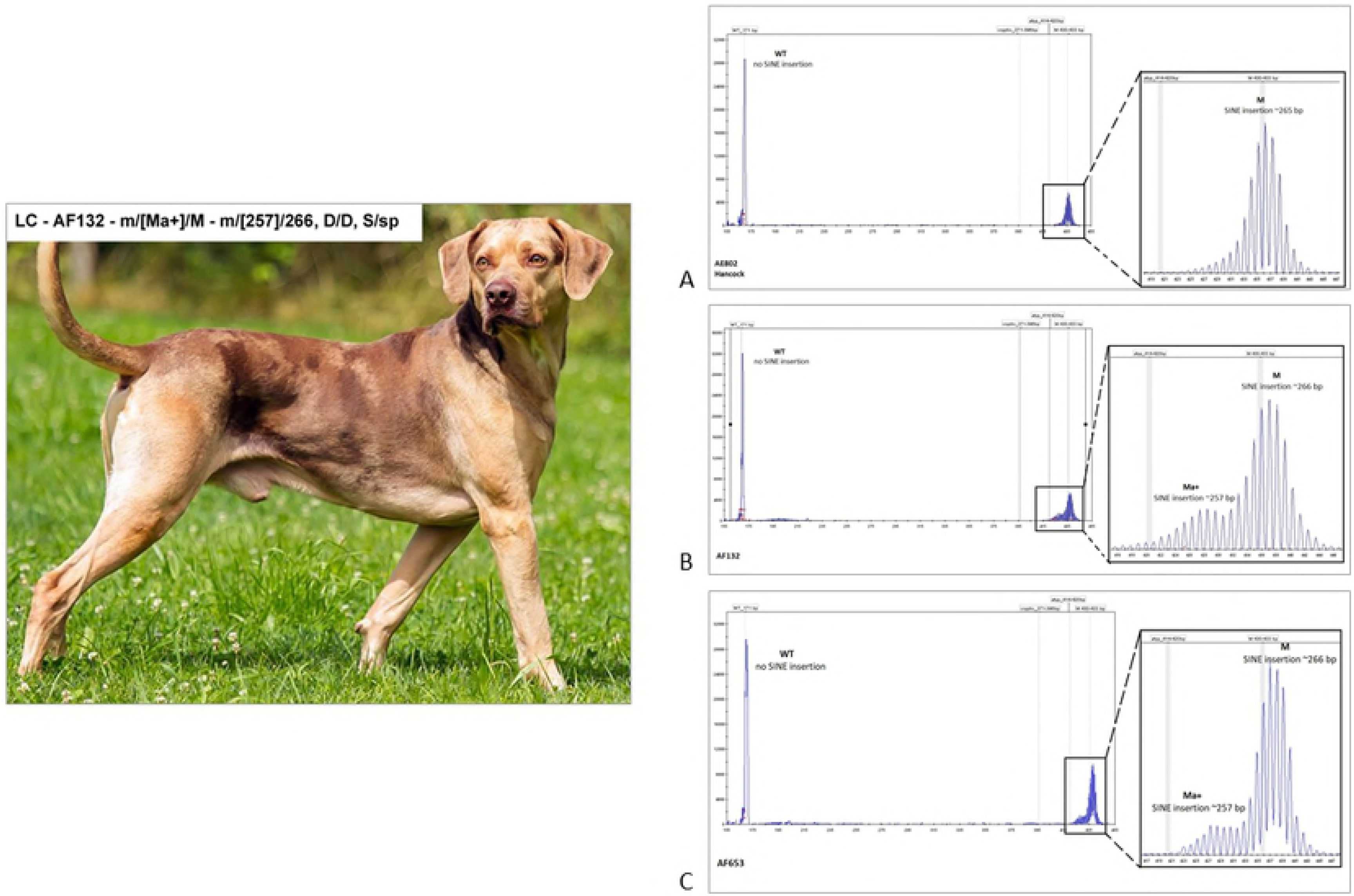

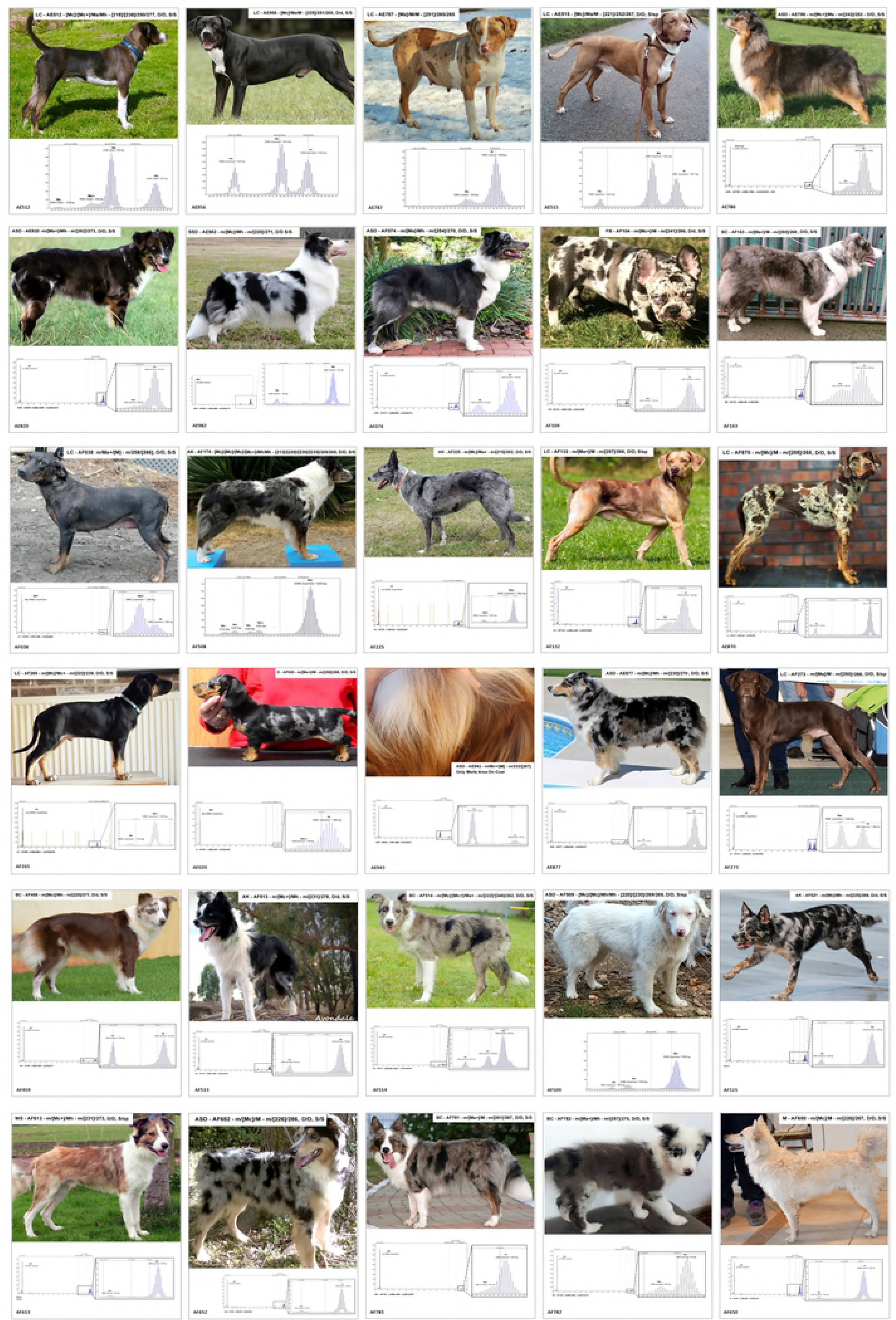

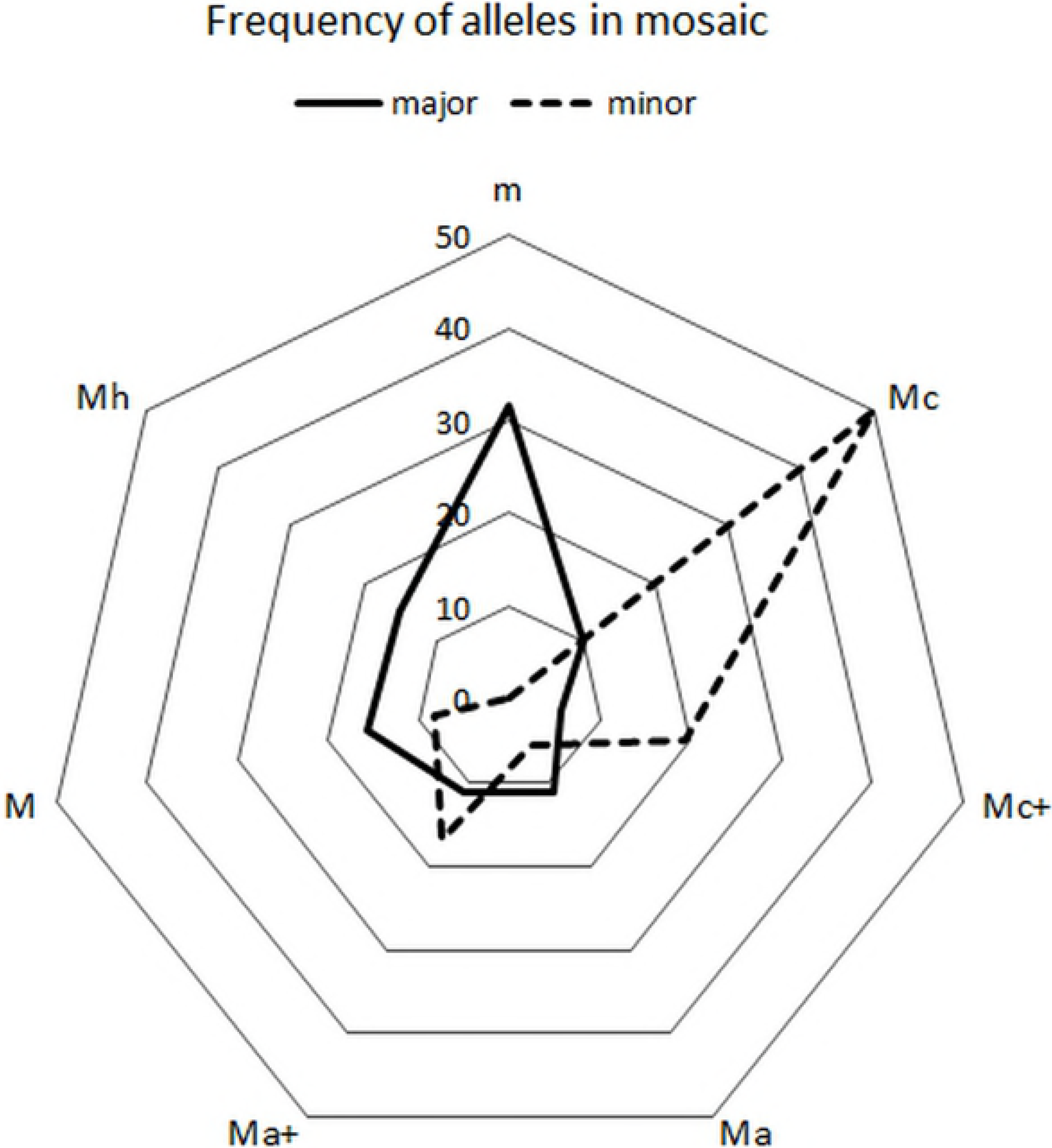

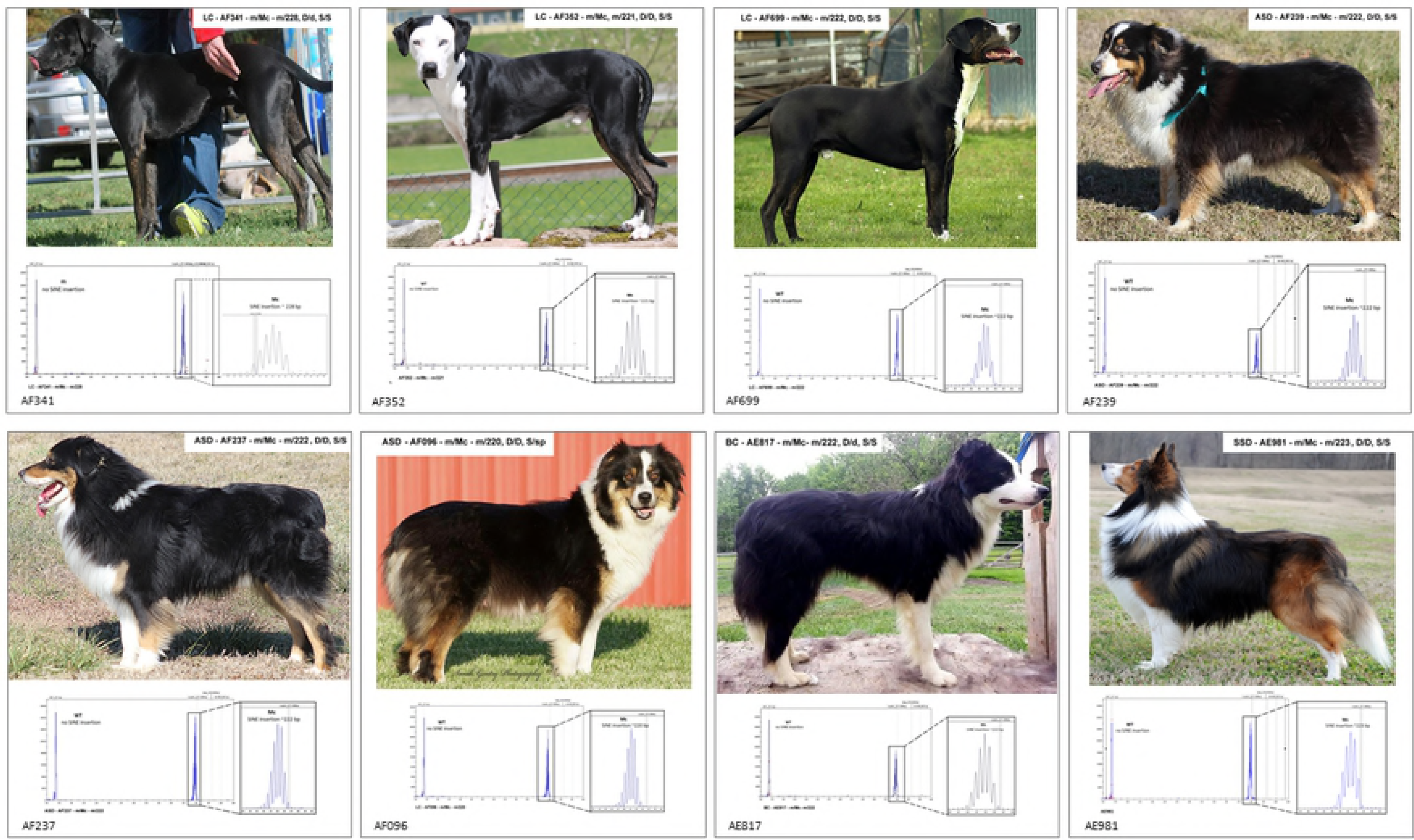

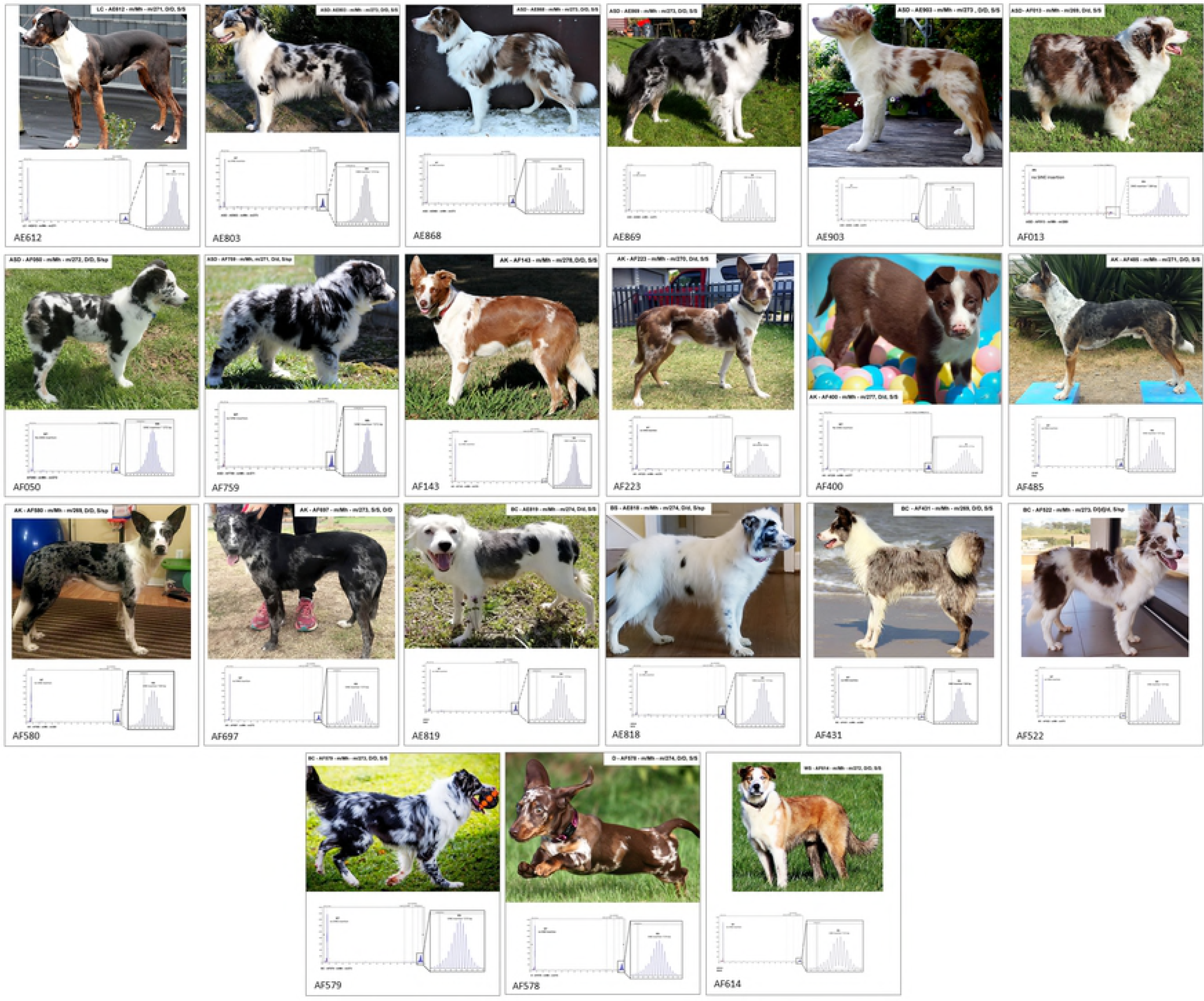

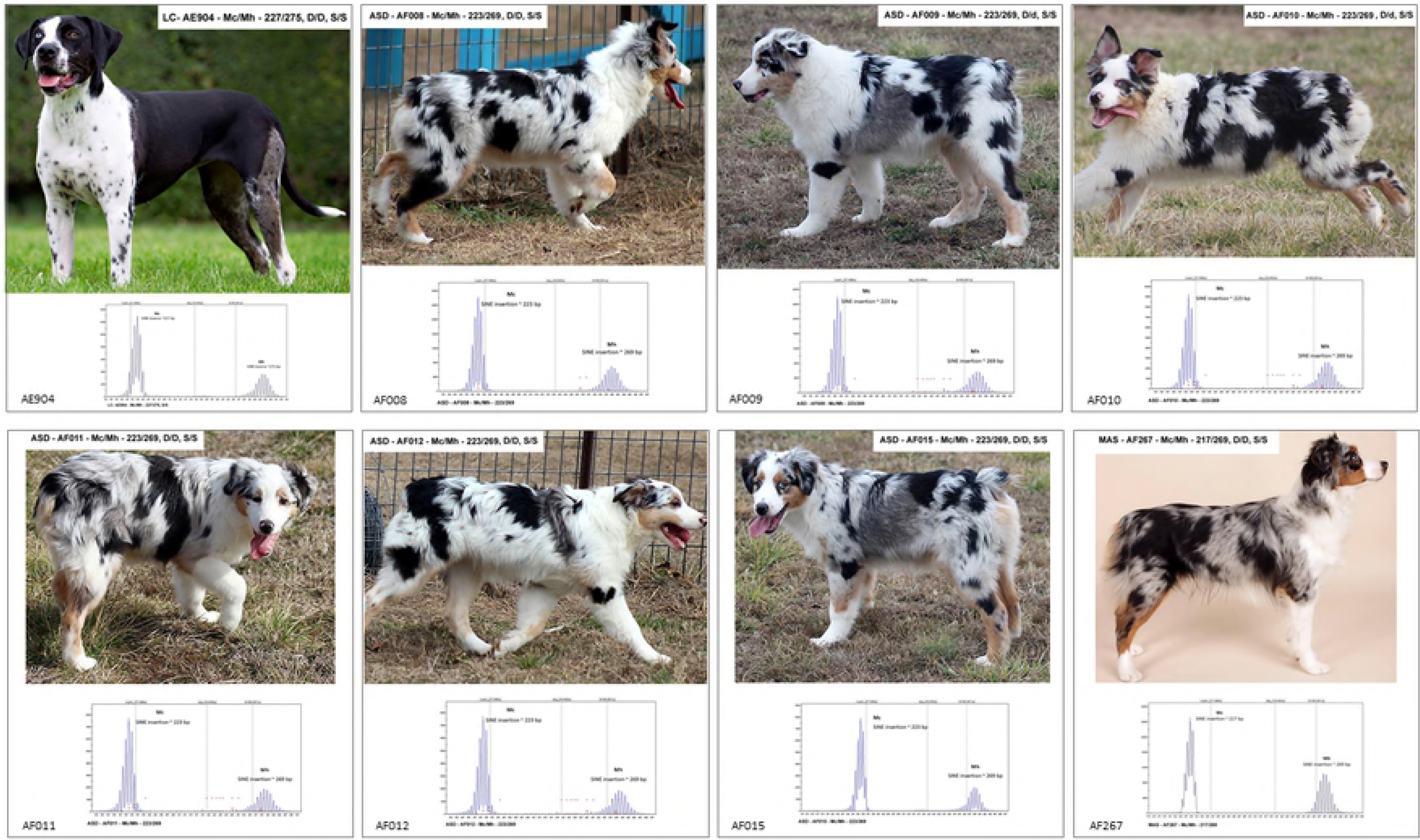

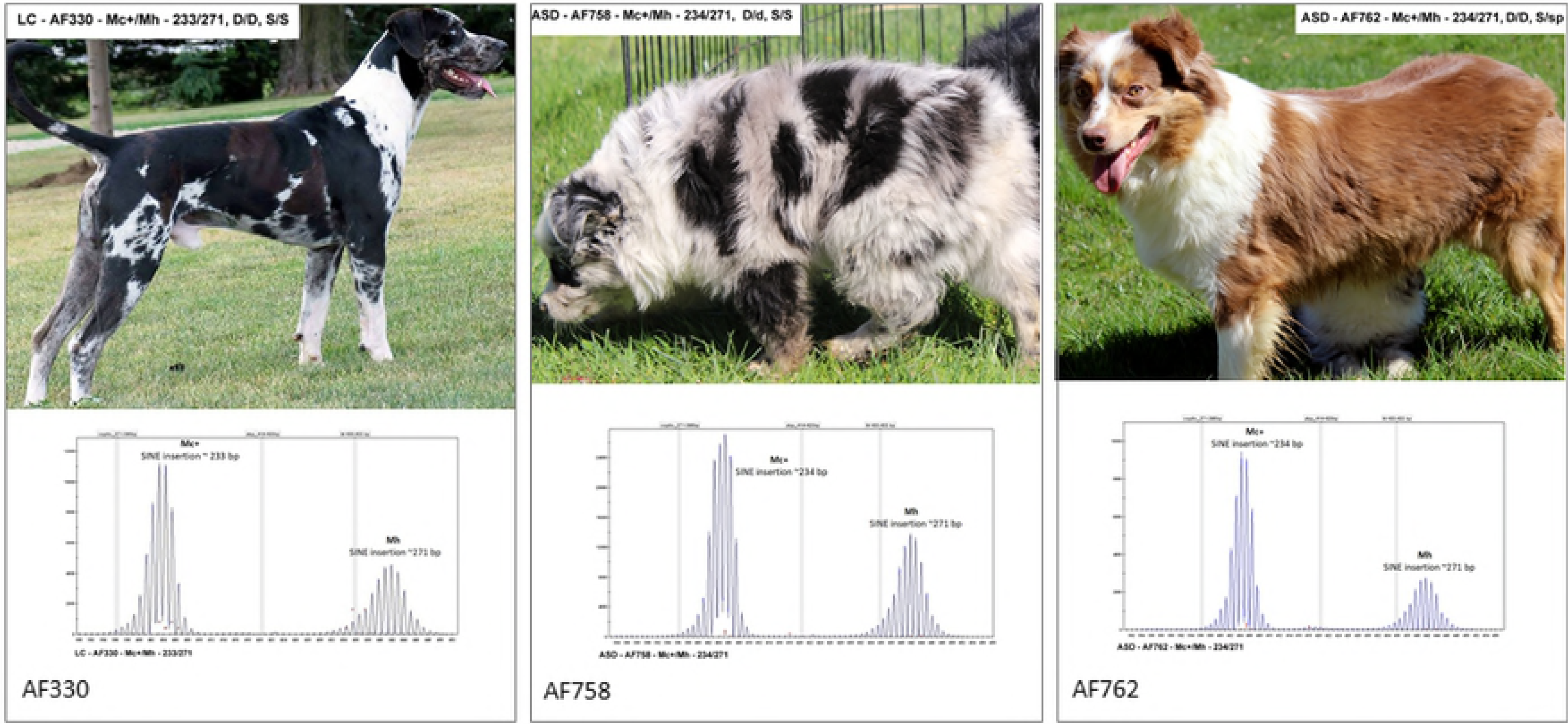

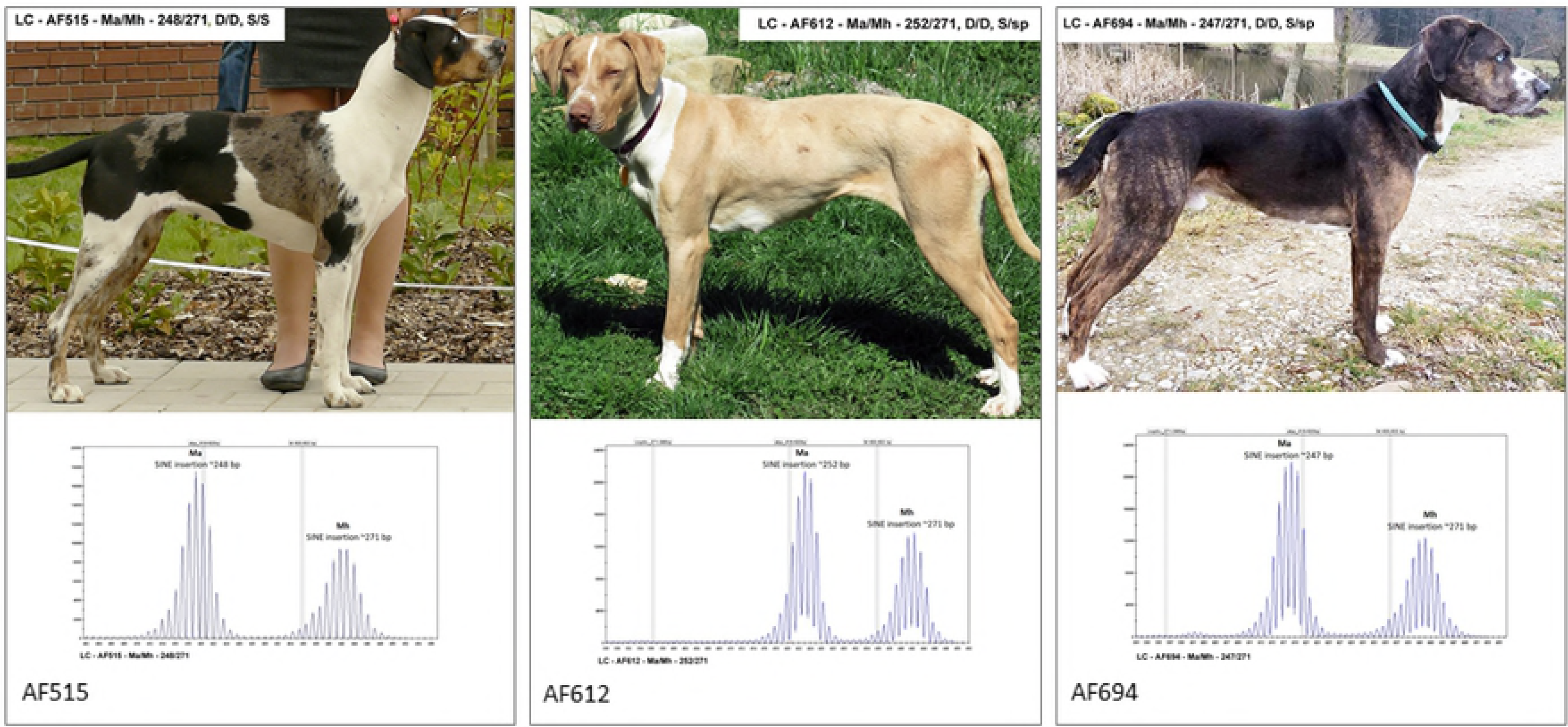

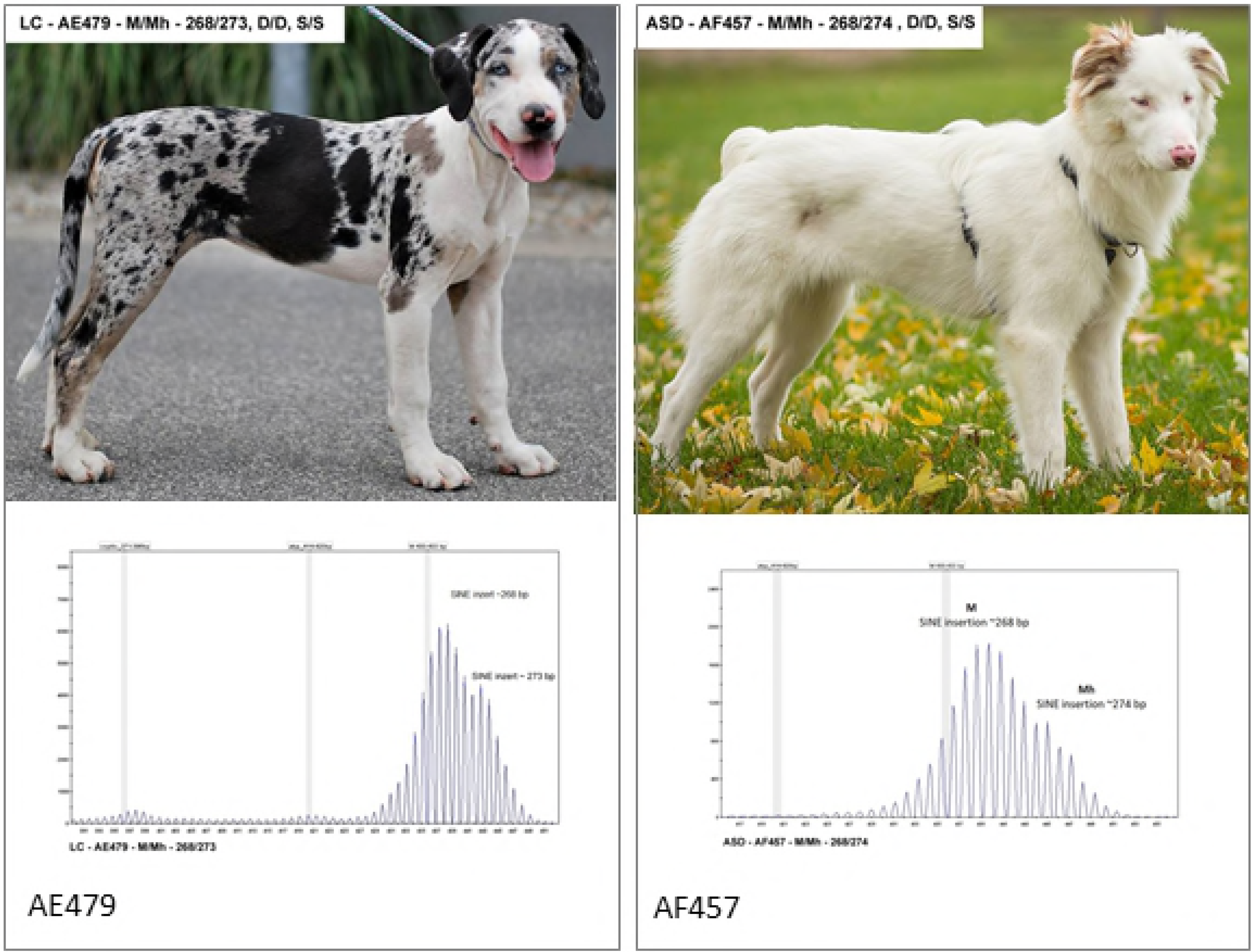

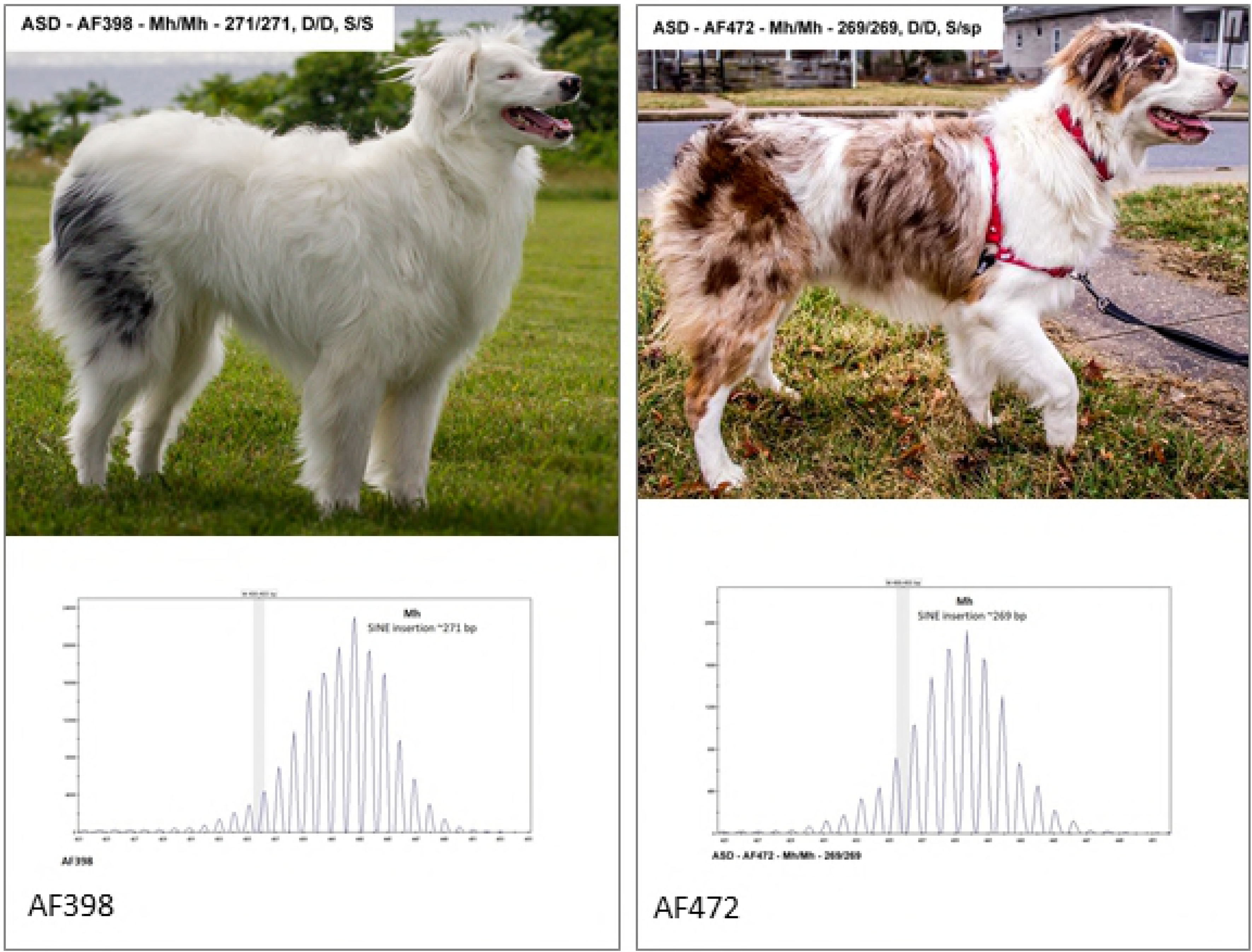

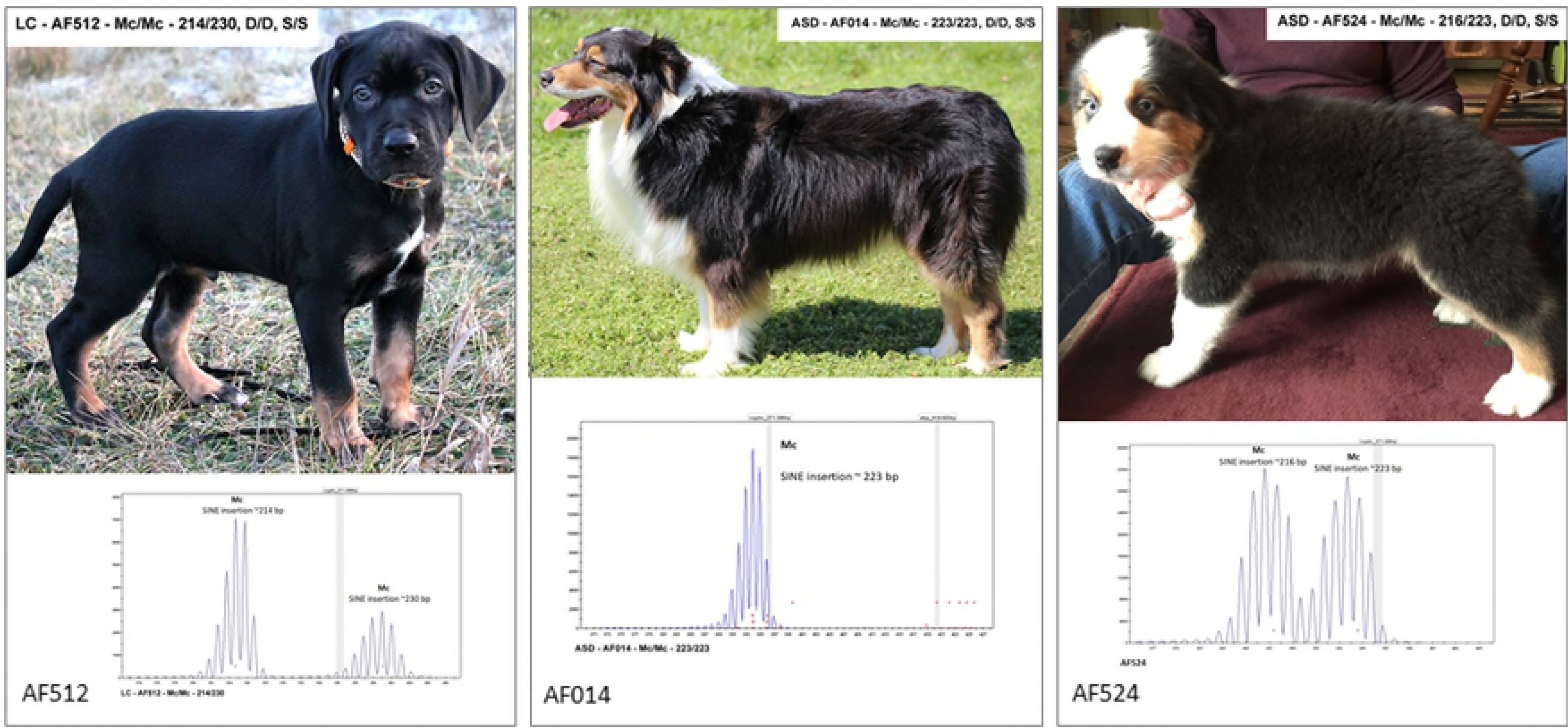

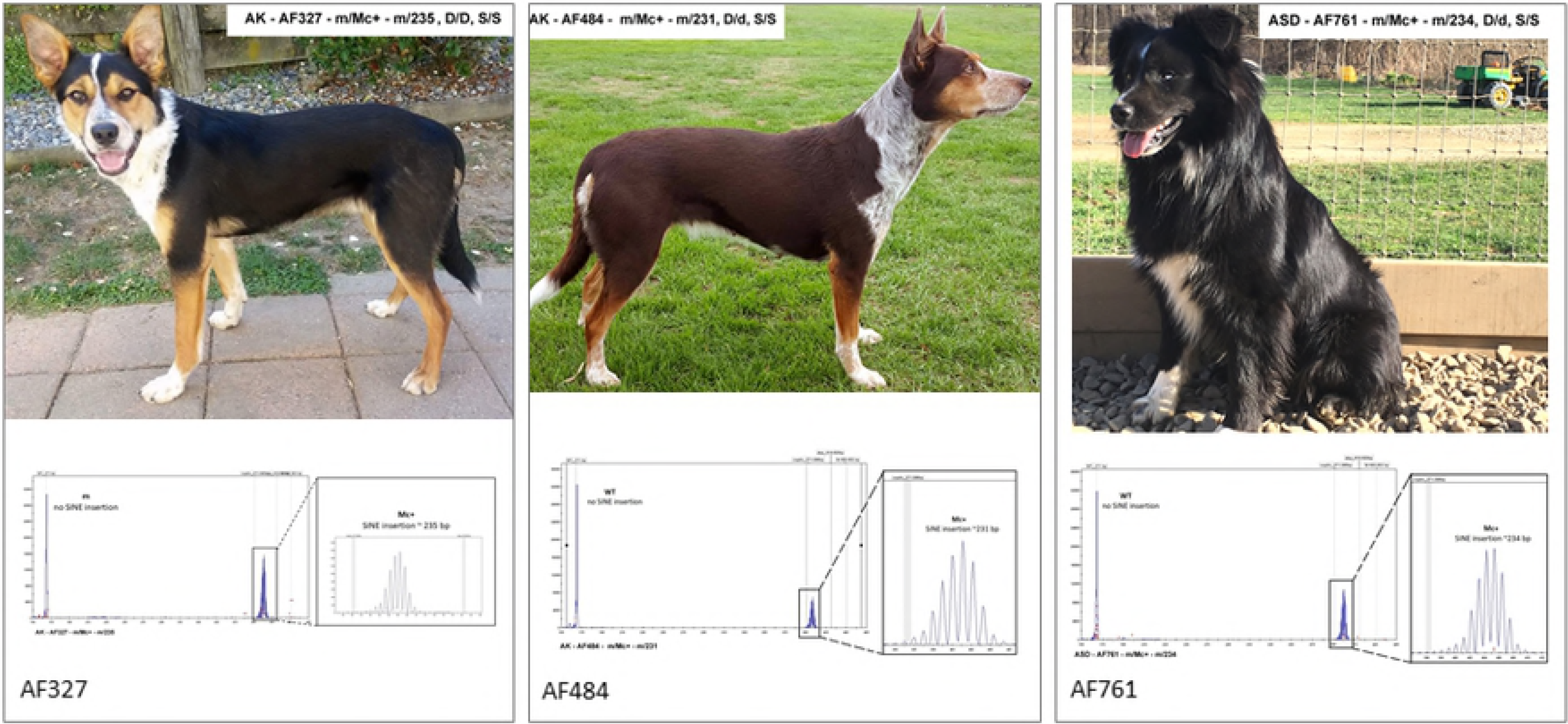

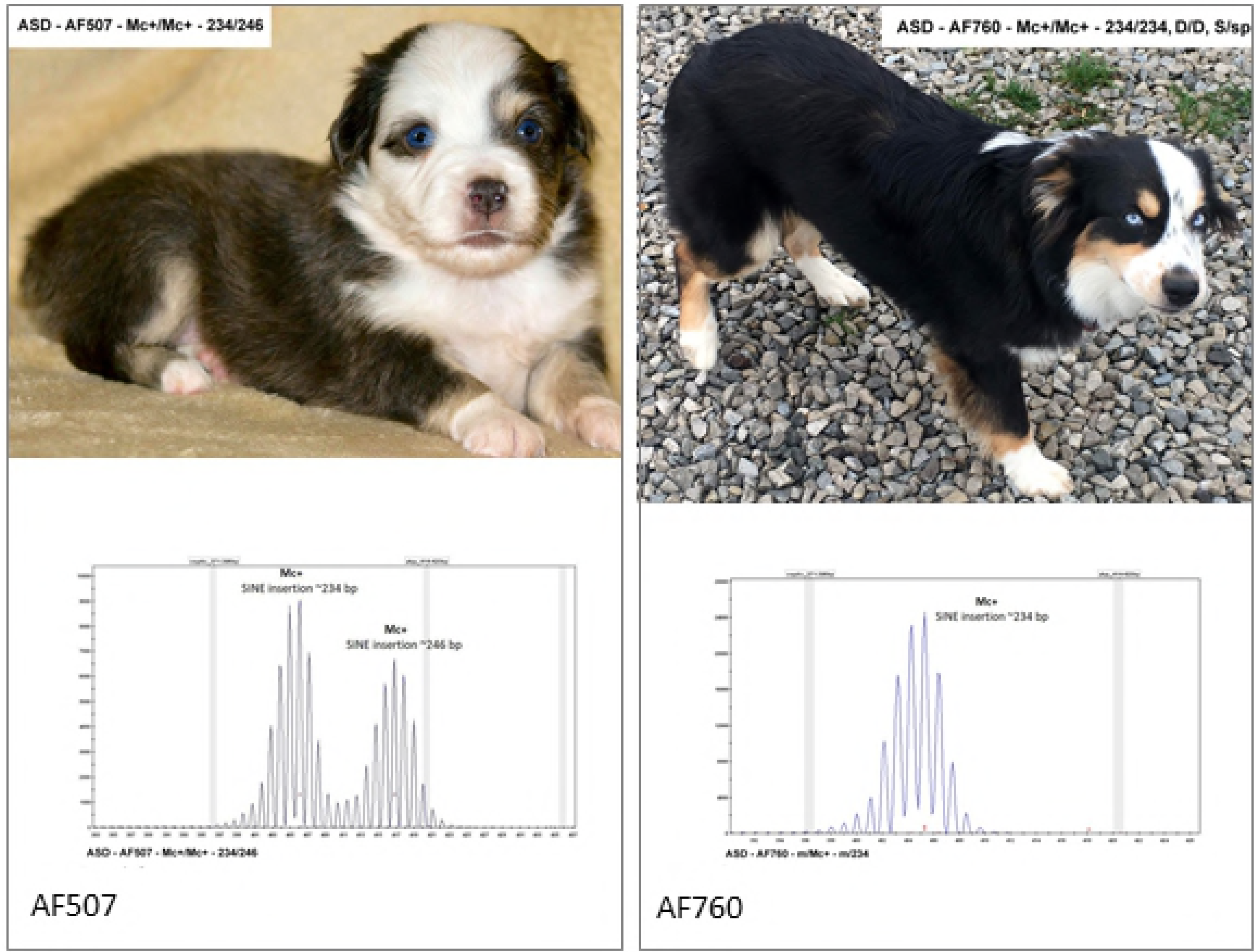

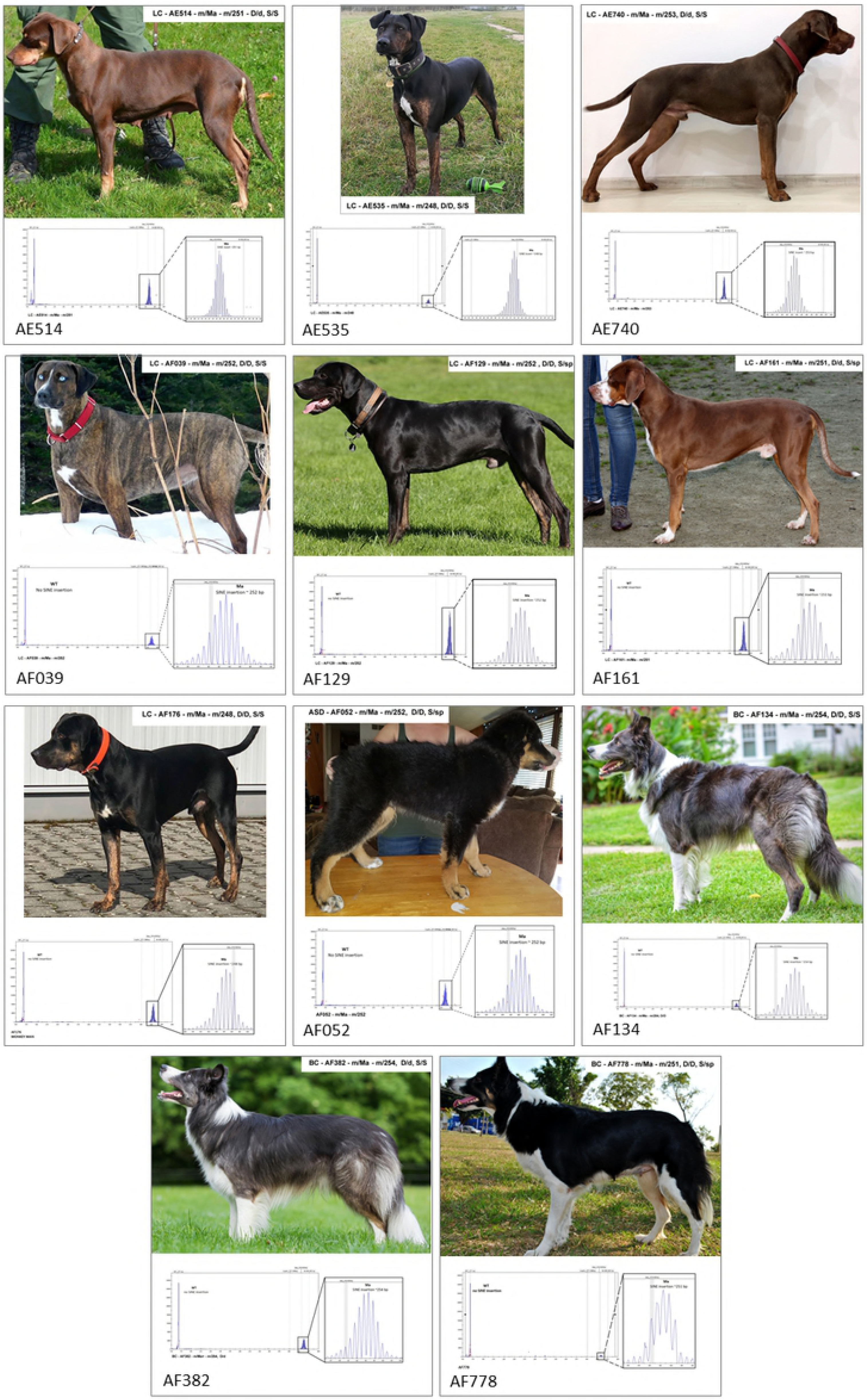

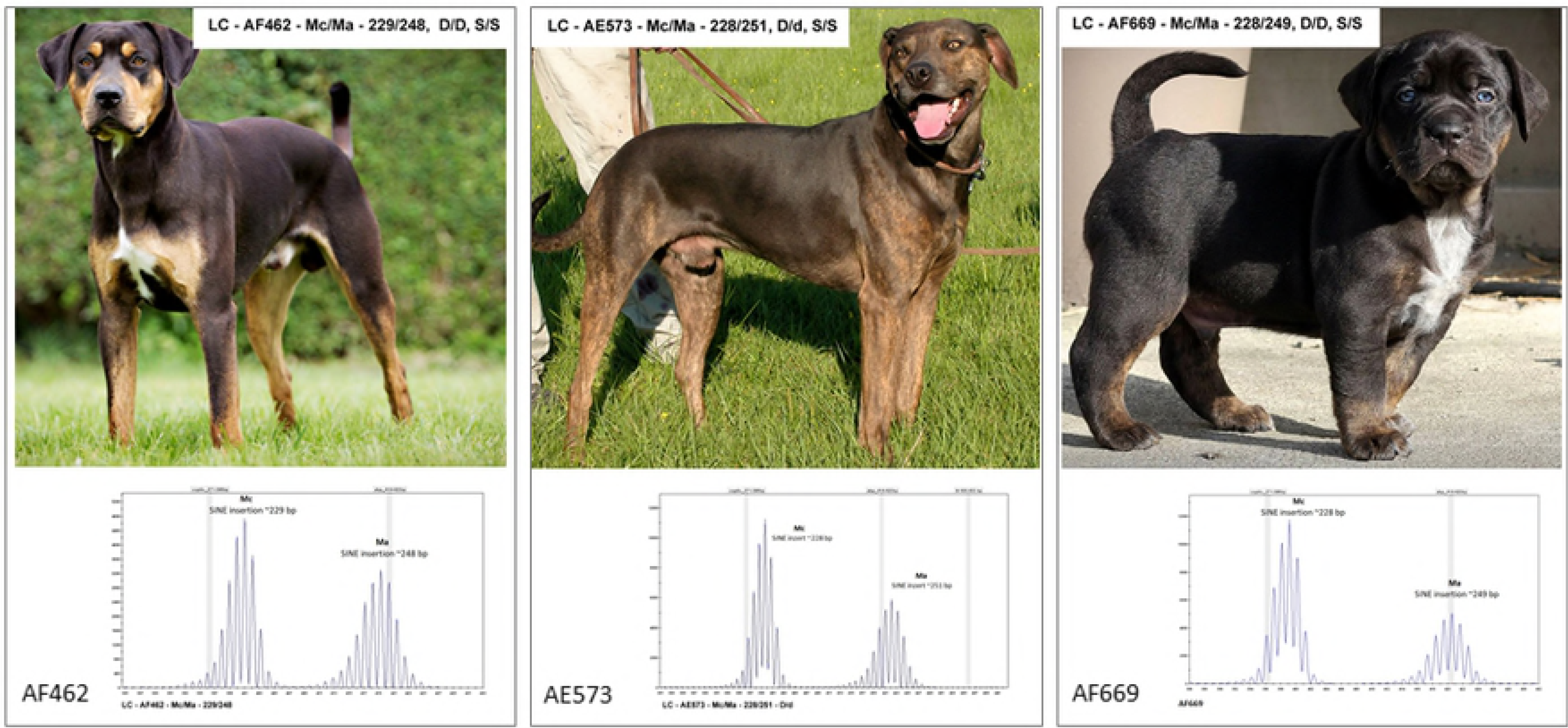

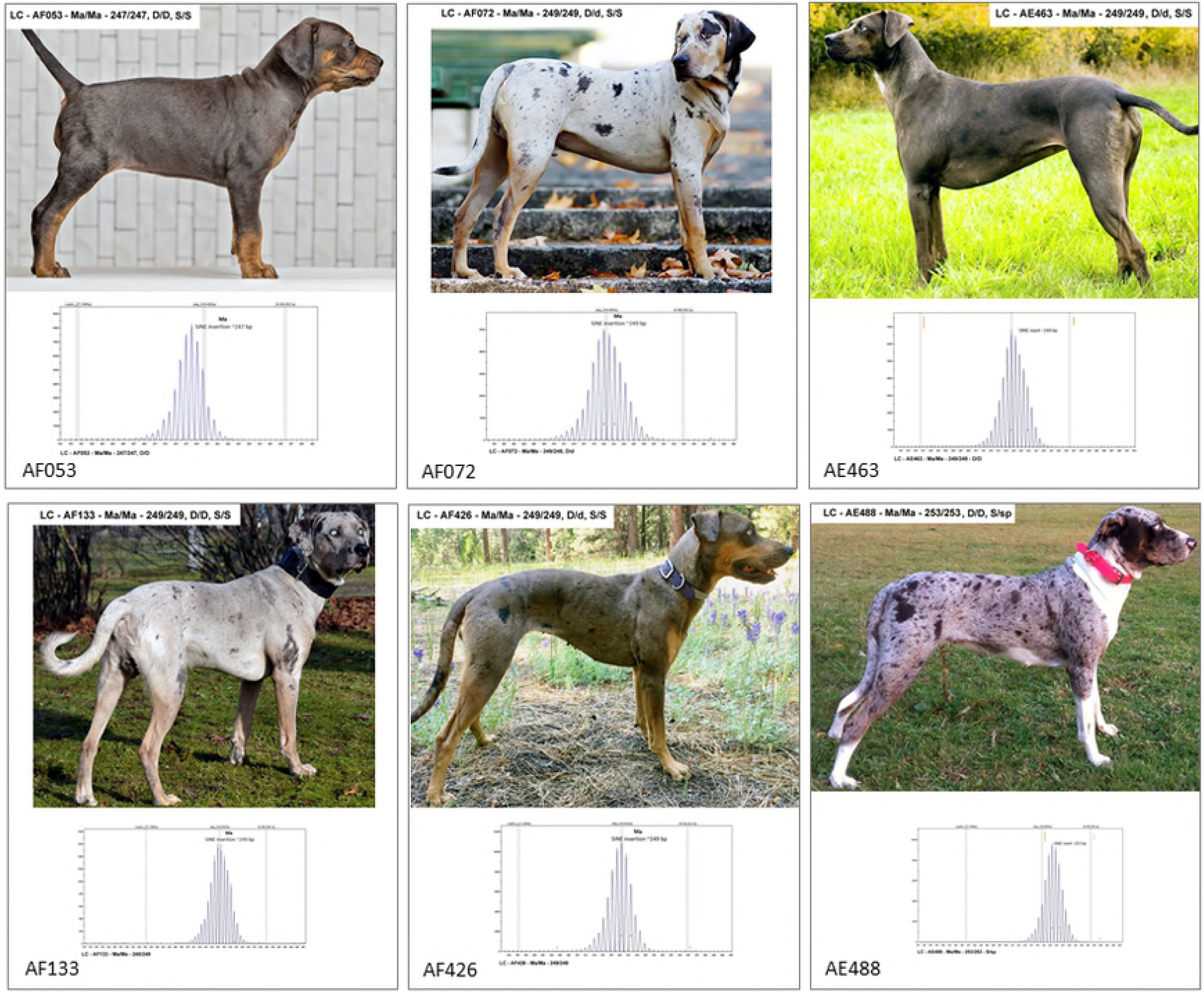

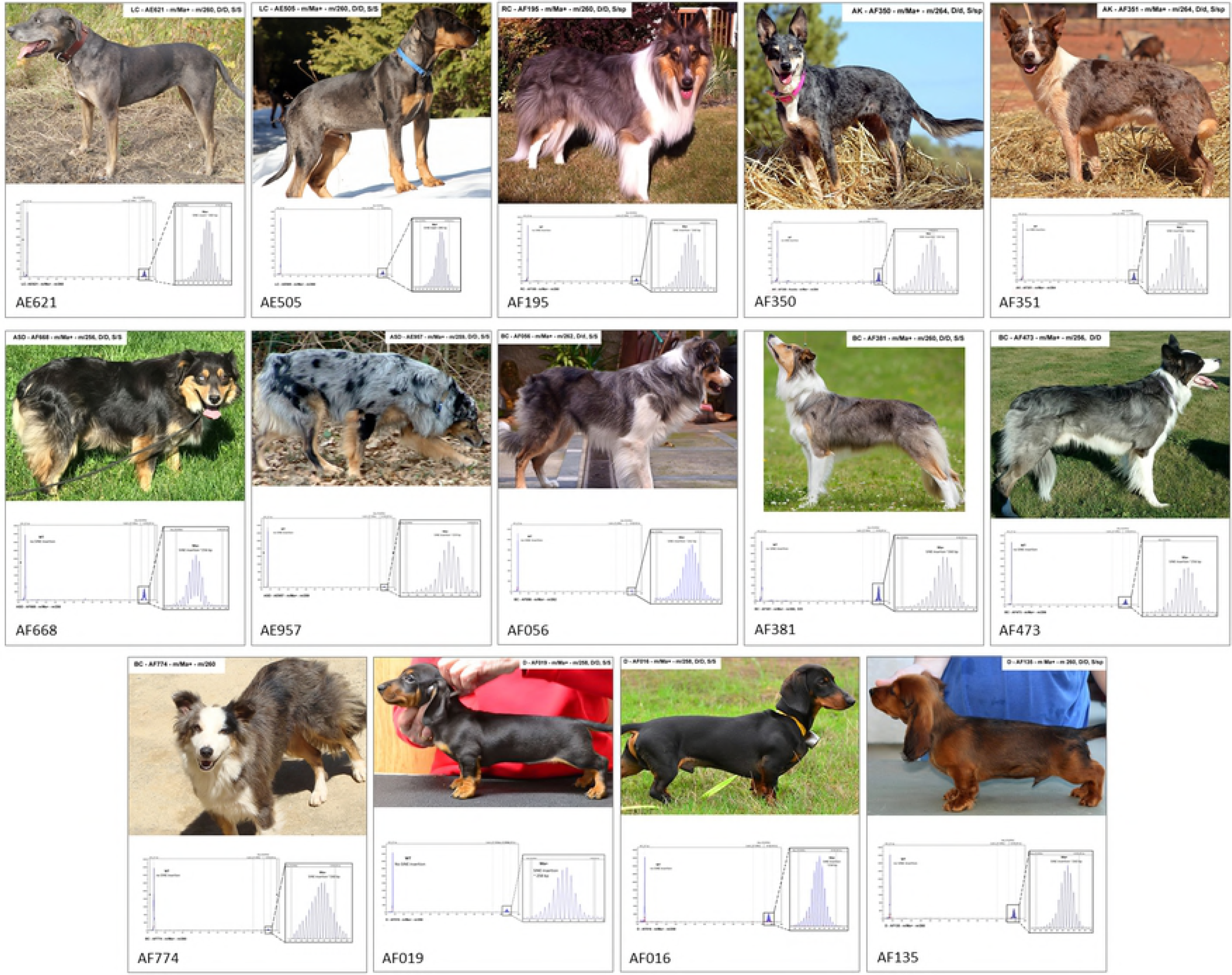

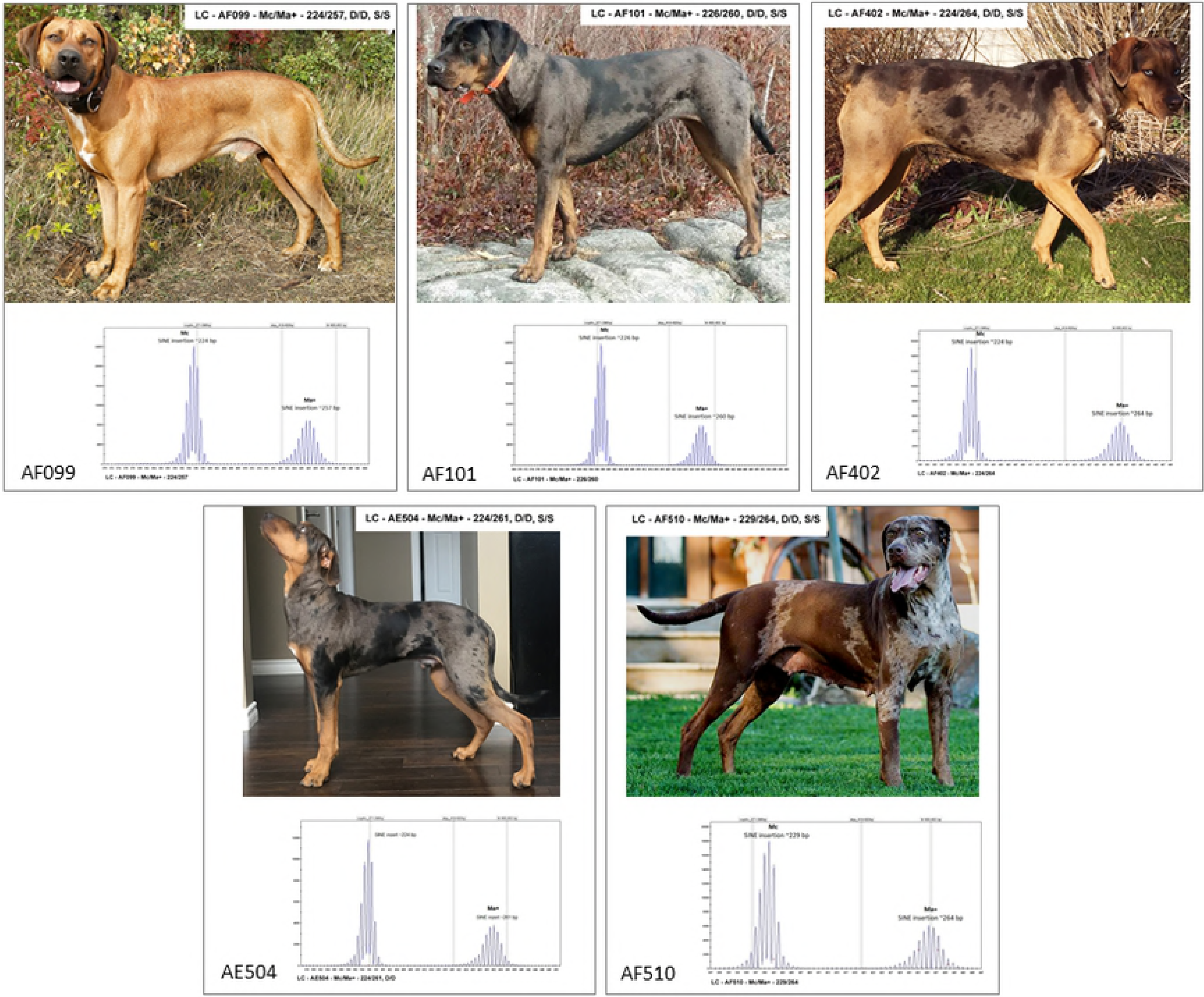

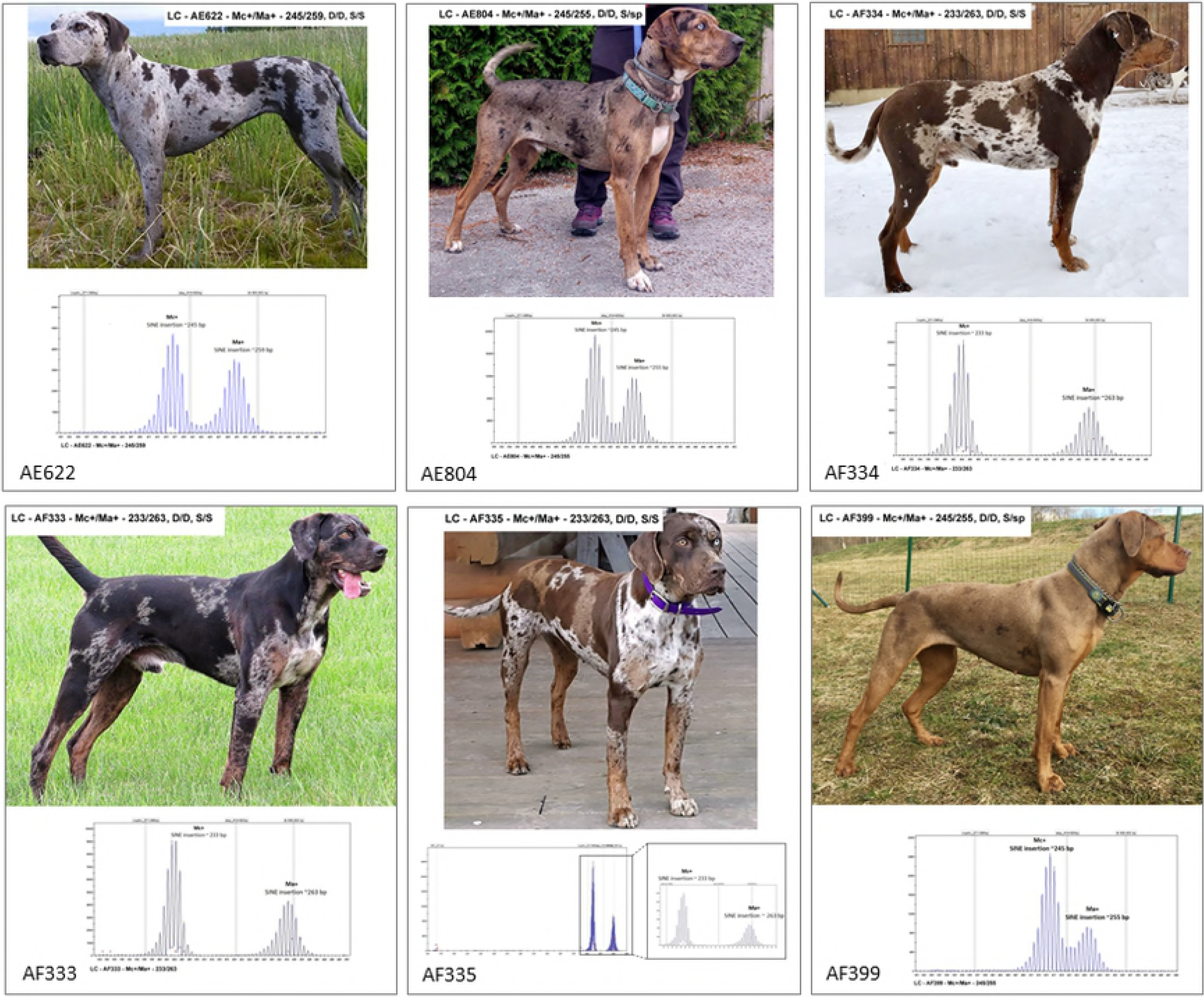

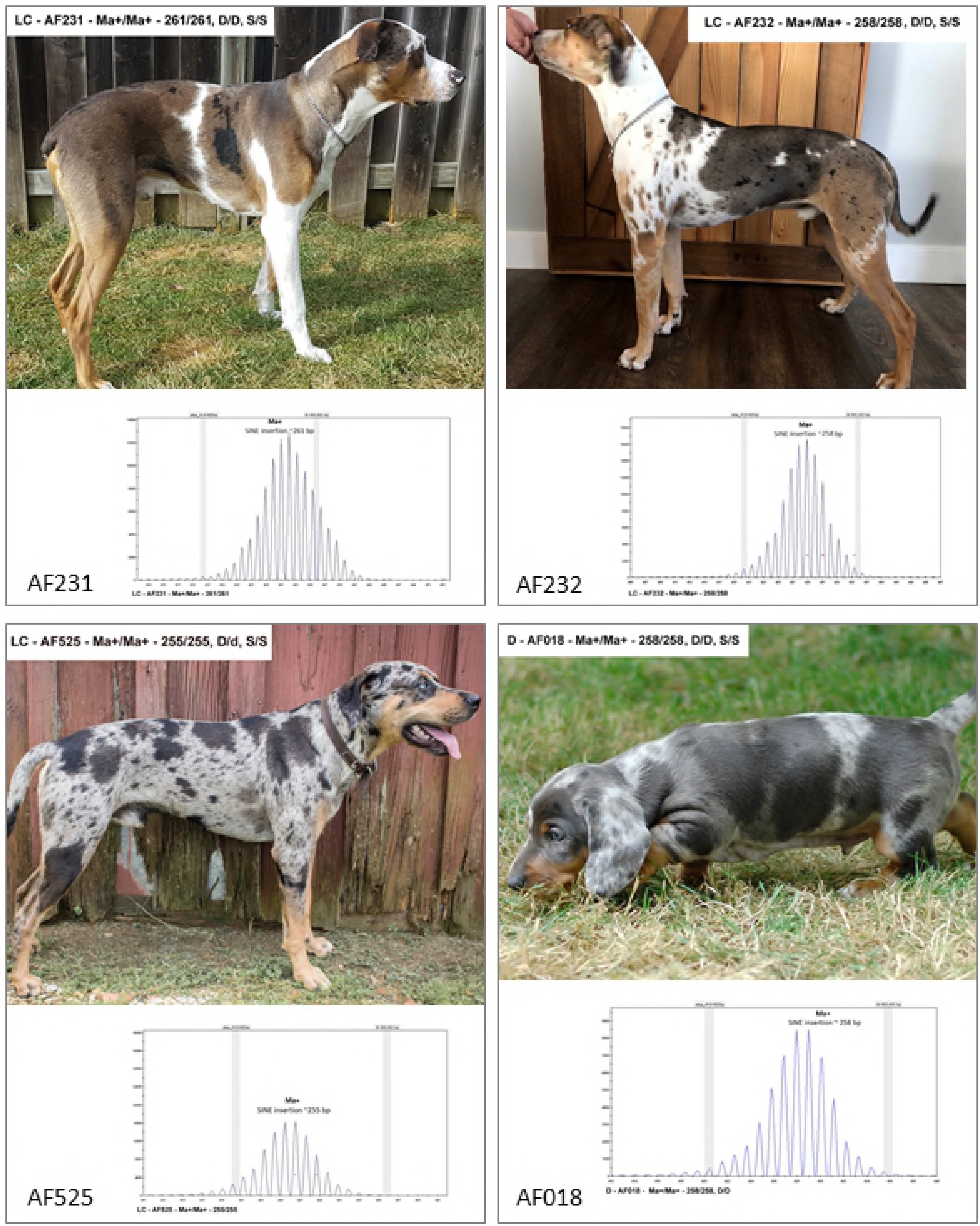

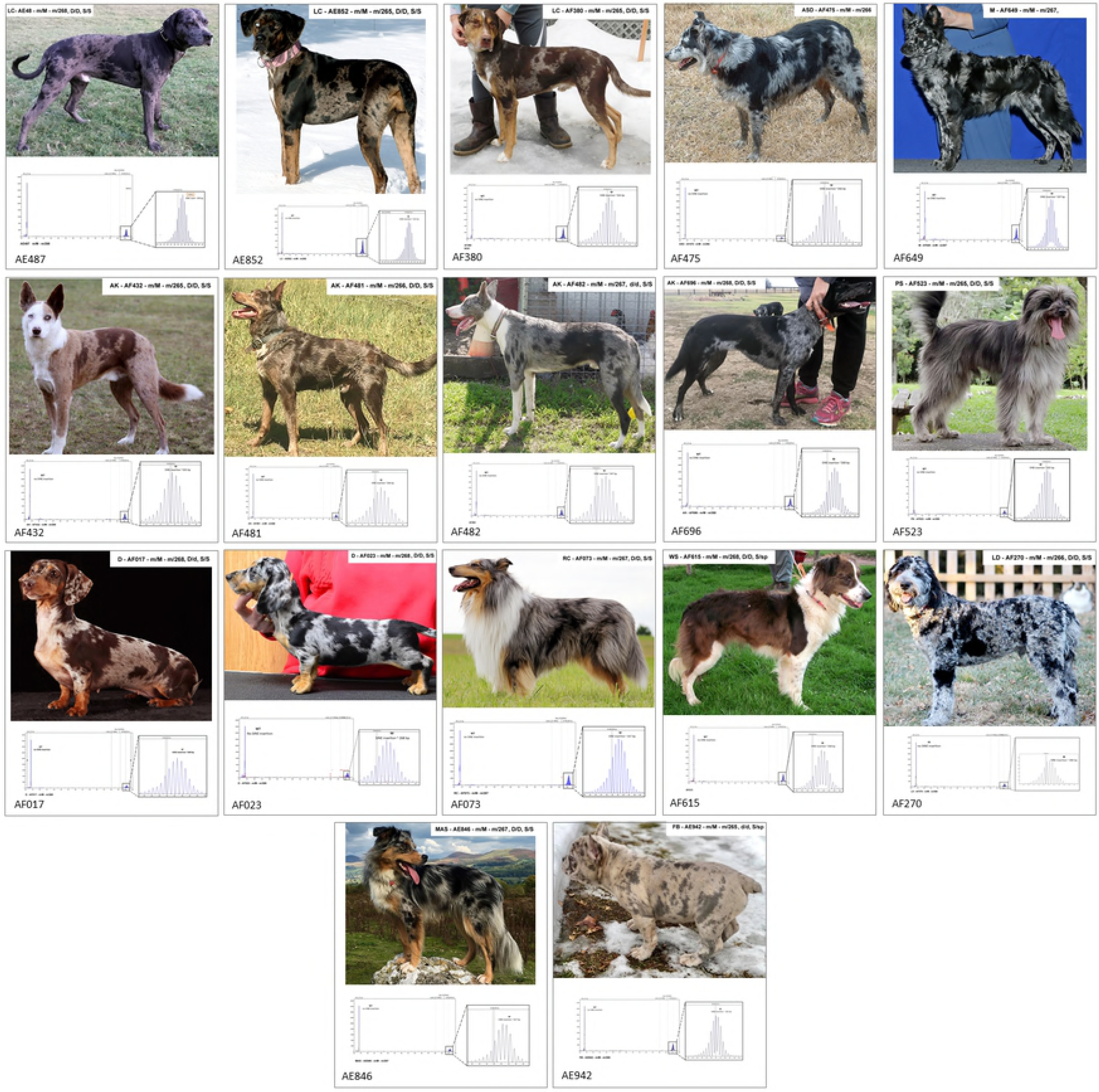

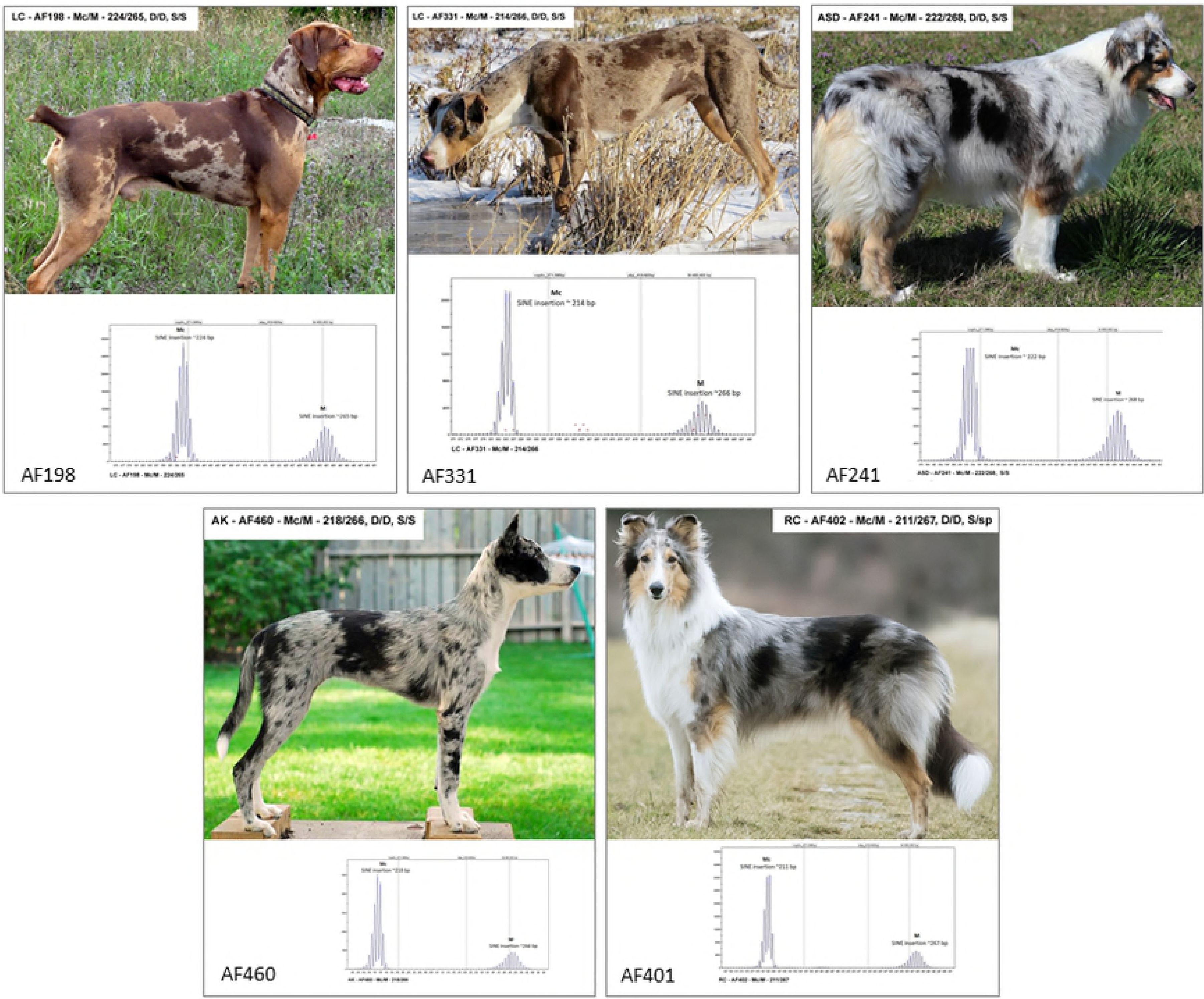

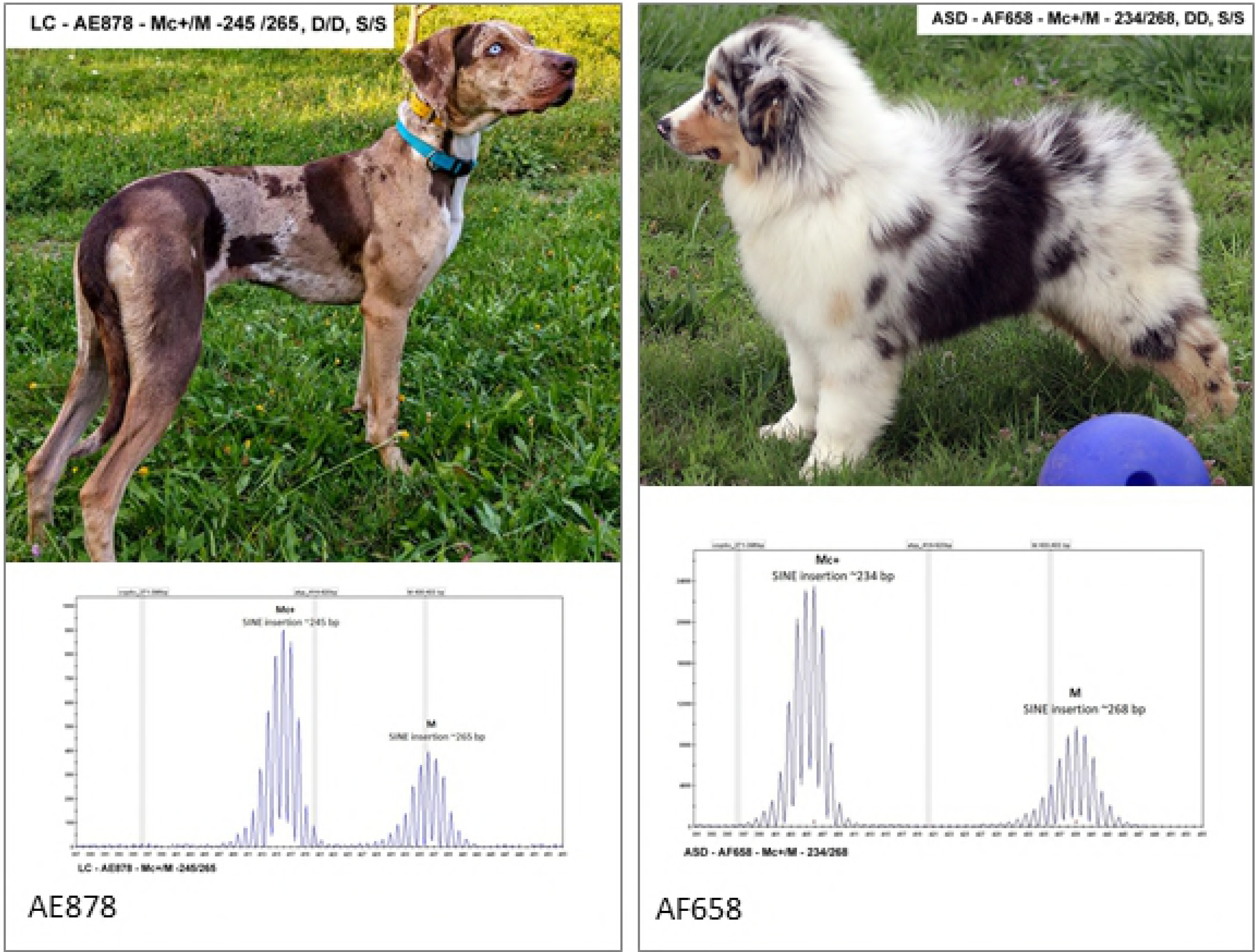

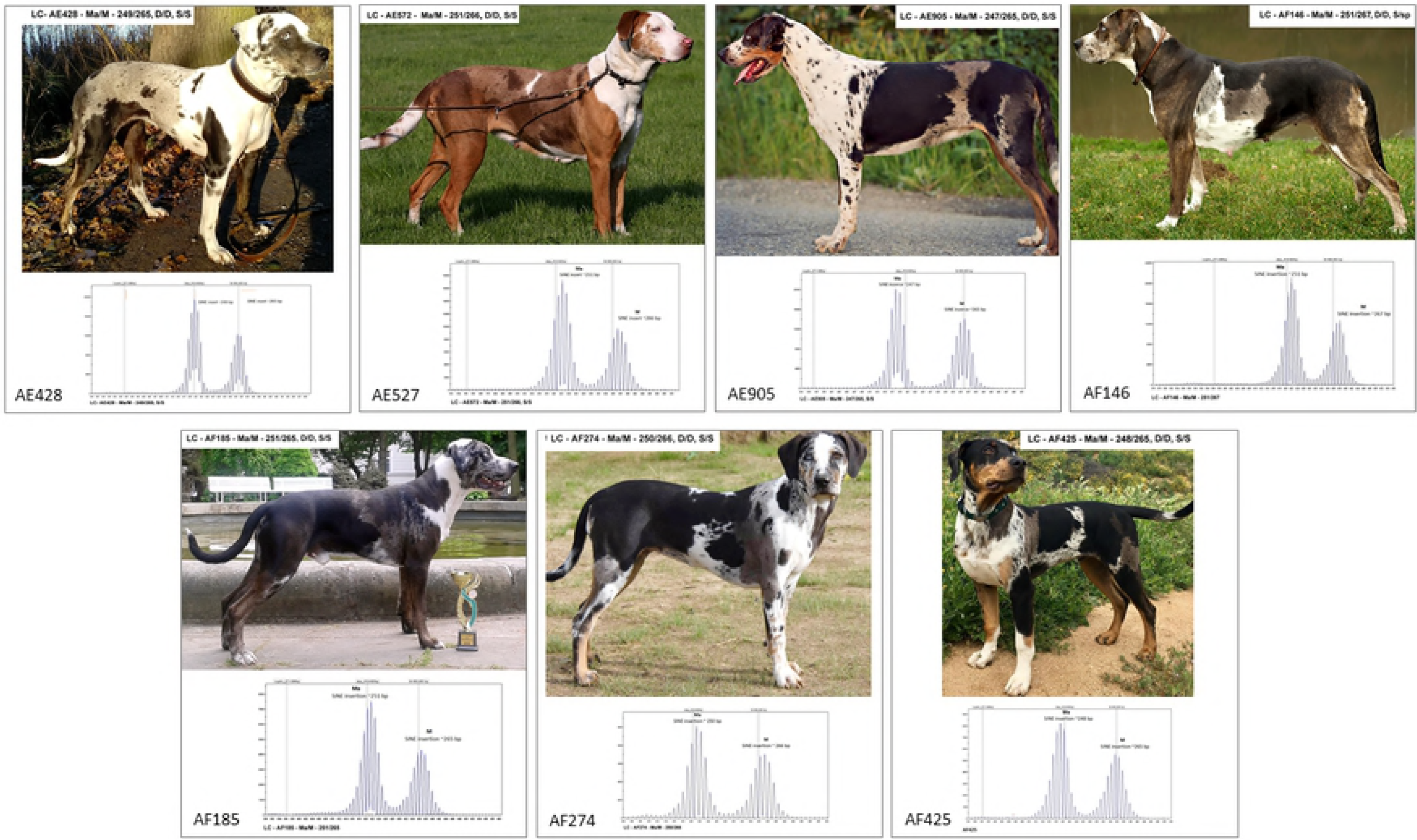

| SIRE | DAM | OFFSPRING | BREED |
| --- | --- | --- | --- |
| AF016 | AF017 | AF022, AF019, AF020, AF018 | Dachshund |
| AF014 | AF013 | AF008, AF009, AF010, AF011, AF012, AF015, AF237 | Australian Shepherd |
| AF014 | N/A | AF239, AF241 | Australian Shepherd |
| AE939 | AE869 | AE803, AE903 | Australian Shepherd |
| N/A | N/A | AE868, AE869 | Australian Shepherd |
| N/A | AE877 | AF050 | Australian Shepherd |
| AF761 | AF762 | AF758, AF759, AF760 | Australian Shepherd |
| N/A | N/A | AE572, AF146 | Catahoula |
| AE574 | AF146 | AF161 | Catahoula |
| AE515 | AF515 | AF612 | Catahoula |
| N/A | N/A | AE428, AE573, AF341 | Catahoula |
| N/A | AE904 | AE512 | Catahoula |
| N/A | N/A | AF699, AF352 | Catahoula |
| AF132 | AF053 | AE878, AE804, AF399 | Catahoula |
| AE505 | AF402 | AE504, AF231 | Catahoula |
| AE505 | N/A | AF101 | Catahoula |
| AF099 | AE622 | AF232 | Catahoula |
| AF129 | AE787 | AE740, AE515 | Catahoula |
| AF133 | N/A | AF176 | Catahoula |
| N/A | N/A | AE463, AE535 | Catahoula |
| AE956 | N/A | AF185 | Catahoula |
| AE505 | AE852 | AF196 | Catahoula |
| AF380 | AF038 | AF379, AF378 | Catahoula |
| AF330 | AF510 | AF333, AF335 | Catahoula |
| AF333 | AF510 | AF334 | Catahoula |
| AF333 | AF331 | AF332 | Catahoula |
| AF513 | N/A | AF400 | Australian Koolie |
| AF508 | AF484 | AF327, AF521, AF485 | Australian Koolie |
| N/A | AE819 | AE818 | Border Collie |
| AF056 | N/A | AF514 | Border Collie |
| N/A | N/A | AF649, AF650 | Mudi |

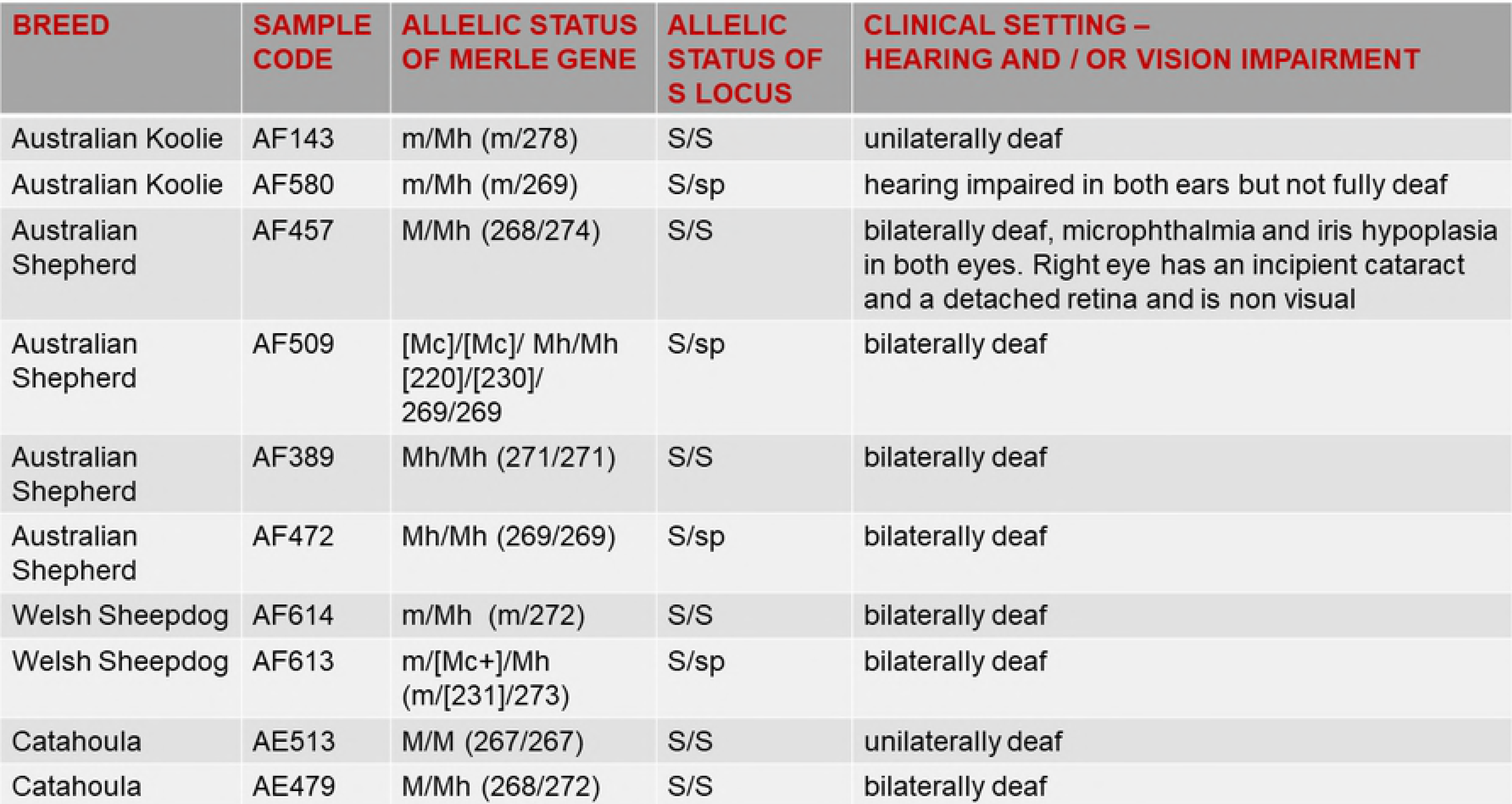

